# Evaluating power to detect recurrent selective sweeps under increasingly realistic evolutionary null models

**DOI:** 10.1101/2023.06.15.545166

**Authors:** Vivak Soni, Parul Johri, Jeffrey D. Jensen

## Abstract

The detection of selective sweeps from population genomic data often relies on the premise that the beneficial mutations in question have fixed very near the sampling time. As it has been previously shown that the power to detect a selective sweep is strongly dependent on the time since fixation as well as the strength of selection, it is naturally the case that strong, recent sweeps leave the strongest signatures. However, the biological reality is that beneficial mutations enter populations at a rate, one that partially determines the mean wait time between sweep events and hence their age distribution. An important question thus remains about the power to detect recurrent selective sweeps when they are modelled by a realistic mutation rate and as part of a realistic distribution of fitness effects (DFE), as opposed to a single, recent, isolated event on a purely neutral background as is more commonly modelled. Here we use forward-in-time simulations to study the performance of commonly used sweep statistics, within the context of more realistic evolutionary baseline models incorporating purifying and background selection, population size change, and mutation and recombination rate heterogeneity. Results demonstrate the important interplay of these processes, necessitating caution when interpreting selection scans; specifically, false positive rates are in excess of true positive across much of the evaluated parameter space, and selective sweeps are often undetectable unless the strength of selection is exceptionally strong.

**Teaser Text:** Outlier-based genomic scans have proven a popular approach for identifying loci that have potentially experienced recent positive selection. However, it has previously been shown that an evolutionarily appropriate baseline model that incorporates non-equilibrium population histories, purifying and background selection, and variation in mutation and recombination rates is necessary to reduce often extreme false positive rates when performing genomic scans. Here we evaluate the power to detect recurrent selective sweeps using common SFS-based and haplotype-based methods under these increasingly realistic models. We find that while these appropriate evolutionary baselines are essential to reduce false positive rates, the power to accurately detect recurrent selective sweeps is generally low across much of the biologically relevant parameter space.

## Introduction

In 1974, Maynard Smith and Haigh demonstrated that when a positively selected mutation increases in frequency within a population, linked variation may increase in frequency along with it. In the case of a beneficial fixation, the resulting selective sweep is expected to temporally reduce local nucleotide diversity owing to the fixation of these linked variants (Berry et al. 1991; and interestingly, deleterious fixations may generate the same effect as well (Maruyama and Kimura 1974; Johri et al. 2021a)). If the strength of selection favouring the beneficial allele is strong it will be expected to reach fixation much faster than under genetic drift, resulting in a local distortion of underlying genealogies (see review of Charlesworth and Jensen 2021). The size of this swept region is dependent on not only the strength of positive selection (*i.e.,* relating to the time to fixation), but also on the rate of recombination given that crossover events may break-up associations between the selected allele and linked variation. The genetic hitchhiking effects associated with selective sweeps have been relatively well-described theoretically, and have been observed empirically as well (see review of Stephan 2019).

However, this classic selective sweep model studies the effect of a single beneficial fixation on surrounding neutral genetic variation in a purely deterministic fashion. Kaplan et al. (1989) extended this model to include the stochastic effects of genetic drift that are particularly important when the beneficial mutation has newly arisen and is vulnerable to stochastic loss. Furthermore, although the single sweep model described remains commonly used, the more realistic scenario is that of recurrent selective sweeps, in which beneficial mutations are modelled as occurring at a rate, as of course is the biological reality (Jensen 2009). In this regard, several studies have modelled beneficial mutations as occurring randomly across a chromosome according to a time-homogenous Poisson process at a per-generation rate (Kaplan et al. 1989; Wiehe and Stephan, 1993; Stephan 1995; Pavlidis et al. 2010).

Besides the reduction in nucleotide diversity surrounding the beneficial fixation, another signature of hitchhiking commonly employed to detect selective sweeps, and related to the underlying distortion of coalescent histories, is a shift in the site frequency spectrum (SFS). Under a single sweep model with recombination, a skew is expected in the direction of increasing both high and low frequency derived alleles within the vicinity of a beneficial mutation (Braverman et al. 1995; Simonsen et al. 1995; Fay and Wu, 2000). The theoretical basis of this effect has been described under the model of a single, recent selective sweep (*e.g.,* Kim and Stephan, 2002; Kim and Nielsen, 2004). Based on this expectation, Kim and Stephan (2002) developed a composite likelihood ratio (CLR) test that detects local reductions in nucleotide diversity and SFS skewing along a chromosome, using this signature to identify the selected locus as well as to estimate the strength of selection acting on this locus. Briefly, the test compares the probability of the observed SFS under the standard neutral model with the probability under the model of a selective sweep. However, because the null model for the CLR test is standard neutrality, violations of the model, such as population size change, may reduce power and inflate false positive rates. For example, Jensen et al. (2005) demonstrated that the CLR test is not robust to strong population bottlenecks, with a false positive rate approaching 80% under these scenarios. They extended the initial model to incorporate a goodness-of-fit test to directly evaluate the fit of the sweep model to the data, greatly improving performance. Relatedly, Nielsen et al. (2005) modified the CLR method for application to genome-wide data. Unlike the Kim and Stephan (2002) approach, this method (termed SweepFinder, and the more recently released SweepFinder2 (DeGiorgio et al. 2016)) instead utilizes a null model directly derived from the empirical SFS, in an attempt to account for deviations from the standard neutral expectation in a model-free manner. Crisci et al. (2013) evaluated the power of SweepFinder, finding that although the method has low (*i.e*., improved) false-positive rates under numerous bottleneck models, the true-positive rates for identifying sweeps when the population experienced bottlenecks also tended to be under 10%. In other words, the fundamental difficulty of distinguishing selective sweeps from neutral population bottlenecks still very much remains (Barton 1998).

The specific details of the beneficial trajectory - whether positive selection began acting on the mutation while it was rare or common, whether a single or multiple beneficial mutations were involved, and whether the beneficial mutation has yet reached fixation - are all important considerations in determining expected patterns of variation. For example, if a sweep occurs from a common standing genetic variant (*i.e.,* if a neutral or deleterious allele segregating at relatively high frequency becomes positively selected upon a shift in selection pressure), the reduction in diversity will be dependent on the starting frequency of the beneficial allele. Furthermore, if that beneficial allele was segregating on multiple genetic backgrounds owing to recombination at the onset of selection, multiple haplotypes may increase in frequency and remain at intermediate frequency in the population at the conclusion of the selective sweep. This scenario has been termed as a soft selective sweep (as opposed to the classic model of a hard selective sweep, in which selection acts on a rare mutation; Hermisson and Pennings 2005). However, some have argued that these models are unlikely across much of the biological parameter space, owing for example to the necessary condition of a relatively high pre-selection frequency and a severe shift in selective effects (Jensen 2014). For example, it may be the case that mutations which strongly impact a phenotype such that they may become strongly beneficial are unlikely to be segregating neutrally prior to the shift in selection pressure. If these mutations are instead segregating as rare deleterious mutations prior to the shift, the outcome is again likely to be a hard selective sweep (Orr and Betancourt 2001).

As an alternative to the SFS-based SweepFinder2, Garud et al. (2015) developed a suite of methods that utilize these expected haplotype shifts under hard and soft selective sweeps to infer sweep location and strength. More specifically, the statistics seek to capture the increase in haplotype homozygosity (*i.e*., the probability of randomly selecting two identical haplotypes within a population (Sabeti et al. 2006)) observed under these sweep models. To facilitate detection of soft as well as hard selective sweeps, their H12 statistic combines the frequencies of the first and second most common haplotypes into a single frequency. The authors observed elevated (relative to standard neutral expectations) H12 levels genome-wide in the *Drosophila melanogaster* DGRP data (Mackay et al. 2012), and suggested that their top 50 outlier loci were likely soft selective sweeps. However, Harris et al. (2018) subsequently demonstrated that the H12 statistic is elevated under a number of neutral non-equilibrium demographic histories relevant for the population in question, and indeed that the results of Garud et al. (2015) can be explained without invoking positive selection. More specifically, although models of recurrent hard selective sweeps, recurrent soft selective sweeps, and neutral non-equilibrium demographic histories were all consistent with the data, the data were found to be insufficient to distinguish amongst these possibilities (and see Johri et al. 2022a,b).

### Important considerations when modelling selective sweeps

Although positive selection has been among the most extensively studied forms of selection, we know comparatively less about the frequency and effect size of beneficial mutations than we do about neutral or deleterious mutations. This is in no small part due to how infrequently beneficial mutations occur relative to these other classes, and the difficulty in accurately identifying their presence in polymorphism and divergence-based datasets (Bank et al. 2014). As already touched upon, population history can introduce confounding effects when attempting to detect selective sweeps. Considering that many commonly studied populations and species may have experienced recent and severe population bottlenecks (*e.g.,* non-African populations of *Drosophila melanogaster* (Li and Stephan, 2006) and humans (Gutenkunst et al. 2009; Gravel et al. 2011; Excoffier et al. 2013), as well as multiple clinically significant pathogens owing to the underlying dynamics of host infection (Jensen and Kowalik, 2020; Jensen 2020; Morales-Arce et al. 2021; Terbot et al. 2023; Howell et al. 2023)), the effects of demography pose a significant limitation to our current knowledge of positive selection. Indeed, from a coalescent perspective, a selective sweep results in the same approximately-star-shaped coalescent history as certain strong neutral population bottlenecks (Barton 1998; Thornton and Jensen 2007; Poh et al. 2014; Harris and Jensen 2020), thereby necessitating the use of an appropriate demographic null model in order to avoid extreme false positive rates (Thornton and Jensen, 2007; Johri et al. 2022c).

Multiple other factors must also be considered. As it has been shown that recurrent selective sweeps may reduce the level of standing variation (Kaplan et al. 1989; Wiehe and Stephan 1993; Gillespie 2000), and as recombination reduces the hitchhiking effect, a positive correlation between local recombination rate and sequence diversity is predicted under recurrent selective sweeps, a correlation first observed in *D. melanogaster* by Begun and Aquadro (1992), and subsequently in numerous other species (see reviews of Cutter and Payseur 2013; Charlesworth and Jensen 2021). However, the hitchhiking effect generated by the removal of deleterious mutations - known as background selection (BGS) - will also generate this same relationship between recombination rate and sequence diversity (Hudson and Kaplan 1995; Charlesworth et al. 1993; Charlesworth 1996). Given that the deleterious mutation rate far exceeds the beneficial mutation rate, BGS is indeed the most parsimonious explanation (Elyashiv et al 2016; Campos and Charlesworth 2019), and thus - as with demography - BGS must be accounted for when performing scans for selective sweeps.

The picture is further complicated by the fact that demographic inference can be biased if selection and recombination-associated biased gene conversion (BGC) are not accounted for (Ewing and Jensen 2016; Johri et al. 2021b), whilst inferring selection parameters and recombination rates are likely to be biased when demographic inference is not taken into consideration, a problem that is ominously circular. Recent methods (*e.g.,* Johri et al. 2020; see also Johri et al. 2022c review) have sought to avoid this circularity, by jointly inferring parameters of both demography and the neutral and deleterious distribution of fitness effects (DFE). This is an important step in constructing a reliable baseline model. Johri et al. (2022a) have further highlighted the many factors that must be considered when constructing such a null model, with sweep inference being dependent on the inference of demography, the DFE, mutation and recombination rates.

With regards to the rate of beneficial mutations, previous studies have often assumed that at most one beneficial mutation is on the way to fixation at a given time (Kaplan et al. 1989; Stephan et al. 1992; Wiehe and Stephan 1993). However, interference between linked beneficial alleles may cause a reduction of their fixation probabilities (Hill and Robertson 1966; Birky and Walsh 1988; Kim and Stephan 2002). Whether interference will occur is dependent on the rate at which beneficial mutations arise and sweep to fixation within a population, as well as on the underlying strength of selection and recombination rate (Braverman et al. 1995; Przeworski 2002). Thus, for genome scans to work well, selection should be rare enough that there is no interference between beneficial mutations, but not so rare that sweeps are too old to detect on average (Teshima et al. 2006; Thornton and Jensen 2007). Relatedly, Jensen (2009) found that published genome scans identified an order of magnitude more sweeps than would be expected under published recurrent hitchhiking estimates, potentially due to a high false positive rate. This again highlights the important question of determining how much statistical power population genomic data provides to accurately estimate recurrent selective sweep parameters within the context of realistic evolutionary null models.

As discussed above, there has been considerable previous work discussing biases introduced by neglecting these many contributing evolutionary processes. Here we take a different approach. Namely, we investigate the power to detect recurrent selective sweeps under a scenario in which a researcher has performed their due diligence in accurately constructing a baseline model consisting of population history, purifying and background selection effects, and mutation and recombination rate variation. We ran simulations to assess the effectiveness of a common SFS-based approach (the SweepFinder framework; Nielsen et al. 2005)) and a haplotype-based approach (H12; Garud et al. 2015) under a variety of models incorporating these factors. We find that although a consideration of these factors indeed greatly reduces false-positive rates, the power to accurately detect selective sweep effects is generally severely limited as well. These results highlight that even under best-case scenarios, sweep scans require careful interpretation and scrutiny.

## Results and Discussion

### Single sweep model

We ran simulations in SLiM3 (Haller and Messer 2019) and quantified power to detect selective sweeps using the frequency spectrum-based composite likelihood method SweepFinder2 (DeGiorgio et al. 2016), as well as the haplotype-based statistic H12 for comparison (Garud et al. 2015).

Parameterizations for simulations were taken from the *D. melanogaster* literature (see Methods section for details). In order to first define power to detect selective sweeps under a best-case scenario, we simulated under a model in which a single sweep occurs in a population under equilibrium, and sampling takes place immediately after the completion of the sweep (*i.e*., *τ* = 0, where *τ* is the number of generations since the beneficial fixation scaled by 4*N*). The decay in diversity around a selective sweep has been shown to be a function of *τ*, the population scaled strength of selection (*α* = *2N_e_s*), and the rate of recombination (Maynard Smith and Haigh 1974; Przeworski 2002, 2003; Kim and Stephan 2000, 2002). Using equation 13 from Kim and Stephan (2000), Figure 1 shows the expected diversity around a selective sweep for different values of *τ* and *2N_e_s* with a fixed recombination rate, using the parameterizations from our simulations, with lower values of *τ* yielding a greater dip in diversity around the selective sweep. Thus, as has been well shown previously, we would expect to have the greatest power to detect a sweep immediately after beneficial fixation.

**Figure 1:**
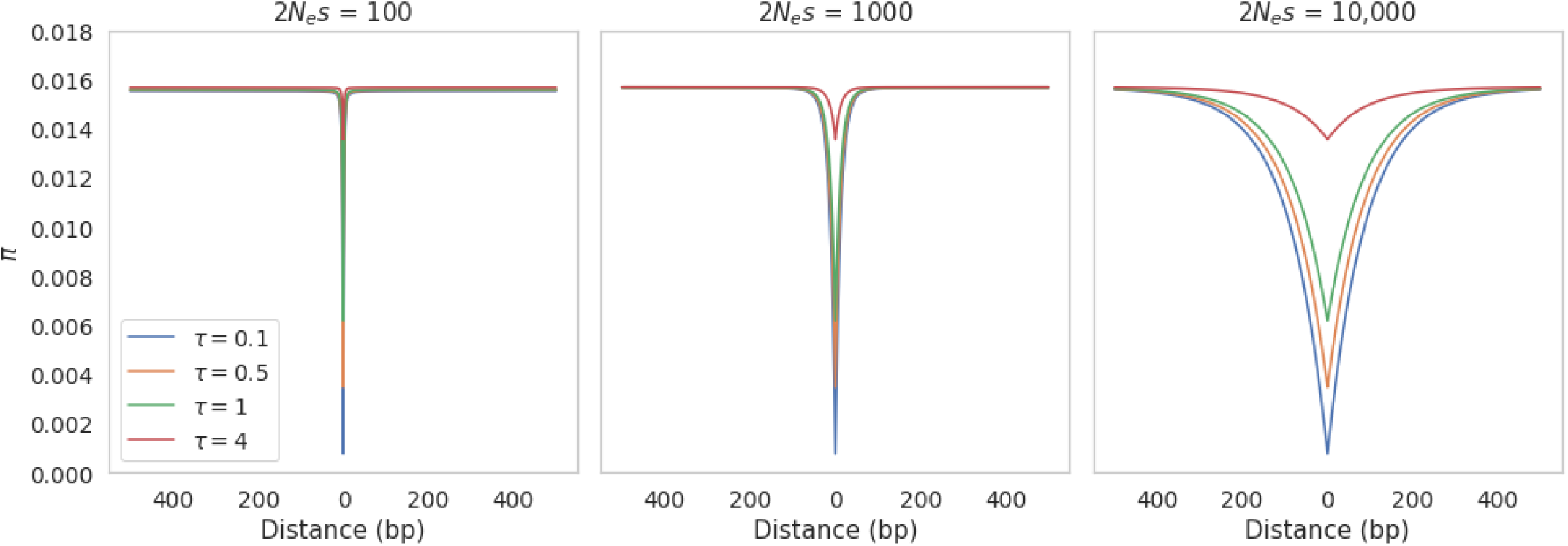
Nucleotide diversity over a physical distance for different values of 2*N_e_s* and *τ*, estimated using Equation 13 from Kim and Stephan (2000).

In order to relax the unrealistic but common assumption of exclusively neutral mutations occurring on the background of the beneficial variant, we next simulated a constant size population with functional regions consisting of 4 exons and 3 introns, separated by intergenic regions, with functional mutations drawn from a discrete DFE previously inferred from *D. melanogaster* (*i.e*., thereby incorporating both purifying and background selection effects; Johri et al. 2020; 2023). Sweep inference was performed using SweepFinder2 and the H12 statistic. For SweepFinder2, inference was performed at each single nucleotide polymorphism (SNP), while H12 inference was performed across 500 basepairs on either side of each SNP in order to quantify haplotype structure. Figure 2 presents results of sweep inference under this model, with a single beneficial mutation for a range of *2N_e_s* values [100, 1000, 10,000], where *N_e_ = N_ancestral_*, and with neutral and deleterious mutations drawn from the full DFE. Here, sampling occurred directly after the completion of the sweep, as is commonly assumed. The threshold for sweep detection was determined by the highest CLR and H12 values across a set of 200 simulations in which all else was modelled identically, except that no beneficial mutations were occurring (see Methods). This is a best-case-scenario threshold that should produce no false positives.

**Figure 2:**
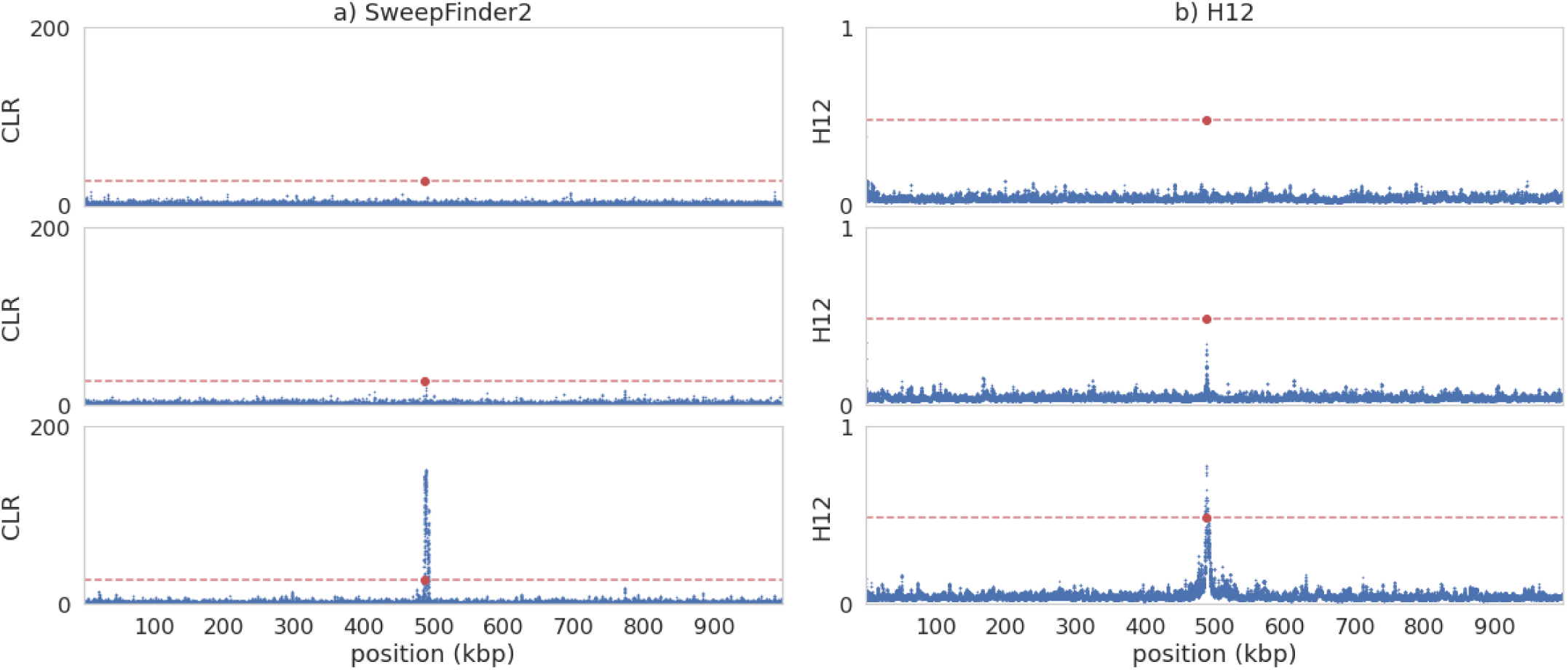
Example patterns around a single selective sweep for different values of 2*N_e_s* in an equilibrium population with fixed mutation and recombination rates. In each case, 2*N_e_s* values go from lowest (top panel) to highest: 100; 1000; 10,000. The red data point is the position of the beneficial fixation. a) Inference results from SweepFinder2. Blue data points are CLR values inferred for each window. The red dashed line is the threshold for sweep detection, determined by the highest CLR value across 200 simulated replicates in which no beneficial mutations are occurring. Inference was performed at each SNP (see Methods section for further details). b) Sweep inference with the H12 statistic. Blue data points are H12 values estimated for each window. As with SweepFinder2, the red dashed line is the threshold for sweep detection. Inference was performed across 1kb windows for each SNP, with the SNP at the center of each window. For the underlying summary statistics (Tajima’s *D*; *π*; and *r^2^*), see supplementary figure, S1.

As shown, essentially no power was observed to detect sweeps at *2N_e_s* values of 100 or 1000 with either SweepFinder2 or H12, suggesting that even in an optimal scenario the strength of positive selection needs be extremely strong for reliable detection. This is consistent with previous results (Jensen et al. 2007; Crisci et al. 2013). As Figure 1 demonstrates, at these *α* values diversity recovers over extremely small distances around the swept locus for the recombination rate considered, and thus the signature quickly dissipates (Supplementary Figure S1; and see Supplementary Figures 2 & 3 for the null thresholds and related summary statistics).

### Recurrent sweep equilibrium model

In contrast to the single sweep model used above, recurrent selective sweep models consider a scenario in which sweeps occur randomly across a chromosome according to a time-homogenous Poisson process at a per-generation rate (Kaplan et al. 1989; Wiehe and Stephan 1993; Stephan 1995; Pavlidis et al. 2010). This model represents a considerably more realistic scenario in which beneficial mutations are simply modelled as occurring at a mutation rate. It has previously been shown that power to detect recurrent sweeps generally increases with the rate of selective events, as would be expected (Jensen et al. 2007; Pavlidis et al. 2010). Generally speaking, if sweeps are rare but strong, they may affect a large proportion of the genomic region, but on average may be too old to detect using patterns of polymorphism; if they are common and weak then the size of the genomic region affected may be too small to be detected.

Here we modelled beneficial mutations as part of a full DFE, requiring definition of the proportion of new mutations that are beneficial (*i.e*., the beneficial mutation rate). Although this proportion is difficult to accurately infer, it has been well-observed that non-coding divergence is higher than coding divergence in *D. melanogaster* (Andolfatto 2005), setting some upper limit on the possible fraction of beneficial fixations in coding regions. We estimated coding and non-coding divergence per gene for different proportions of beneficial mutations [5%, 0.5% and 0.05%] (see Supplementary Figure S4). At 5% and 0.5%, coding divergence was either higher than or equal to non-coding divergence at the highest *2N_e_s* value. As such we chose to proceed with 0.05% of coding mutations being beneficial, giving a beneficial point mutation rate of 1.4e-12. Supplementary Table 1 provides the proportion of fixations that are beneficial under this scenario. Furthermore, although our model assumed a continuous supply of beneficial mutations, it has been shown that there is very little power to detect older sweeps (Przeworski 2002; Kim and Stephan 2002). We therefore only included fixations that occurred up to 0.5*N* generations ago when attempting sweep inference.

Figure 3 presents summary statistics and the results of sweep inference for an equilibrium demographic model; namely, nucleotide diversity, *r*^2^ (a measure of linkage disequilibrium), and a summary of the site frequency spectrum, Tajima’s *D* (Tajima 1989). As before, we implemented the DFE from Johri *et al*. (2020), though with the introduction of beneficial mutations, the proportion of effectively neutral mutations were correspondingly reduced to account for the addition. As with the single sweep model, there was little power to detect selective sweeps at *2N_e_s* = 100; indeed, very few beneficial mutations fixed 0.5*N* generations prior to sampling across all simulated replicates. With increasing *2N_e_s* the sweeps leave a greater genomic signature as expected.

**Figure 3:**
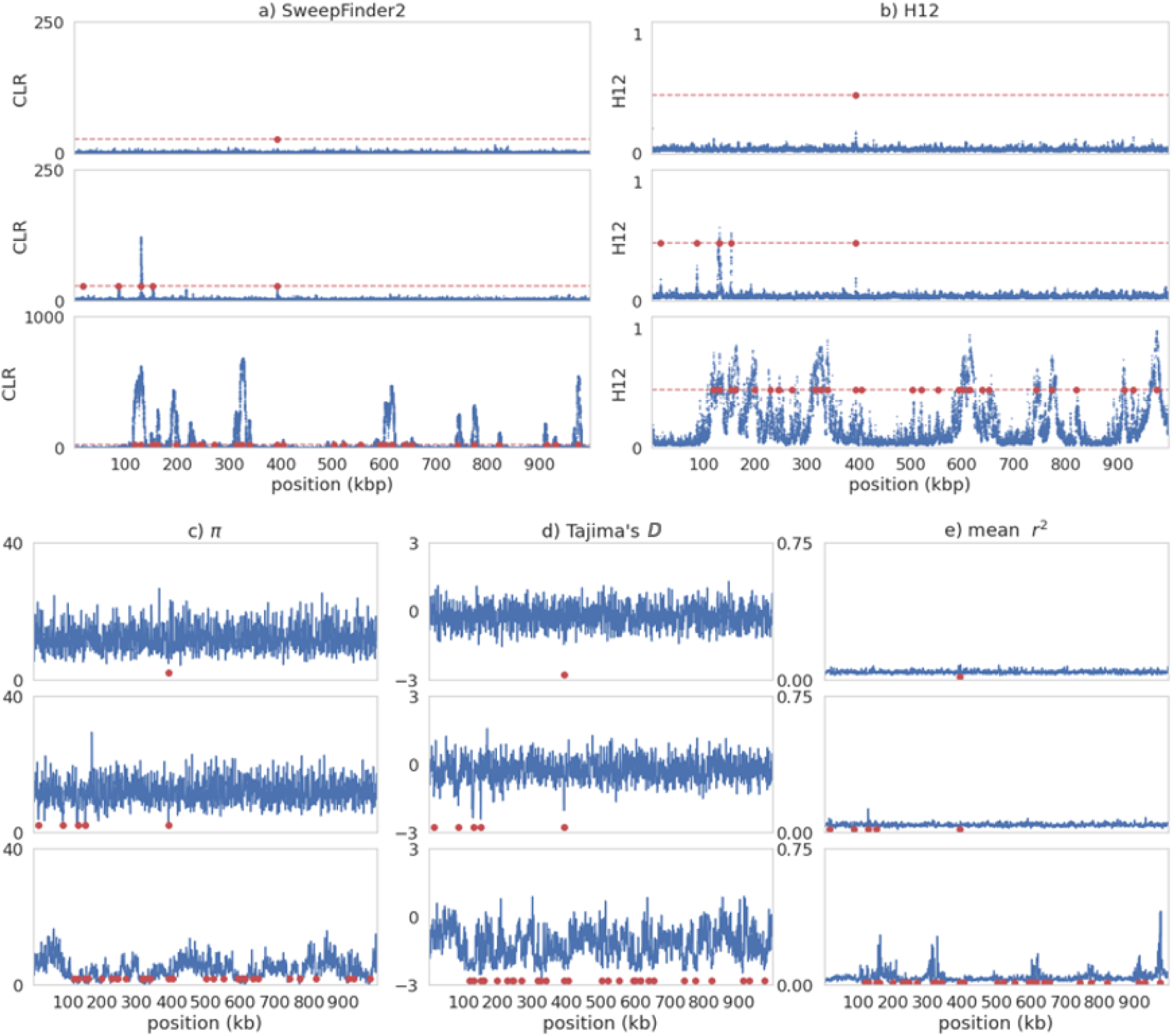
Sweep inference and summary statistics for a single simulation replicate of recurrent selective sweeps for different values of 2*N_e_s* in an equilibrium population with fixed mutation and recombination rates. In each case, 2*N_e_s* values go from lowest (top panel) to highest: 100; 1000; 10,000. For all panels, red data points are the positions of beneficial fixations within the previous 0.5*N* generations prior to sampling. a) Inference results from SweepFinder2. Blue data points are CLR values inferred for each window. The red dashed line is the threshold for sweep detection, determined by the highest CLR value across 200 simulated replicates in which no beneficial mutations are modelled. Inference was performed at each SNP (see Methods section for further details). b) Sweep inference with the H12 statistic. Blue data points are H12 values estimated for each window. As with SweepFinder2, the red dashed line is the threshold for sweep detection. Inference was performed across 1kb windows for each SNP, with the SNP at the center of each window. c-e) Summary statistics across the simulated region.

To better visualise the power of both methods, true positive rates (TPR) and false positive rates (FPR) were calculated across 10kb non-overlapping windows, and receiver operating characteristic (ROC) curves across 200 simulated replicates were plotted (Figure 4). A true positive (TP) was defined as a window containing a SNP that has met the inference threshold and was within 500bp of a beneficial mutation that had fixed within 0.5*N* generations of sampling. One potential issue with this definition is that if the signature of the sweep extends beyond the window in which it is located, adjacent hitchhiked windows will be classified as false positives (FP) if they are not within 500bp of the beneficial mutation. At high strengths of selection this could result in a chain of FPs which are all the result of a true sweep signature. To address this issue, we defined any FP window that met the inference threshold and was adjacent to a TP window as a TP. This procedure could continue indefinitely, until a window which failed to meet the inference threshold was encountered, thereby accounting for the size of the sweep signature. Supplementary Figure S5 provides the comparison between the standard and “adjacent windows” methods, with a notable increase in power using the latter approach, which is the approach utilized throughout this study.

**Figure 4:**
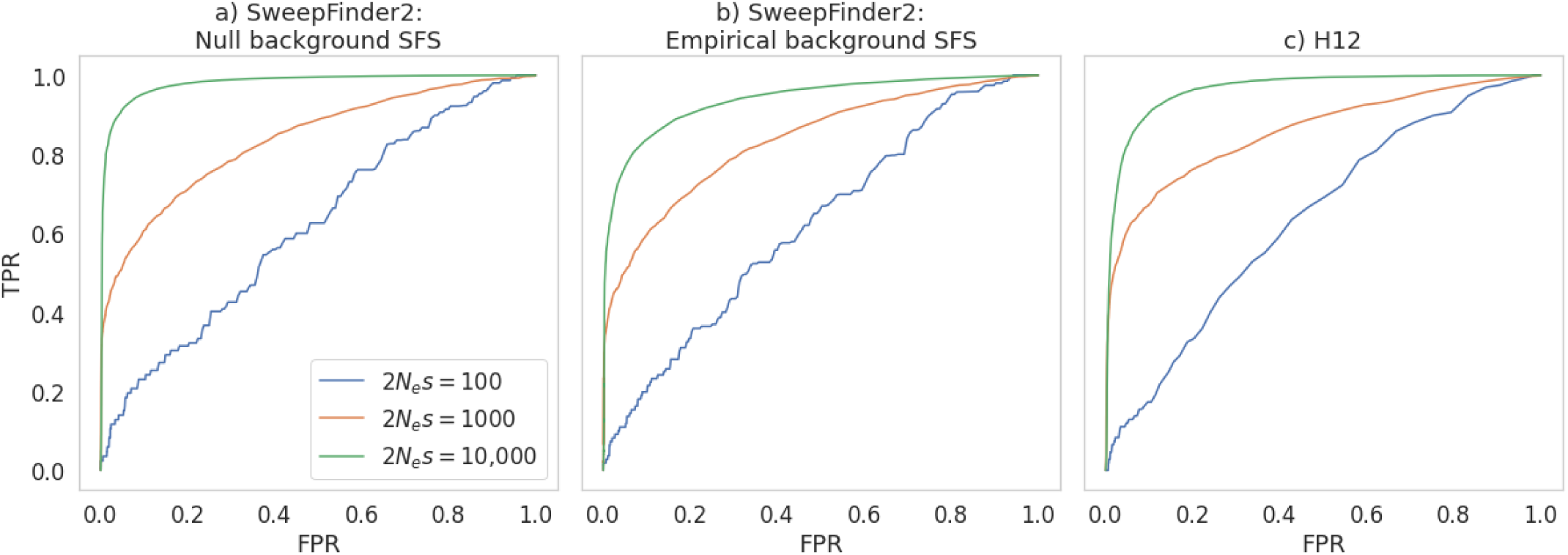
ROC curves, showing the change in true positive rate (TPR) as the false positive rate (FPR) increases, for sweep inference in an equilibrium population with fixed mutation and recombination rates across 200 simulated replicates, for 10kb windows. a) ROC curves for SweepFinder2 when using a null background SFS (*i.e.,* the background SFS is generated across a simulation run in which all else is modelled identically, except that no beneficial mutations occur). b) ROC curves for SweepFinder2 when using an empirical background SFS (*i.e.,* the background SFS is the empirical data itself). c) ROC curves for H12.

It is notable that nearly the entire genomic region will be affected by recurrent sweeps at larger *2N_e_s* values, violating the assumption of a sweep-free background SFS (Pavlidis et al. 2010). In other words, sweeps become undetectable if the background comparison is swept as well. This can be seen in the reduction in power when using the empirical background SFS (Figure 4b) and the null background SFS (Figure 4a). Consistent with this logic, the most notable reduction in power is at *2N_e_s* = 10,000.

At 10kb window sizes, both SweepFinder2 and H12 had considerable power to detect recurrent selective sweeps at *2N_e_s* = 10,000, though this power decreased with *2N_e_s* (Figure 4). At lower *2N_e_s* values power to detect sweeps remained low. We repeated the analysis using a 1kb window size (Supplementary Figure S6), finding that inference power was greatly improved with both methods at *2N_e_s* = 1000, and somewhat reduced at *2N_e_s* = 10,000. These results highlight the fact that optimal window-sizes are a factor of the strength of selection and the recombination rate, and here we see that the optimal window-size appears to be roughly the size of the sweep. In natural populations one often has limited information on the recombination rate, and no information on the strength of selection (and therefore on the size of the sweep), making window size choice somewhat arbitrary. This can be problematic given that – as we show here – the size of the window is an important consideration that affects inference power. One approach suggested by Pavlidis et al. (2010) is to use a variable window size approach, where the window size adopted in any region of the genome is that which optimises the strength of the observation.

Thus, under this basic model that makes a number of simplifying assumptions (*i.e*., equilibrium populations with fixed recombination and mutation rates), there is considerable power to accurately detect recurrent selective sweeps if the strength of selection is great enough to leave a detectable signature (which will be dependent on window size, as discussed above). In order to consider increasingly realistic evolutionary baseline models, we next explored the effects of non-equilibrium demography on recurrent sweep inference, as well as heterogenous recombination and mutation rates.

### Recurrent sweep inference under non-equilibrium population histories

The confounding effects of demography on sweep inference are well documented (Barton 2000; Kim and Nielsen 2004; Jensen et al. 2005; Nielsen et al. 2005; Jensen et al. 2007; Pavlidis et al. 2008; Pavlidis et al. 2010; Poh et al. 2014; see also reviews of Pavlidis and Alachiotis 2017 and Stephan 2019). We simulated several instantaneous population size changes, where *N_current_* = [2, 0.5, 0.1] *N_ancestral_*. In each case the size change occurred *N* generations prior to sampling, where *N = N_ancestral_*. Figure 5 presents ROC curves for windows of size 10kb (see Supplementary Figure S7 for ROC curves corresponding to windows of size 1kb). Supplementary Figures S8-10 provide inference results and summary statistics for an example replicate from each of the population expansion, 50% contraction, and 90% contraction models, respectively. For example, and as expected, there was a small increase in variation and frequency spectrum skew, and a small reduction in *r^2^,* with the population expansion (Supplementary Figure S8), while population contractions increased levels of linkage disequilibrium (Supplementary Figures S9-10). The thresholds for sweep detection (set by the “null” simulations) followed a clear pattern with the H12 statistic (Table 1), with the long haplotypes generated by a population contraction resulting in higher H12 values. A 90% population contraction generated the maximum H12 value of 1, rendering sweep detection under this framework highly fraught. As these population contractions will generate neutral sweep-like coalescent events across the genome, the power to detect sweep-related multiple merger events is greatly reduced owing to the use of the background SFS as a null in SweepFinder2 under these models (*e.g.,* Barton 1998; Harris and Jensen 2020).

**Figure 5:**
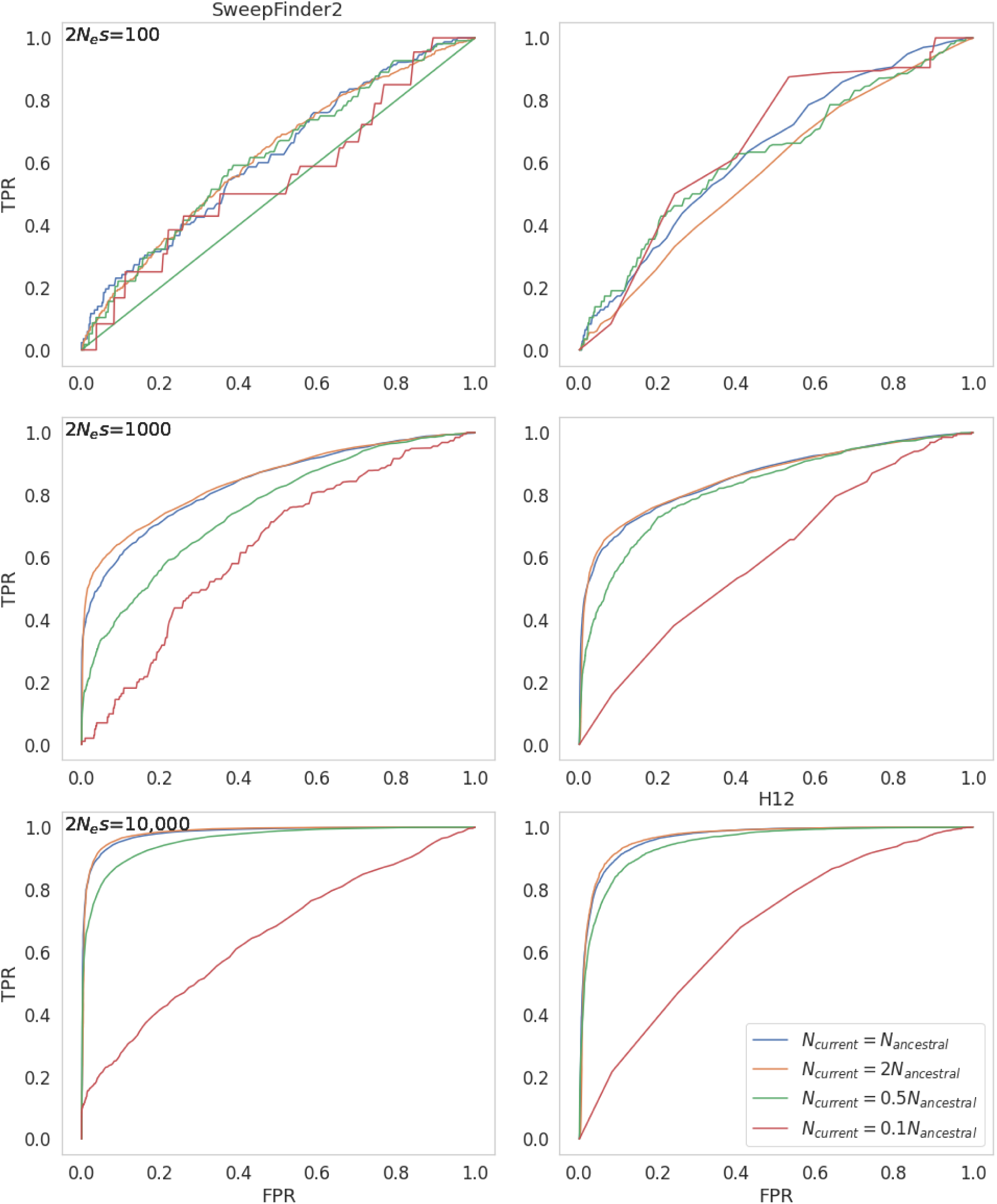
ROC curves, showing the change in true positive rate (TPR) as the false positive rate (FPR) increases, for sweep inference in populations with differing demographic histories, across 200 replicates each, for windows of size 10kb. The panels on the left are for inference with SweepFinder2, and on the right with the H12 statistic. Where population size change occurs, it is instantaneous, occurring *N* generations prior to sampling.

**Table 1:**
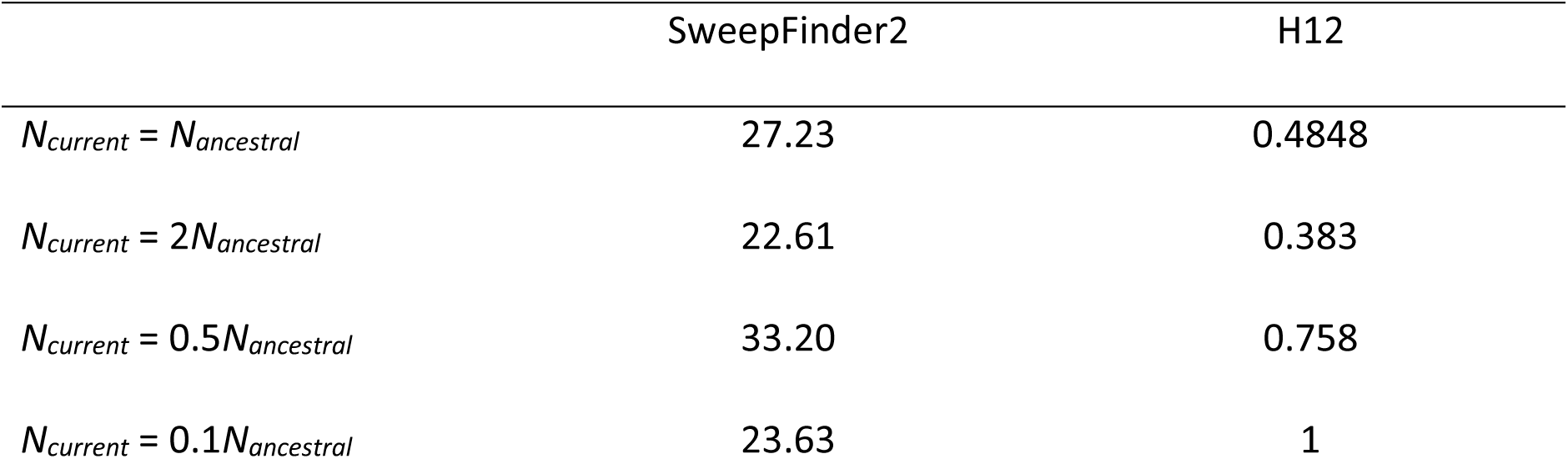
Threshold values for sweep detection for different demographic models, in which the threshold value is the highest CLR or H12 value across null simulations.

We directly compared the power of both methods under different demographic scenarios using ROC curves (Figure 5). In all cases power was relatively low for both methods, though there were some notable effects of demography. With H12 there was little perceptible change in power with the 0.5x contraction or the 2x expansion. SweepFinder2 showed similar patterns. With the 0.1x contraction the reduction in power was considerable owing to the increase of stochastic effects. Importantly, humans (Gutenkunst et al. 2009; Gravel et al. 2011) and *D. melanogaster* (David and Capy 1988; Lachaise et al. 1988; Baudry et al. 2004; Thornton and Andolfatto 2006) - two of the most widely studied organisms in evolutionary genomics - have undergone strong population bottlenecks in the process of migrating out of Africa; these species are also unlikely to be unique in this regard of having experienced major changes in population size.

### The effects of variable recombination and mutation rates on recurrent sweep inference

We next considered the effects of the common assumption of constant mutation and recombination rates. In reality, both the rate of recombination (Kong et al. 2002; Cox et al. 2009; Rockman and Kruglyak 2009; Comeron et al. 2012; Kawakami et al. 2014; and see the review of Stapley et al. 2017) and the rate of mutation (Lynch 2010; Hodgkinson and Eyre-Walker 2011; Rahbari et al. 2016; Carlson et al. 2018; Pfeifer 2020; and see the review of Baer et al. 2007) have been shown to be heterogenous across the genomes of numerous taxa. To assess the impact of variable mutation and recombination rates, we considered simulated data in which each 10kb region has a rate drawn from a uniform distribution, such that each simulated variable rate replicate has the same mean rate as the fixed rate comparisons (See Methods for further details). We examined three scenarios: fixed recombination rate / variable mutation rate; variable recombination rate / fixed mutation rate; variable recombination rate / variable mutation rate; and compared these to the fixed rate scenarios presented in the above sections. Figure 6 presents ROC curves for SweepFinder2 and H12 inference for equilibrium population demography across all four scenarios.

**Figure 6:**
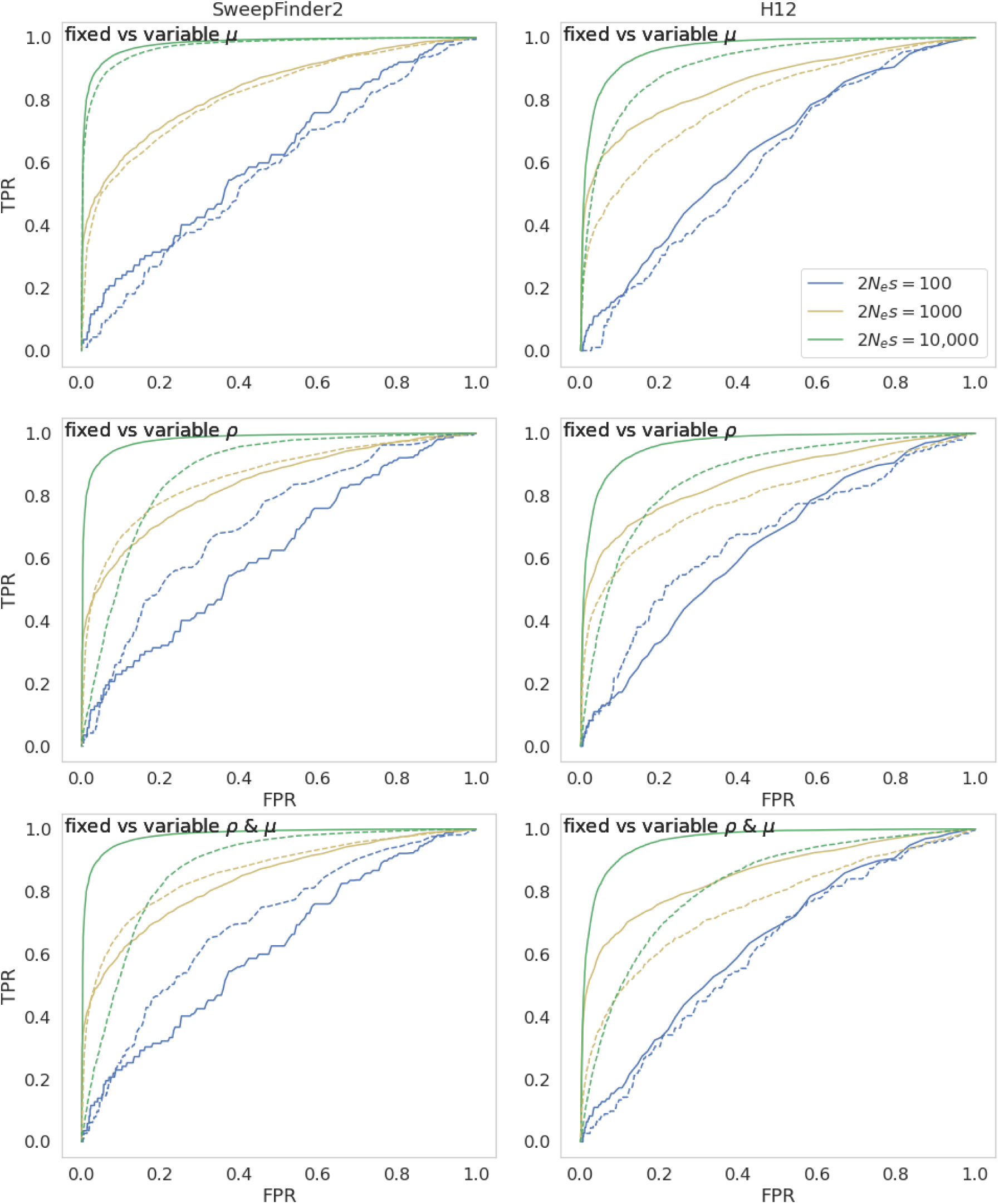
ROC curves comparing sweep inference for fixed and variable recombination and mutation rates under equilibrium demographic conditions, across 200 simulation replicates using SweepFinder2 using the null background SFS (left) and H12 (right), for 10kb windows. Dashed lines indicate variable rates, whilst filled lines indicate fixed rates. For variable rates, each 10kb region has a rate drawn from a distribution such that each simulated replicate has the same mean rate as the fixed rate comparison (See Methods for further details).

The general effect of variable mutation rates on both SweepFinder2 and H12 inference was a reduction in power (Figure 6; see Supplementary Figure S11 for 1kb inference windows). However, this reduction in power is mediated by both the strength of selection and the window size. For example, whilst the reduction in SweepFinder2 inference power is small at *2N_e_s* = 10,000 for both 10kb and 1kb window sizes, the reduction in power at *2N_e_s* = 1000 is small when using a window size of 10kb, but substantial at 1kb. To better quantify whether mutation rate is driving this reduction in power, results were divided into low and high mutation rate bins (Supplementary Figure S12), with a mean population-scaled mutation rate for the low rate bin of 2.03e-9 and 3.65e-9 for the high rate bin. As shown, the effect is modest.

Variable recombination rates also affected recurrent sweep inference power, with the magnitude of *2N_e_s* influencing whether there was a corresponding increase or decrease in power (Figure 6). Once again, by splitting simulated data into low and high recombination rate bins (Supplementary Figure S13), one may better interpret these results - with a mean sex-averaged recombination rate for the low rate bin of 0.09cM/Mb and 2.18cM/Mb for the high rate bin. As shown, inference power for both Sweepfinder2 and H12 was generally reduced in high recombination rate regions relative to low recombination rate regions, owing to the decreasing size of the resulting selective sweep. This pattern was evident across values of *2N_e_s.* When both mutation and recombination rates were variable, recombination rate variation appeared to be driving the change in power (Figure 6 and Supplementary Figure S11), although this will naturally depend on the range of the variation in both rates. Supplementary Figures S14-16 provide sweep inference results and summary statistics for an example replicate with variable mutation and recombination rates.

The interplay of nonequilibrium demography and heterogeneity in mutation and recombination rates was also examined. Supplementary Figures S17-19 present ROC plots for variable rate models combined with the non-equilibrium demographic models discussed in the sections above for 10kb windows; Supplementary Figures S20-22 present ROC plots for 1kb windows. Finally, Supplementary Figures S23-31 present sweep inference results and summary statistics for example replicates of these scenarios. As shown, there is a relatively complex interplay between these factors, with both population size change and variable rates being associated with losses in power, and the extent of these effects being dependent on both the strength of selection and the window size, with the effect of the latter being correlated with the former.

These complex dynamics highlight the necessity of utilizing high quality mutation and recombination rate maps when attempting to infer selective sweeps. For organisms in which these rates are not well-characterized, this marks the importance of firstly attempting to characterize these rates prior to further inference (see reviews of Lynch et al. 2016 and Pfeifer 2020 for common mutation rate inference approaches; Stumpf and McVean 2003 and Peñalba and Wolf 2020 for common recombination rate inference approaches). Additionally, as these underlying rates will always be associated with uncertainty, the range of feasible parameter values can be sampled in order to accurately quantify sweep-detection performance in light of this uncertainty (Johri et al. 2022c). We note that in the case of mutation rate variation, inter-species divergence information can be used to distinguish between the signal of a selective sweep and the effects of a reduction in polymorphism due to a reduced mutation rate. The efficacy of this approach will however depend on a mutation rate that is constant over relatively deep evolutionary time and on the presence of appropriate outgroup species.

### Comparing the power of an evolutionarily appropriate null model with outlier approaches

A common approach when searching for loci that may have experienced recent positive selection is the identification of genomic outliers (*e.g.,* Akey et al. 2002; Payseur et al. 2002; Harr et al. 2002; Glinka et al. 2003; Bauer DuMont and Aquadro 2005; Jensen et al. 2007; Garud et al. 2015; and see reviews of Thornton et al. 2007 and Akey et al. 2009). Briefly, this involves scanning across a large number of genomic regions, and identifying outlier regions that fall in some pre-determined tail of this observed empirical distribution. To compare the power of utilizing an evolutionarily appropriate null - which explicitly models these evolutionary processes - with the power of genomic scans for empirical outliers, we calculated: a) the fraction of windows that met the threshold (*i.e.,* the 5% tail for the outlier approach, compared to the maximum CLR and H12 values from our null model simulations) that contained a selective sweep (labelled as TP); b) the fraction of windows that met the threshold and did not contain a selective sweep (labelled as FP); and c) the fraction of windows that contained a selective sweep and did not meet the threshold value (labelled as FN). Tables 2 and 3 present the results of this analysis for H12 and SweepFinder2, respectively, for an equilibrium population with fixed rates. For non-equilibrium population and variable rate results see Supplementary Tables S2-9.

**Table 2:** Comparing the power of H12 using the evolutionarily appropriate null model, with an outlier approach that assumes windows in the 5% right-tail contain selective sweeps. Here TP is defined as the percentage of windows that have met the threshold value and contain a selective sweep; FP is defined as the percentage of windows that have met the threshold value and do not contain a selective sweep; FN is defined as the percentage of windows that contain a sweep and have not met the threshold value. Values are averaged across the 200 simulated replicates under population equilibrium with fixed recombination and mutation rates. Window size is 10kb.

| Population history | 2*N<sub>e</sub>s* of beneficial mutations | 95% tail |  |  | Null threshold |  |  |
| --- | --- | --- | --- | --- | --- | --- | --- |
|  |  | mean TP | mean FP | mean FN | mean TP | mean FP | mean FN |
| *N<sub>current</sub>* = *N<sub>ancestral</sub>* | No sweeps | 0.00 | 100.00 | 0.00 | - | - | 0.00 |
|  | 100 | 5.10 | 94.90 | 87.25 | 0.00 | 0.00 | 100.00 |
|  | 1000 | 55.81 | 44.19 | 50.02 | 84.01 | 15.99 | 77.52 |
|  | 10,000 | 96.28 | 3.72 | 84.75 | 97.27 | 2.73 | 46.44 |

**Table 3:** Comparing the power of SweepFinder2 using the evolutionarily appropriate null model, with an outlier approach that assumes windows in the 5% right-tail contain selective sweeps. Here TP is defined as the percentage of windows that have met the threshold value and contain a selective sweep; FP is defined as the percentage of windows that have met the threshold value and do not contain a selective sweep; FN is defined as the percentage of windows that contain a sweep and have not met the threshold value. Values are averaged across the 200 simulated replicates under population equilibrium with fixed recombination and mutation rates. Window size is 10kb.

| Population history | 2*N<sub>e</sub>s* of beneficial mutations | 95% tail |  |  | Null threshold |  |  |
| --- | --- | --- | --- | --- | --- | --- | --- |
|  |  | mean TP | mean FP | mean FN | mean TP | mean FP | mean FN |
| *N<sub>current</sub>* = *N<sub>ancestral</sub>* | No sweeps | 0.00 | 100.00 | 0.00 | - | - | 0.00 |
|  | 100 | 5.64 | 94.36 | 88.46 | 100.00 | 0.00 | 50.00 |
|  | 1000 | 49.65 | 50.35 | 57.08 | 95.02 | 4.98 | 70.12 |
|  | 10,000 | 98.39 | 1.61 | 84.29 | 96.70 | 3.30 | 15.74 |

In the absence of positive selection, the outlier approach is of course defined by 100% false positives, as any neutral distribution will naturally still contain 5% tails. Only in the presence of extremely strong positive selection (*2N_e_s* = 10,000) is the outlier approach observed to achieve considerable power. This high level of false positives associated with outlier approaches supports previous findings (Teshima et al. 2006; Thornton and Jensen 2007; Jensen et al. 2008). In non-equilibrium populations, power was greatly reduced with both the null threshold and genomic scan approaches with both SweepFinder2 and H12 (Supplementary Tables S2-9). The loss of power is particularly evident in the 90% population contraction models, reiterating our findings discussed above. Importantly however, it should be noted that using an evolutionarily appropriate baseline model generally resulted in lower FP rates and higher TP rates, providing a more desirably conservative approach.

### Concluding Thoughts

Our general results demonstrate that there exists relatively little power to detect recurrent selective sweeps using either frequency spectrum- or haplotype-based approaches, unless beneficial selection coefficients are large enough to result in large sweep effects whilst not so large so as to sweep entire genetic backgrounds, and beneficial mutation rates are high enough such that sweeps may be recent on average whilst not so high that sweep patterns overlap. Although there are numerous polymorphism-based sweep inference approaches (see the review of Stephan 2019), they generally rely fundamentally upon the SFS and/or LD-based patterns here evaluated, and thus will likely be subject to the same constraints that we have described. The type of recurrent sweep model here considered is in many ways the most appropriate for studying selective sweep effects, as beneficial mutations are naturally characterized by a rate of input as here modelled; whereas the common assumption that a strong single sweep reached fixation on an otherwise neutral background immediately prior to sampling is difficult to justify.

In examining performance under this model, we have here relaxed three additional undesirable assumptions common in sweep scans. Firstly, it is common to model only neutral and beneficial mutations, whereas in reality the majority of mutations in functional regions are expected to be deleterious. Hence, we have here modelled a full DFE including strongly deleterious, weakly deleterious, neutral, and beneficial mutations. Secondly, it is standard to assume fixed mutation and recombination rates across the genomic region in question, whereas in reality these rates are generally heterogeneous. Hence, we have here modelled this heterogeneity, demonstrating important trade-offs in these effects. Finally, it is standard to perform sweep scans under the assumption that these detected patterns are robust to underlying demographic effects, whereas in reality population size change histories are known to be problematic. Thus, we have here modelled multiple demographic histories, demonstrating the impact on resulting true- and false-positive rates even when these histories are accurately known *a priori* and are part of the baseline model.

On that point, it is important to emphasize that we have here considered a best-case scenario, in which a researcher has carefully inferred underlying mutation and recombination rates, the deleterious distribution of fitness effects, and population history - as has previously been recommended prior to performing sweep scans (Johri *et al*. 2020, 2022a,b). Although this outlook may appear bleak, it is important to quantify the expected performance of these commonly used inference approaches under these increasingly realistic population models. Specifically, there exist particular demographic histories in which sweep detection will not be feasible, and the statistical power across a genome will be dependent on the local mutation and recombination rates in a given genomic window. Apart from emphasizing the importance of estimating an appropriate baseline model in order to define these expectations and reduce false positive rates, this realization also highlights the limited visibility on the weakly and moderately beneficial tail of the DFE provided by polymorphism-based inference - a class about which we continue to have relatively little knowledge even in well-studied species.

### Materials and Methods Simulations

We simulated a single population using the forward-in-time software SLiM 3.7 (Haller and Messer 2019). For simulations with only effectively neutral mutations, a single 1Mb region was simulated. For simulations that included deleterious mutations, each simulation replicate consisted of 127 genes separated by intergenic regions of size 3,811bp. Each gene contained four exons (of size 588bp) and three introns (of six 563bp), forming a gene length of 4,041bp, and a total chromosome length of 997,204bp. See Supplementary Figure S32 for a schematic of chromosome structure. The numbers and lengths of exons, introns and intergenic regions were used to simulate a chromosome with *Drosophila melanogaster*-type structure and parameterizations, with averages estimated from Ensembl’s BDGP6.32 dataset (Adams et al. 2000), obtained from Ensemble release 107 (Cunningham et al. 2022). A fixed recombination rate of 2.32cM/Mb was taken from Comeron et al.’s (2012) genome-wide average estimate. The per site per generation mutation rate was taken from Keightley et al. (2014). This rate of 2.8e-9 implies an effective population size of 1.4 million.

Mutations in intronic and intergenic regions were modelled as effectively neutral, whilst exonic mutations were drawn from a DFE comprised of four fixed classes (Johri et al. 2020), whose frequencies are denoted by *f_i_*: *f_0_* with 0 ≤ 2*N_e_s* < 1 (*i.e*., effectively neutral mutations), *f_1_* with 1 ≤ 2*N_e_s* < 10 (*i.e*., weakly deleterious mutations), *f_2_* with 10 ≤ 2*N_e_s* < 100 (*i.e*., moderately deleterious mutations), and *f_3_* with 100 ≤ 2*N_e_s* < 2*N_e_* (*i.e*., strongly deleterious mutations), where *N_e_* is the effective population size and *s* is the reduction in fitness of the mutant homozygote relative to wild-type. Within each bin, *s* was drawn from a uniform distribution. We utilized the DFE inferred by Johri et al. (2020) in *D. melanogaster*. When simulating recurrent selective sweeps, beneficial mutations were incorporated into the DFE, though the 2*N_e_s* value for beneficials was fixed (as opposed to uniformly distributed within a range). The frequency of beneficials (*f_B_*) was incorporated into the DFE by subtracting it from the frequency of effectively neutral mutations (*i.e*., *f_0_* is set to *f_0_ – f_B_*).

For each replicate, 100 chromosomes were sampled after 17*N* generations (a 16*N* generation burn-in followed by any demographic change - see below - with sampling *N* generations later). For each scenario 200 replicates were simulated. Following the approach of Hill and Robertson (1966), all parameters were scaled down 200-fold in order to reduce runtimes, resulting in an initial population size of *N* = 7,000.

For each scenario a separate “null” run with no beneficial mutations was simulated as the baseline for the sweep detection plots. See below for further information.

### Simulating demographic change and variable recombination and mutation rates

We simulated four scenarios: fixed mutation rate and fixed recombination rate, fixed mutation rate and variable recombination rate, variable mutation rate and fixed recombination rate, and variable mutation rate and variable recombination rate. Where rates were variable, each 10kb region of the simulated chromosome had a different rate. Rates were drawn from a uniform distribution such that the chromosome-wide average was approximately the fixed rate. For variable recombination rates, the minimum and maximum parameters of the uniform distribution were 0.0127 and 7.3993cM/Mb, respectively. The maximum value is the maximum value in the sex-averaged Comeron et al. (2012) *D. melanogaster* recombination map. For variable mutation rates the minimum and maximum parameters of the uniform distribution were set at 1.5e-9 and 4.825e-9, to give a mean rate across each replicate that was equal to the fixed rate.

Four demographic scenarios were simulated: demographic equilibrium, 2x instantaneous population expansion, 0.5x instantaneous population contraction, and 0.1x instantaneous population contraction. The size change occured after 16*N* generations, and *N* generations prior to sampling.

### Detecting selective sweeps with SweepFinder2

SweepFinder2 was run on each simulated replicate to detect selective sweeps. We generated allele frequency files for each replicate. Because we have information on whether alleles are derived or ancestral, we followed Huber et al. (2016) in including only polymorphisms and substitutions. Inference was performed at each SNP via a grid file, following Nielsen et al. (2005). The background SFS was taken from the sweep-free null simulations. The following command line was used for inference:

SweepFinder2 –lu GridFile FreqFile SpectFile OutFile

For the variable recombination rate scenarios, SweepFinder2 was run with recombination rate information to improve inference power. In this case, the following command line was used for inference:

SweepFinder2 –lru GridFile FreqFile SpectFile RecFile OutFile

The maximum CLR value across the null run of simulations was set as the minimum threshold for detecting selective sweeps in Figures 2 and 3

### Detecting selective sweeps with H12

We used the H12 method of Garud et al. (2015) on each simulated replicate to detect selective sweeps, using a custom python script to implement the approach. H12 was estimated over 1kb windows at each SNP, with the SNP at the center of each window.

As with SweepFinder2, a baseline H12 was estimated using the “null” run of simulations, with the maximum H12 value across these replicates set as the minimum threshold for detecting selective sweeps in Figures 2 and 3 However, for both SweepFinder2 and H12, ROC curves were also generated by combining the inference results from all 200 replicates using Python’s Scikit-learn library (Pedregosa et al. 2011).

### Generating ROC curves

True positive rates (TPR) and false positive rates (FPR) were calculated across 1kb and 10kb non-overlapping windows. If a SNP was within 500bp of a beneficial mutation that had fixed within 0.5*N* generations of sampling, and the inference threshold was met, then the window containing that SNP was defined as a true positive. Furthermore, adjacent windows that met the inference threshold were also defined as true positives to account for hitchhiking effects. Adjacent windows to these would also be defined as true positives in a sequential pattern until a window that did not meet the threshold was encountered. This was performed across all 200 simulation replicates, with the results combined, for a range of threshold values. The metrics.roc_curve function (Pedregosa et al. 2011) from python’s scikit learn library was used to generate minimum and maximum thresholds. We then generated 1000 thresholds between the minimum and maximum values at which to estimate TPRs and FPRs.

To better quantify performance when mutation or recombination rates are variable, inference results were split into 10kb rate regions, binning the top 50% and bottom 50% of rates to enable direct comparison (see Results).

### Estimating divergence

Coding and non-coding divergence for each gene within a simulation replicate were estimated using a custom python script.

### Calculating summary statistics

Summary statistics were calculated across 1kb sliding windows with a step size of 500bp, using the python implementation of Libsequence (Thornton, 2003) via a custom script.

## ACKNOWLEDGEMENTS

This research was conducted using resources provided by Research Computing at Arizona State University (http://www.researchcomputing.asu.edu) and the Open Science Grid, which is supported by the National Science Foundation and the U.S. Department of Energy’s Office of Science. This work was funded by National Institutes of Health grant R35GM139383 to JDJ.

---

**Table S1:** Proportion of fixed mutations that are beneficial across all simulated generations, and across the last 0.5*N* generations before sampling.

| $2N_e s$ | Proportion of fixations that are beneficial mutations | |
| --- | --- | --- |
| | All generations | Within $0.5N$ generations of sampling |
| 100 | 0.00023 | 0.000178 |
| 1000 | 0.002223 | 0.001813 |
| 10,000 | 0.01285 | 0.011793 |

**Table S2:** Comparing the power of **H12** using the evolutionarily appropriate null model, with an outlier approach that assumes windows in the 5% right-hand tail contain selective sweeps. Here TP is defined as the percentage of windows that have met the threshold value and contain a selective sweep; FP is defined as the percentage of windows that have met the threshold value and do not contain a selective sweep; FN is defined as the percentage of windows that contain a sweep and have not met the threshold value. Values are averaged across the 200 simulation replicates under various population histories, **with fixed recombination and mutation rates**. Window size is 10kb.

| Population history | $2N_e$s of beneficial mutations | 95% tail | | | Null threshold | | |
| --- | --- | --- | --- | --- | --- | --- | --- |
|  |  | mean TP | mean FP | mean FN | mean TP | mean FP | mean FN |
| $N_{current} = N_{ancestral}$ | No sweeps | 0.00 | 100.00 | 0.00 | - | - | 0.00 |
|  | 100 | 5.10 | 94.90 | 87.25 | 0.00 | 0.00 | 100.00 |
|  | 1000 | 55.81 | 44.19 | 50.02 | 84.01 | 15.99 | 77.52 |
|  | 10,000 | 96.28 | 3.72 | 84.75 | 97.27 | 2.73 | 46.44 |
| $N_{current} = 2N_{ancestral}$ | No sweeps | 0.00 | 100.00 | 0.00 | - | - | 0.00 |
|  | 100 | 3.50 | 96.50 | 94.17 | 0.00 | 0.00 | 100.00 |
|  | 1000 | 76.60 | 23.40 | 60.70 | 76.44 | 23.56 | 86.42 |
|  | 10,000 | 95.11 | 4.89 | 90.69 | 99.07 | 0.93 | 32.86 |
| $N_{current} = 0.5N_{ancestral}$ | No sweeps | 0.00 | 100.00 | 0.00 | - | - | 0.00 |
|  | 100 | 5.71 | 94.29 | 88.39 | 0.00 | 0.00 | 100.00 |
|  | 1000 | 27.17 | 72.83 | 63.99 | 65.52 | 34.48 | 76.29 |
|  | 10,000 | 91.50 | 8.50 | 72.52 | 93.89 | 6.11 | 73.44 |
| $N_{current} = 0.1N_{ancestral}$ | No sweeps | 0.00 | 100.00 | 0.00 | - | - | - |
|  | 100 | 2.38 | 97.62 | 91.67 | 2.38 | 97.62 | 91.67 |
|  | 1000 | 4.57 | 95.43 | 85.11 | 4.57 | 95.43 | 85.11 |
|  | 10,000 | 9.61 | 90.39 | 82.12 | 9.56 | 90.44 | 82.96 |

**Table S3:** Comparing the power of **SweepFinder2** using the evolutionarily appropriate null model, with an outlier approach that assumes windows in the 5% right-hand tail contain selective sweeps. Here TP is defined as the percentage of windows that have met the threshold value and contain a selective sweep; FP is defined as the percentage of windows that have met the threshold value and do not contain a selective sweep; FN is defined as the percentage of windows that contain a sweep and have not met the threshold value. Values are averaged across the 200 simulation replicates under various population histories, **with fixed recombination and mutation rates**. Window size is 10kb.

| Population history | $2N_e$s of beneficial mutations | 95% tail | | | Null threshold | | |
| --- | --- | --- | --- | --- | --- | --- | --- |
|  |  | mean TP | mean FP | mean FN | mean TP | mean FP | mean FN |
| $N_{current} = N_{ancestral}$ | No sweeps | 0.00 | 100.00 | 0.00 | - | - | 0.00 |
|  | 100 | 5.64 | 94.36 | 88.46 | 100.00 | 0.00 | 50.00 |
|  | 1000 | 49.65 | 50.35 | 57.08 | 95.02 | 4.98 | 70.12 |
|  | 10,000 | 98.39 | 1.61 | 84.29 | 96.70 | 3.30 | 15.74 |
| $N_{current} = 2N_{ancestral}$ | No sweeps | 0.00 | 100.00 | 0.00 | - | - | 0.00 |
|  | 100 | 6.00 | 94.00 | 89.38 | 0.00 | 0.00 | 100.00 |
|  | 1000 | 79.20 | 20.80 | 59.76 | 89.37 | 10.63 | 62.16 |
|  | 10,000 | 99.00 | 1.00 | 90.58 | 96.93 | 3.07 | 2.34 |
| $N_{current} = 0.5N_{ancestral}$ | No sweeps | 0.00 | 100.00 | 0.00 | - | - | 0.00 |
|  | 100 | 4.29 | 95.71 | 89.29 | 0.00 | 0.00 | 100.00 |
|  | 1000 | 19.50 | 80.50 | 73.56 | 73.21 | 26.79 | 75.00 |
|  | 10,000 | 96.50 | 3.50 | 71.28 | 97.93 | 2.07 | 54.72 |
| $N_{current} = 0.1N_{ancestral}$ | No sweeps | 0.00 | 100.00 | 0.00 | - | - | 0.00 |
|  | 100 | 3.33 | 96.67 | 91.67 | 0.00 | 0.00 | 100.00 |
|  | 1000 | 2.61 | 97.39 | 94.57 | 0.00 | 100.00 | 100.00 |
|  | 10,000 | 17.25 | 82.75 | 83.04 | 80.15 | 19.85 | 62.59 |

**Table S4:** Comparing the power of **H12** using the evolutionarily appropriate null model, with an outlier approach that assumes windows in the 5% right-hand tail contain selective sweeps. Here TP is defined as the percentage of windows that have met the threshold value and contain a selective sweep; FP is defined as the percentage of windows that have met the threshold value and do not contain a selective sweep; FN is defined as the percentage of windows that contain a sweep and have not met the threshold value. Values are averaged across the 200 simulation replicates under various population histories, **with fixed recombination and variable mutation rates**. Window size is 10kb.

| Population history | $2N_e$s of beneficial mutations | 95% tail | | | Null threshold | | |
| --- | --- | --- | --- | --- | --- | --- | --- |
|  |  | mean TP | mean FP | mean FN | mean TP | mean FP | mean FN |
| $N_{current} = N_{ancestral}$ | No sweeps | 0.00 | 100.00 | 0.00 | - | - | 0.00 |
|  | 100 | 3.26 | 96.74 | 92.25 | 0.00 | 0.00 | 100.00 |
|  | 1000 | 38.70 | 61.30 | 65.11 | 62.50 | 37.50 | 86.73 |
|  | 10,000 | 94.63 | 5.37 | 84.86 | 94.79 | 5.21 | 77.48 |
| $N_{current} = 2N_{ancestral}$ | No sweeps | 0.00 | 100.00 | 0.00 | - | - | 0.00 |
|  | 100 | 2.72 | 97.28 | 95.43 | 0.00 | 0.00 | 100.00 |
|  | 1000 | 66.67 | 33.33 | 66.99 | 62.11 | 37.89 | 92.66 |
|  | 10,000 | 95.85 | 4.15 | 90.59 | 96.57 | 3.43 | 48.74 |
| $N_{current} = 0.5N_{ancestral}$ | No sweeps | 0.00 | 100.00 | 0.00 | - | - | 0.00 |
|  | 100 | 4.62 | 95.38 | 89.10 | 0.00 | 0.00 | 100.00 |
|  | 1000 | 16.08 | 83.92 | 75.49 | 0.00 | 100.00 | 100.00 |
|  | 10,000 | 76.08 | 23.92 | 78.32 | 89.36 | 10.64 | 91.44 |
| $N_{current} = 0.1N_{ancestral}$ | No sweeps | 0.00 | 100.00 | 0.00 | - | - | 0.00 |
|  | 100 | 1.67 | 98.33 | 90.00 | 1.67 | 98.33 | 90.00 |
|  | 1000 | 3.56 | 96.44 | 83.18 | 3.56 | 96.44 | 83.18 |
|  | 10,000 | 6.99 | 93.01 | 79.17 | 6.99 | 93.01 | 79.17 |

**Table S5:**
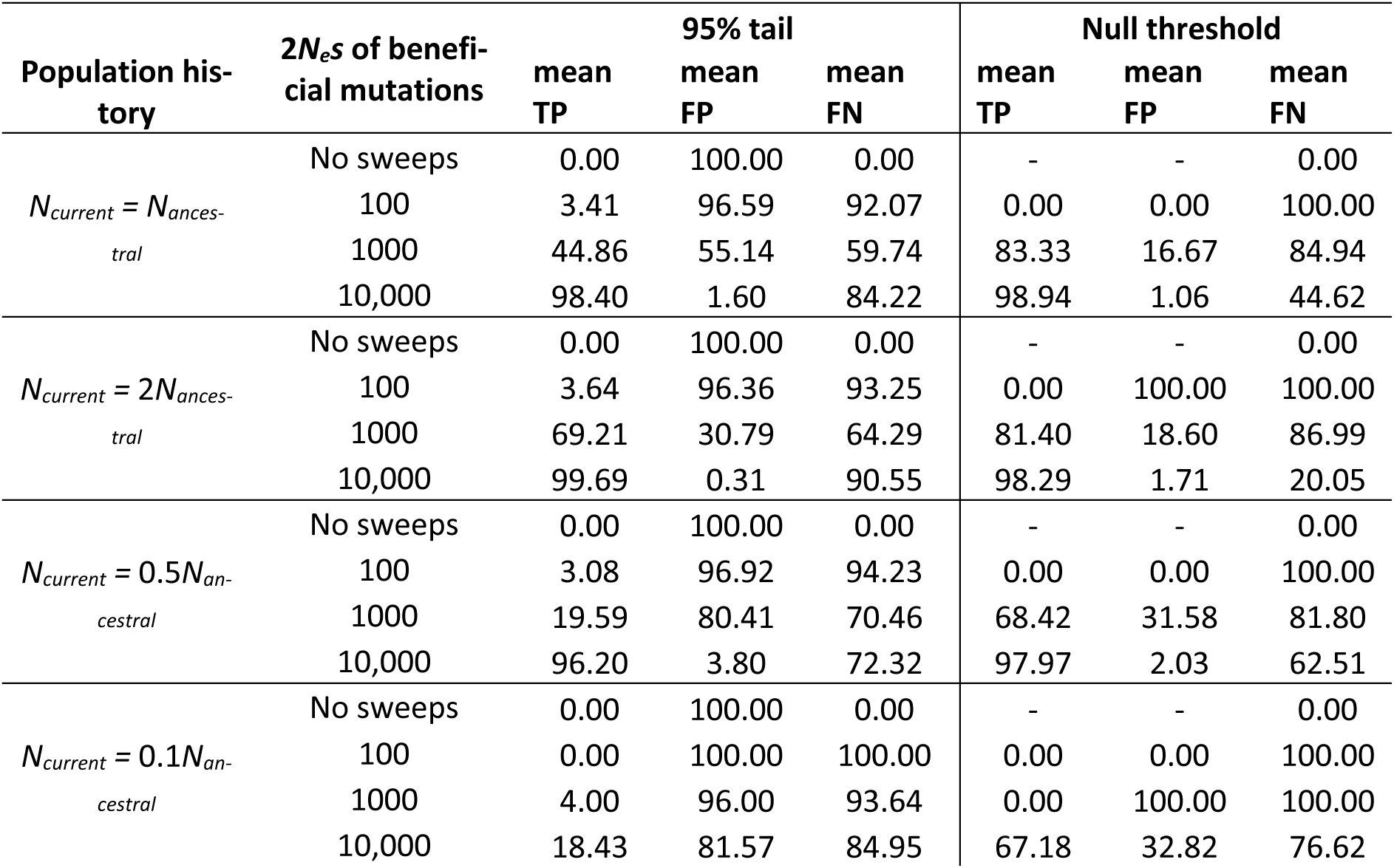
Comparing the power of **SweepFinder2** using the evolutionarily appropriate null model, with an outlier approach that assumes windows in the 5% right-hand tail contain selective sweeps. Here TP is defined as the percentage of windows that have met the threshold value and contain a selective sweep; FP is defined as the percentage of windows that have met the threshold value and do not contain a selective sweep; FN is defined as the percentage of windows that contain a sweep and have not met the threshold value. Values are averaged across the 200 simulation replicates under various population histories, **with fixed recombination and variable mutation rates**. Window size is 10kb.

| Population history | $2N_e s$ of beneficial mutations | 95% tail | | | Null threshold | | |
| --- | --- | --- | --- | --- | --- | --- | --- |
|  |  | mean TP | mean FP | mean FN | mean TP | mean FP | mean FN |
| $N_{current} = N_{ancestral}$ | No sweeps | 0.00 | 100.00 | 0.00 | - | - | 0.00 |
|  | 100 | 3.41 | 96.59 | 92.07 | 0.00 | 0.00 | 100.00 |
|  | 1000 | 44.86 | 55.14 | 59.74 | 83.33 | 16.67 | 84.94 |
|  | 10,000 | 98.40 | 1.60 | 84.22 | 98.94 | 1.06 | 44.62 |
| $N_{current} = 2N_{ancestral}$ | No sweeps | 0.00 | 100.00 | 0.00 | - | - | 0.00 |
|  | 100 | 3.64 | 96.36 | 93.25 | 0.00 | 100.00 | 100.00 |
|  | 1000 | 69.21 | 30.79 | 64.29 | 81.40 | 18.60 | 86.99 |
|  | 10,000 | 99.69 | 0.31 | 90.55 | 98.29 | 1.71 | 20.05 |
| $N_{current} = 0.5N_{ancestral}$ | No sweeps | 0.00 | 100.00 | 0.00 | - | - | 0.00 |
|  | 100 | 3.08 | 96.92 | 94.23 | 0.00 | 0.00 | 100.00 |
|  | 1000 | 19.59 | 80.41 | 70.46 | 68.42 | 31.58 | 81.80 |
|  | 10,000 | 96.20 | 3.80 | 72.32 | 97.97 | 2.03 | 62.51 |
| $N_{current} = 0.1N_{ancestral}$ | No sweeps | 0.00 | 100.00 | 0.00 | - | - | 0.00 |
|  | 100 | 0.00 | 100.00 | 100.00 | 0.00 | 0.00 | 100.00 |
|  | 1000 | 4.00 | 96.00 | 93.64 | 0.00 | 100.00 | 100.00 |
|  | 10,000 | 18.43 | 81.57 | 84.95 | 67.18 | 32.82 | 76.62 |

**Table S6:**
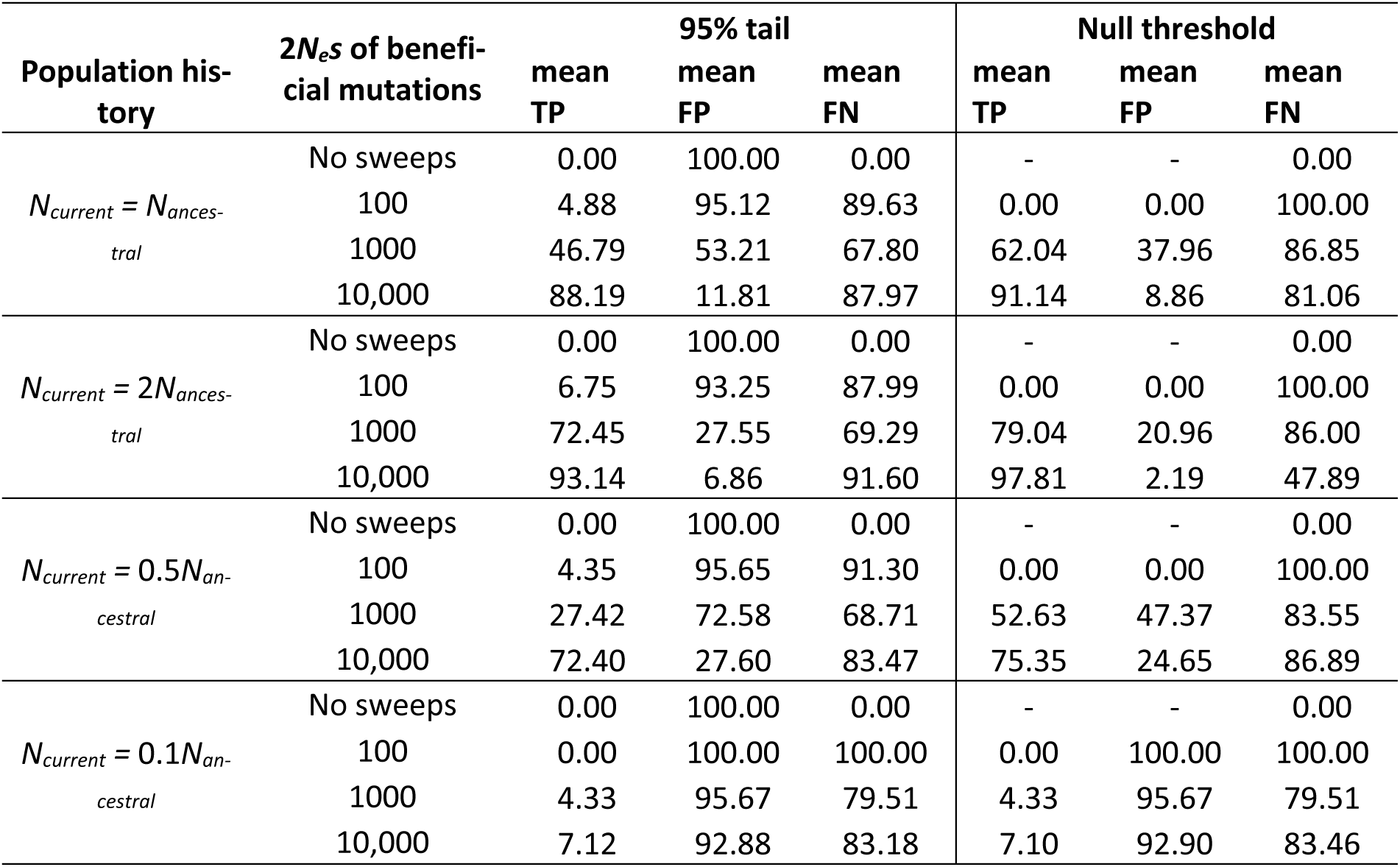
Comparing the power of **H12** using the evolutionarily appropriate null model, with an outlier approach that assumes windows in the 5% right-hand tail contain selective sweeps. Here TP is defined as the percentage of windows that have met the threshold value and contain a selective sweep; FP is defined as the percentage of windows that have met the threshold value and do not contain a selective sweep; FN is defined as the percentage of windows that contain a sweep and have not met the threshold value. Values are averaged across the 200 simulation replicates under various population histories, **with variable recombination and fixed mutation rates**. Window size is 10kb.

**Table S7:** Comparing the power of **Sweepfinder2** using the evolutionarily appropriate null model, with an outlier approach that assumes windows in the 5% right-hand tail contain selective sweeps. Here TP is defined as the percentage of windows that have met the threshold value and contain a selective sweep; FP is defined as the percentage of windows that have met the threshold value and do not contain a selective sweep; FN is defined as the percentage of windows that contain a sweep and have not met the threshold value. Values are averaged across the 200 simulation replicates under various population histories, **with variable recombination and fixed mutation rates**. Window size is 10kb.

| Population history | $2N_e$s of beneficial mutations | 95% tail | | | Null threshold | | |
| --- | --- | --- | --- | --- | --- | --- | --- |
|  |  | mean TP | mean FP | mean FN | mean TP | mean FP | mean FN |
| $N_{current} = N_{ancestral}$ | No sweeps | 0.00 | 100.00 | 0.00 | - | - | 0.00 |
|  | 100 | 6.50 | 93.50 | 89.38 | 0.00 | 0.00 | 100.00 |
|  | 1000 | 51.32 | 48.68 | 64.37 | 65.16 | 34.84 | 79.61 |
|  | 10,000 | 81.82 | 18.18 | 89.75 | 86.29 | 13.71 | 54.78 |
| $N_{current} = 2N_{ancestral}$ | No sweeps | 0.00 | 100.00 | 0.00 | - | - | 0.00 |
|  | 100 | 8.31 | 91.69 | 86.58 | 0.00 | 100.00 | 100.00 |
|  | 1000 | 71.70 | 28.30 | 69.53 | 80.22 | 19.78 | 69.52 |
|  | 10,000 | 90.40 | 9.60 | 92.99 | 94.83 | 5.17 | 18.66 |
| $N_{current} = 0.5N_{ancestral}$ | No sweeps | 0.00 | 100.00 | 0.00 | - | - | 0.00 |
|  | 100 | 4.55 | 95.45 | 89.77 | 0.00 | 100.00 | 100.00 |
|  | 1000 | 23.62 | 76.38 | 75.52 | 54.63 | 45.37 | 77.93 |
|  | 10,000 | 69.20 | 30.80 | 84.77 | 75.88 | 24.12 | 71.60 |
| $N_{current} = 0.1N_{ancestral}$ | No sweeps | 0.00 | 100.00 | 0.00 | - | - | 0.00 |
|  | 100 | 0.00 | 100.00 | 100.00 | 0.00 | 0.00 | 100.00 |
|  | 1000 | 2.08 | 97.92 | 96.35 | 0.00 | 100.00 | 100.00 |
|  | 10,000 | 9.89 | 90.11 | 90.84 | 41.46 | 58.54 | 77.17 |

**Table S8:** Comparing the power of **H12** using the evolutionarily appropriate null model, with an outlier approach that assumes windows in the 5% right-hand tail contain selective sweeps. Here TP is defined as the percentage of windows that have met the threshold value and contain a selective sweep; FP is defined as the percentage of windows that have met the threshold value and do not contain a selective sweep; FN is defined as the percentage of windows that contain a sweep and have not met the threshold value. Values are averaged across the 200 simulation replicates under various population histories, **with variable recombination and mutation rates**. Window size is 10kb.

| Population history | $2N_e s$ of beneficial mutations | 95% tail | | | Null threshold | | |
| --- | --- | --- | --- | --- | --- | --- | --- |
|  |  | mean TP | mean FP | mean FN | mean TP | mean FP | mean FN |
| $N_{current} = N_{ancestral}$ | No sweeps | 0.00 | 100.00 | 0.00 | - | - | 0.00 |
|  | 100 | 3.20 | 96.80 | 92.45 | 0.00 | 100.00 | 100.00 |
|  | 1000 | 35.56 | 64.44 | 72.69 | 73.33 | 26.67 | 89.20 |
|  | 10,000 | 81.53 | 18.47 | 88.53 | 83.99 | 16.01 | 84.73 |
| $N_{current} = 2N_{ancestral}$ | No sweeps | 0.00 | 100.00 | 0.00 | - | - | 0.00 |
|  | 100 | 4.39 | 95.61 | 91.87 | 0.00 | 100.00 | 100.00 |
|  | 1000 | 62.11 | 37.89 | 76.38 | 71.01 | 28.99 | 86.65 |
|  | 10,000 | 89.77 | 10.23 | 91.49 | 92.45 | 7.55 | 54.11 |
| $N_{current} = 0.5N_{ancestral}$ | No sweeps | 0.00 | 100.00 | 0.00 | - | - | 0.00 |
|  | 100 | 4.80 | 95.20 | 91.00 | 0.00 | 0.00 | 100.00 |
|  | 1000 | 15.76 | 84.24 | 80.40 | 80.00 | 20.00 | 76.67 |
|  | 10,000 | 63.15 | 36.85 | 84.22 | 61.60 | 38.40 | 93.71 |
| $N_{current} = 0.1N_{ancestral}$ | No sweeps | 0.00 | 100.00 | 0.00 | - | - | 0.00 |
|  | 100 | 3.36 | 96.64 | 66.67 | 3.36 | 96.64 | 66.67 |
|  | 1000 | 1.66 | 98.34 | 84.00 | 1.66 | 98.34 | 84.00 |
|  | 10,000 | 7.72 | 92.28 | 73.05 | 7.72 | 92.28 | 73.05 |

**Table S9:**
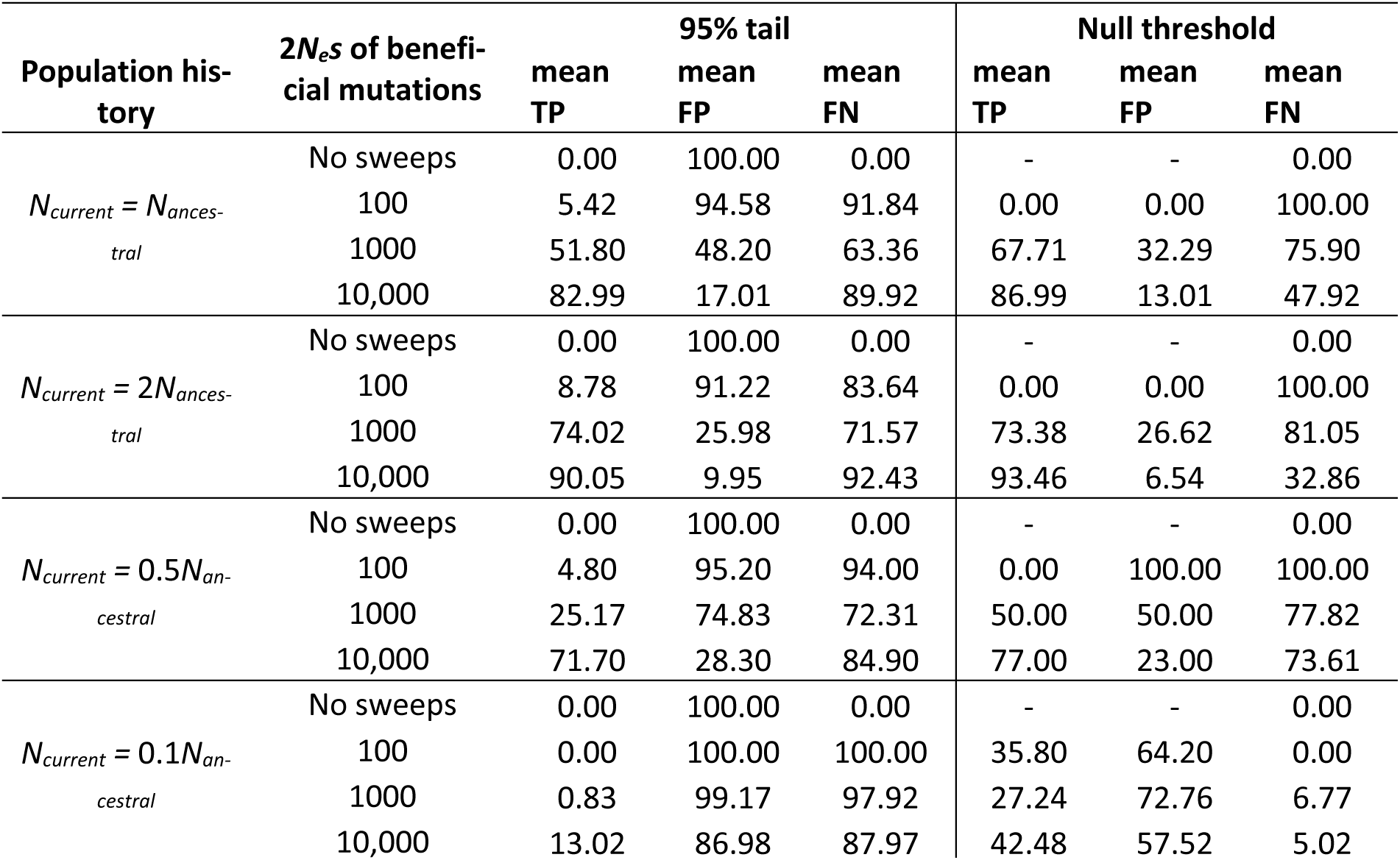
Comparing the power of **SweepFinder2** using the evolutionarily appropriate null model, with an outlier approach that assumes windows in the 5% right-hand tail contain selective sweeps. Here TP is defined as the percentage of windows that have met the threshold value and contain a selective sweep; FP is defined as the percentage of windows that have met the threshold value and do not contain a selective sweep; FN is defined as the percentage of windows that contain a sweep and have not met the threshold value. Values are averaged across the 200 simulation replicates under various population histories, **with variable recombination and mutation rates**. Window size is 10kb.

| Population history | $2N_e$s of beneficial mutations | 95% tail | | | Null threshold | | |
| --- | --- | --- | --- | --- | --- | --- | --- |
|  |  | mean TP | mean FP | mean FN | mean TP | mean FP | mean FN |
| $N_{current} = N_{ancestral}$ | No sweeps | 0.00 | 100.00 | 0.00 | - | - | 0.00 |
|  | 100 | 5.42 | 94.58 | 91.84 | 0.00 | 0.00 | 100.00 |
|  | 1000 | 51.80 | 48.20 | 63.36 | 67.71 | 32.29 | 75.90 |
|  | 10,000 | 82.99 | 17.01 | 89.92 | 86.99 | 13.01 | 47.92 |
| $N_{current} = 2N_{ancestral}$ | No sweeps | 0.00 | 100.00 | 0.00 | - | - | 0.00 |
|  | 100 | 8.78 | 91.22 | 83.64 | 0.00 | 0.00 | 100.00 |
|  | 1000 | 74.02 | 25.98 | 71.57 | 73.38 | 26.62 | 81.05 |
|  | 10,000 | 90.05 | 9.95 | 92.43 | 93.46 | 6.54 | 32.86 |
| $N_{current} = 0.5N_{ancestral}$ | No sweeps | 0.00 | 100.00 | 0.00 | - | - | 0.00 |
|  | 100 | 4.80 | 95.20 | 94.00 | 0.00 | 100.00 | 100.00 |
|  | 1000 | 25.17 | 74.83 | 72.31 | 50.00 | 50.00 | 77.82 |
|  | 10,000 | 71.70 | 28.30 | 84.90 | 77.00 | 23.00 | 73.61 |
| $N_{current} = 0.1N_{ancestral}$ | No sweeps | 0.00 | 100.00 | 0.00 | - | - | 0.00 |
|  | 100 | 0.00 | 100.00 | 100.00 | 35.80 | 64.20 | 0.00 |
|  | 1000 | 0.83 | 99.17 | 97.92 | 27.24 | 72.76 | 6.77 |
|  | 10,000 | 13.02 | 86.98 | 87.97 | 42.48 | 57.52 | 5.02 |

**Figure S1:**
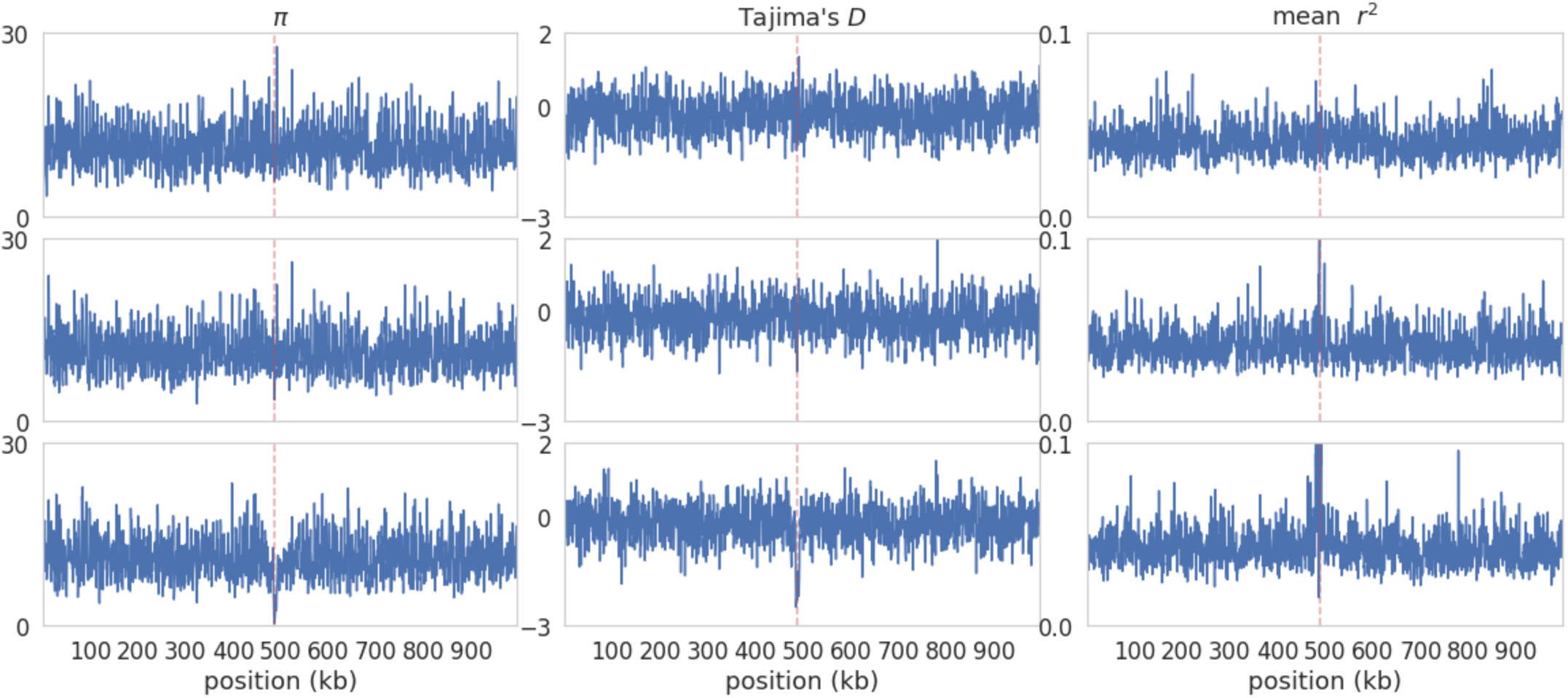
Example patterns around a single selective sweep across 1kb overlapping windows, with a step size of 500bp, for different values of 2*N_e_s*, from a single simulation replicate. **These summary statistics correspond to the sweep inference results in** Figure 2 **of the main text**. In each case, 2*N_e_s* values go from lowest (top panel) to highest: 100; 1000; 10,000. The red dashed line is the position of the beneficial mutation.

**Figure S2:**
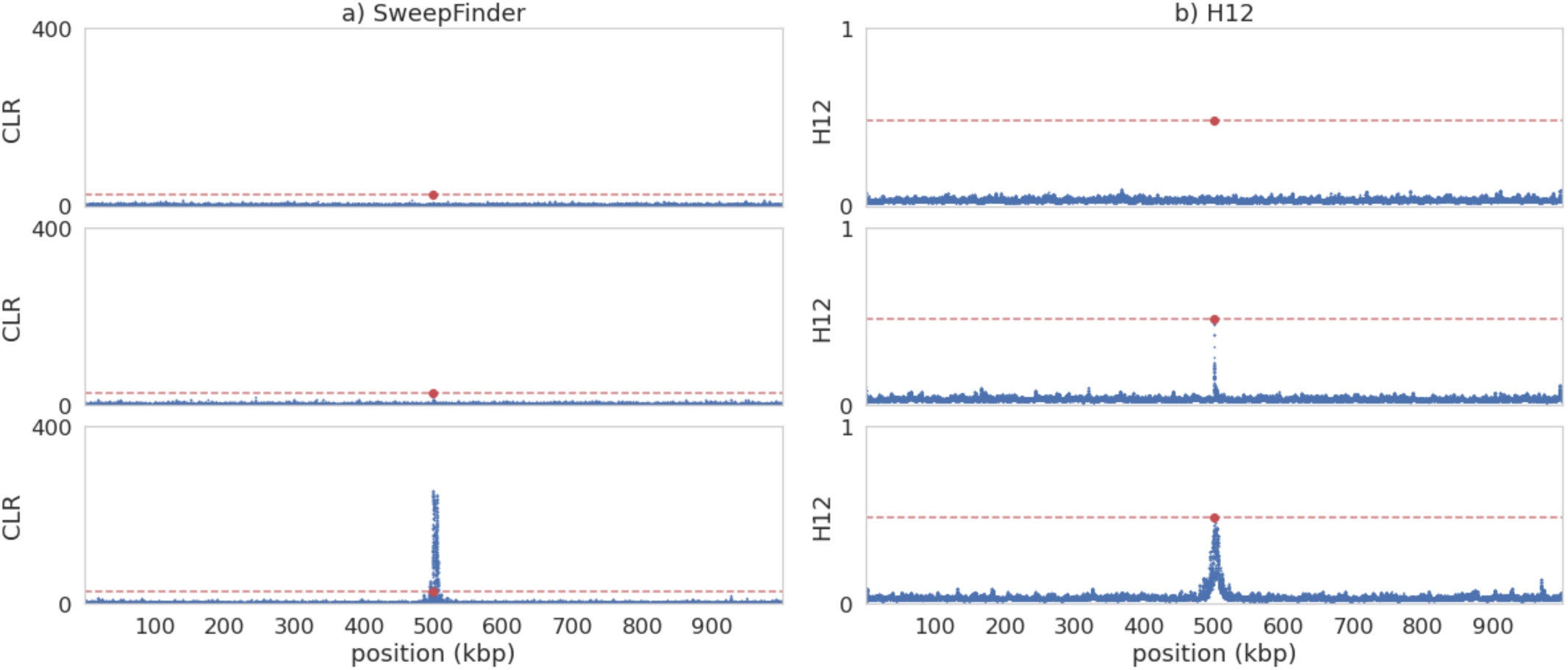
Example patterns around a single selective sweep for different values of 2*N_e_s* in an equilibrium population with fixed mutation and recombination rates. **Unlike** Figure 2 **in the main text, here sweeps have occurred on a completely neutral background (*i.e*., no underlying DFE).** In each case, 2*N_e_s* values go from lowest (top panel) to highest: 100; 1000; 10,000. The red data point is the position of the beneficial fixation. a) Inference results from SweepFinder2. Blue data points are CLR values inferred for each window. The red dashed line is the threshold for sweep detection, determined by the highest CLR value across 200 simulation replicates in which no beneficial mutations are modelled. Inference was performed at each SNP (see Methods section for further details). b) Sweep inference with the H12 statistic. Blue data points are H12 values estimated for each window. As with SweepFinder2, the red dashed line is the threshold for sweep detection. Inference was performed across 1kb windows for each SNP, with the SNP at the center of each window.

**Figure S3:**
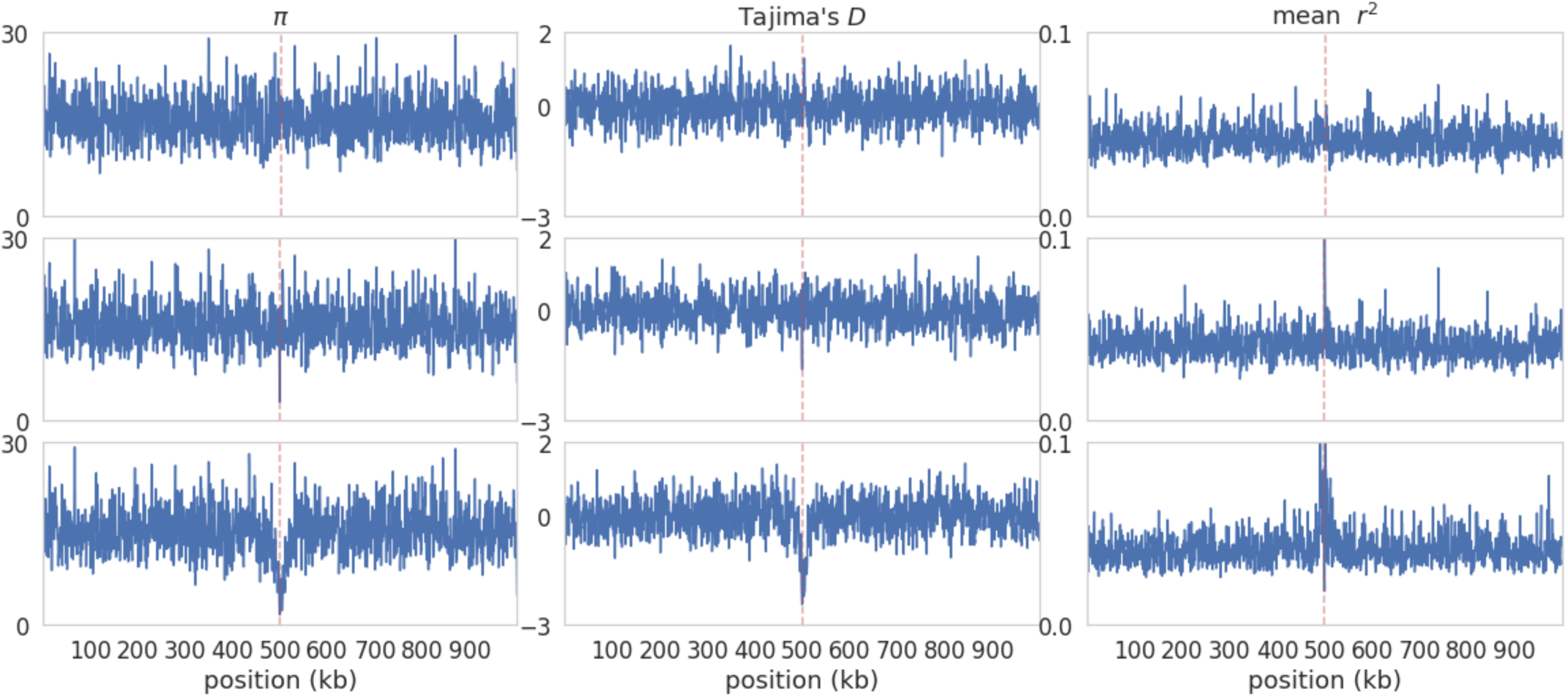
Example patterns around a single selective sweep across 1kb overlapping windows, with a step size of 500bp, for different values of 2*N_e_s*, from a single simulation replicate. **Unlike Figure S1, here sweeps are occurring on completely neutral backgrounds (*i.e*., no underlying DFE).** In each case, 2*N_e_s* values go from lowest (top panel) to highest: 100; 1000; 10,000. The red dashed line is the position of the beneficial mutation.

**Figure S4:**
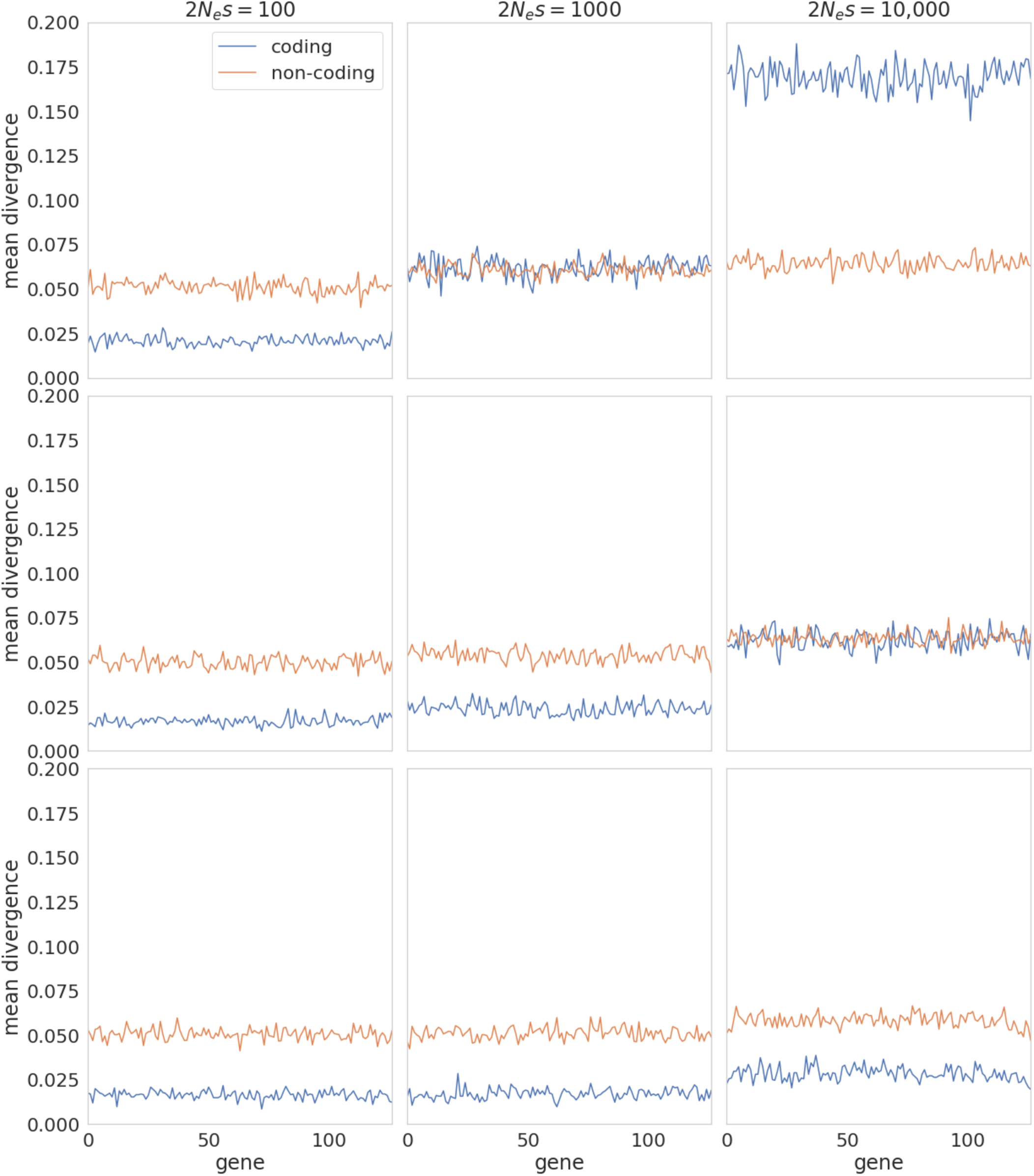
Per-gene coding and non-coding divergence for a single simulation replicate for different values of 2*N_e_s*. In the top row the proportion of mutations that are beneficial is 5%; middle row is 0.5%; bottom row is 0.05%.

**Figure S5:**
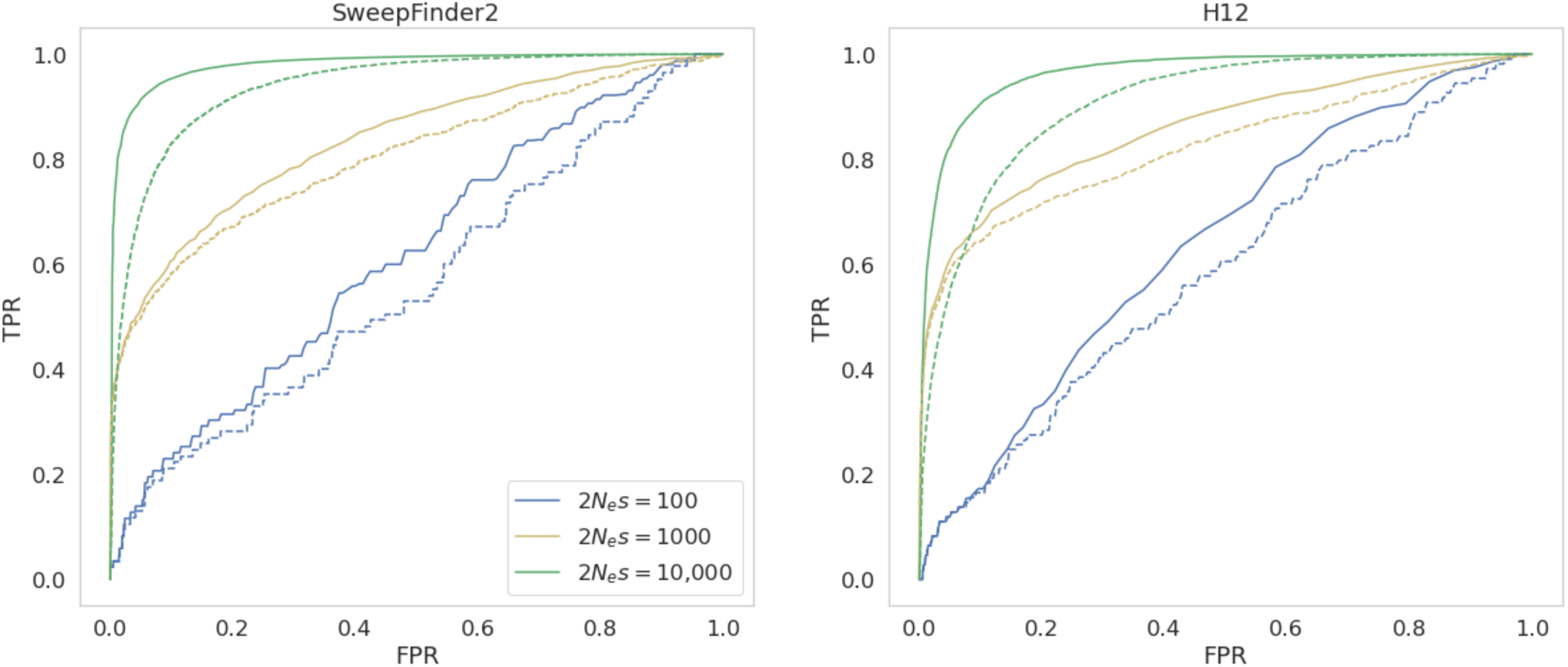
ROC curves comparing the standard method of classifying only windows containing a SNP that is within 500bp of a beneficial mutation and that meet the inference threshold as a true positive, with the “adjacent windows” method wherein adjacent windows that also meet the inference threshold are also defined as true positives. The filled line is for the “adjacent windows method”, whilst the dashed line is for the standard method. The ROC curves correspond to an equilibrium population, fixed rate model, with TPR and FPR calculated across 10kb windows.

**Figure S6:**
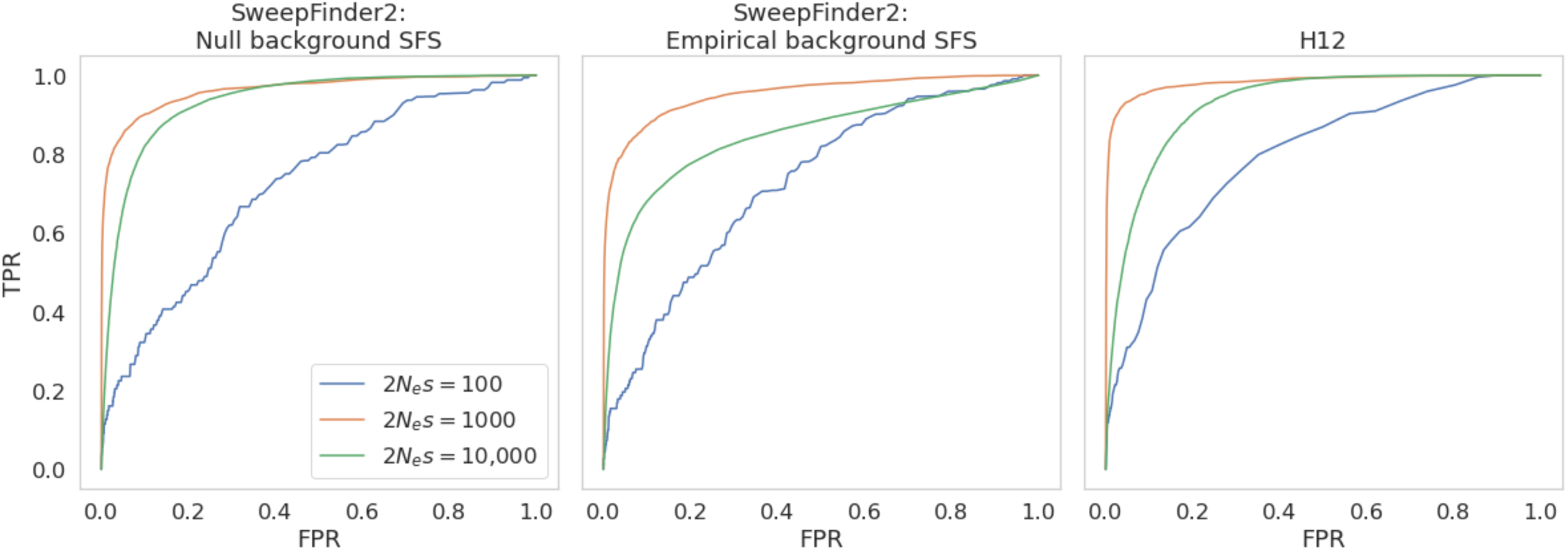
ROC curves, showing the change in true positive rate (TPR) as the false positive rate (FPR) increases, for sweep inference in an equilibrium population with fixed mutation and recombination rates across 200 simulation replicates, for 1kb windows. **a) ROC curves for SweepFinder2 when using a null background SFS** (*i.e.,* the background SFS is generated from a simulation run in which all else is modelled identically, except that there are no beneficial mutations**); b) ROC curves for SweepFinder2 when using an empirical background SFS** (*i.e.,* the background SFS is the empirical data itself); **c) ROC curves for H12**.

**Figure S7:**
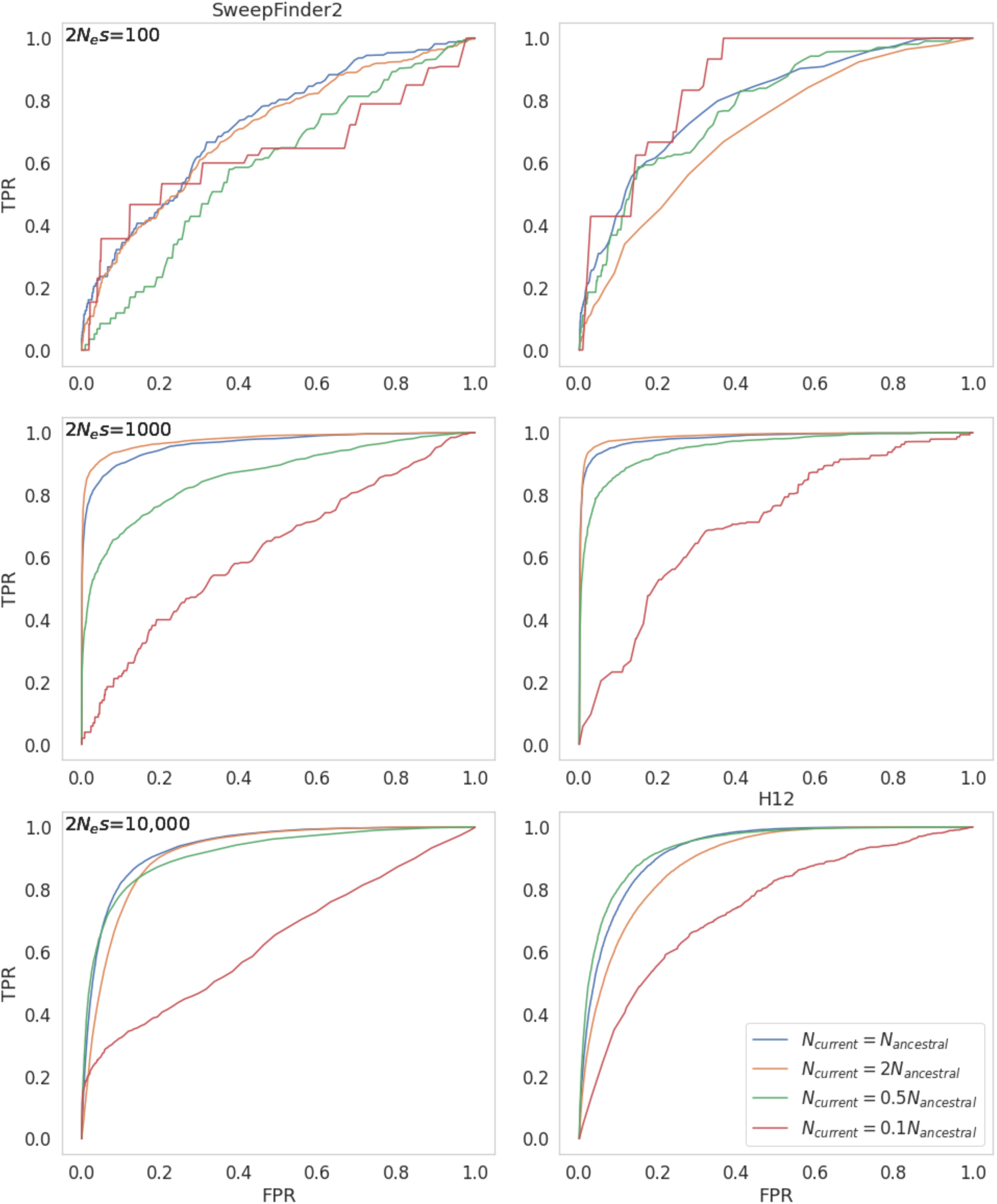
ROC curves, showing the change in true positive rate (TPR) as the false positive rate (FPR) increases, for sweep inference in populations with differing demographic histories, across 200 replicates each, **for windows of size 1kb**. The panels on the left are for inference with SweepFinder2, and on the right with the H12 statistic. Where population size change occurs, it is instantaneous, occurring 1*N* generations before sampling.

**Figure S8:**
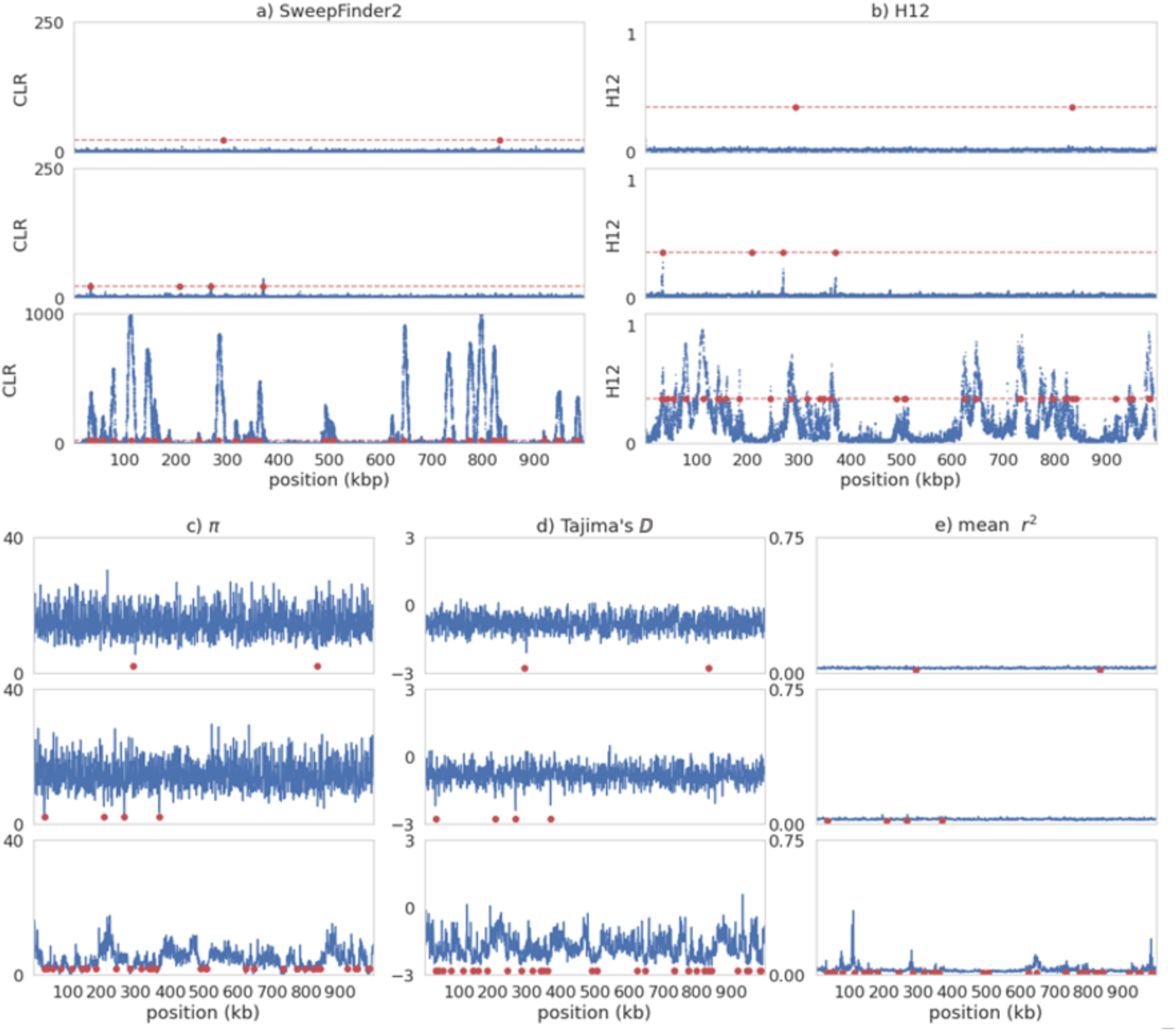
Sweep inference and summary statistics for a single simulation replicate of recurrent selective sweeps for different values of 2*N_e_s* **under instantaneous population expansion (such that *N_current_* = *2N_ancestral_*) with fixed mutation and recombination rates**. Population size change occurs 1*N* generations before sampling. In each case, 2*N_e_s* values go from lowest (top panel) to highest: 100; 1000; 10,000. For all panels, red data points are the positions of beneficial fixations with the previous 0.5*N* generations prior to sampling. a) Inference results from SweepFinder2. Blue data points are CLR values inferred for each window. The red dashed line is the threshold for sweep detection, determined by the highest CLR value across 200 simulation replicates in which no beneficial mutations are modelled. Inference was performed at each SNP (see Methods section for further details). b) Sweep inference with the H12 statistic. Blue data points are H12 values estimated for each window. As with SweepFinder2, the red dashed line is the threshold for sweep detection. Inference was performed across 1kb windows for each SNP, with the SNP at the center of each window. c-e) Summary statistics across the simulated region.

**Figure S9:**
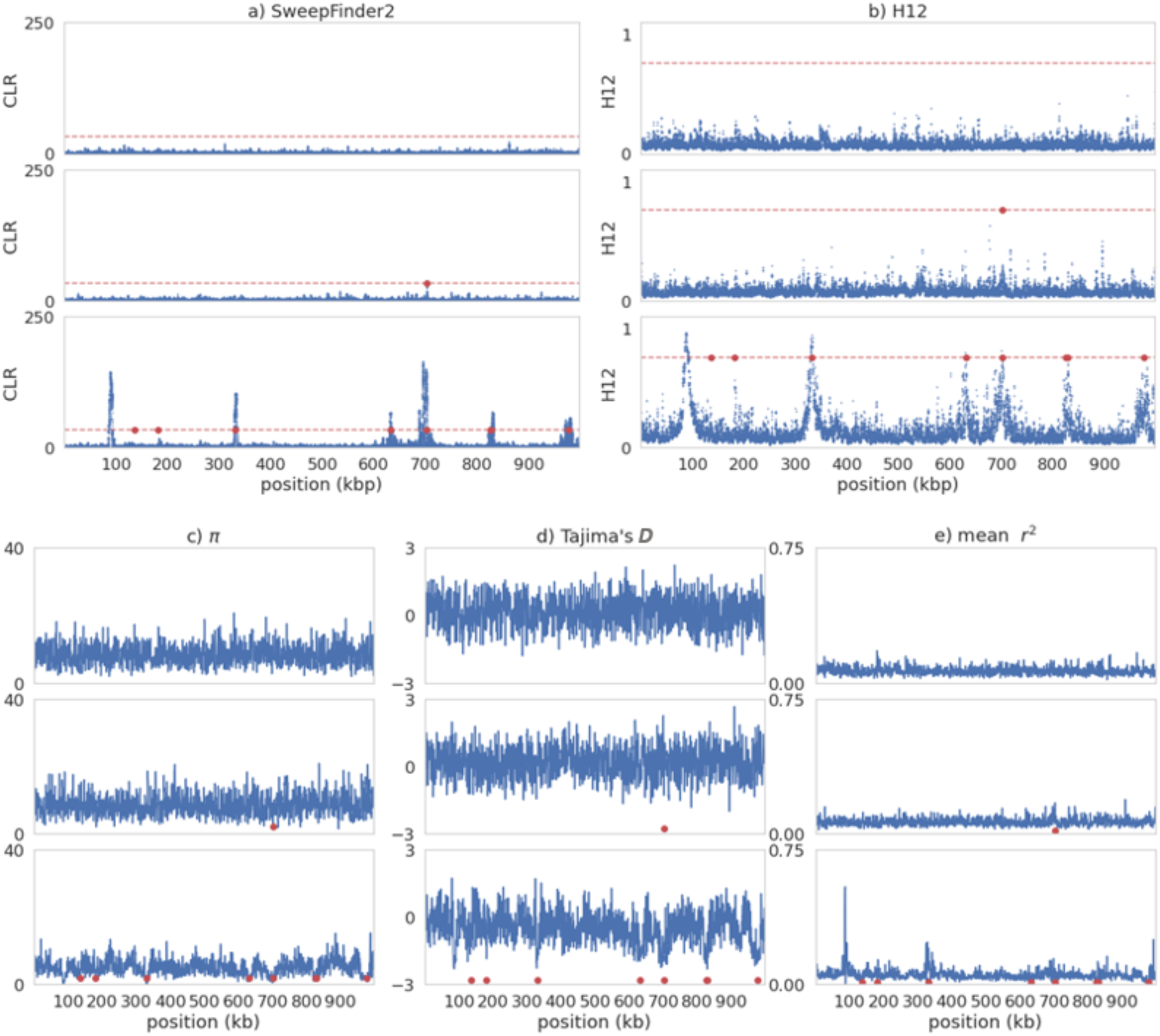
Sweep inference and summary statistics for a single simulation replicate of recurrent selective sweeps for different values of 2*N_e_s* **under instantaneous population contraction (such that *N_current_* = *0.5N_ancestral_*) with fixed mutation and recombination rates**. Population size change occurs 1*N* generations before sampling. In each case, 2*N_e_s* values go from lowest (top panel) to highest: 100; 1000; 10,000. For all panels, red data points are the positions of beneficial fixations with the previous 0.5*N* generations prior to sampling. a) Inference results from SweepFinder2. Blue data points are CLR values inferred for each window. The red dashed line is the threshold for sweep detection, determined by the highest CLR value across 200 simulation replicates in which no beneficial mutations are modelled. Inference was performed at each SNP (see Methods section for further details). b) Sweep inference with the H12 statistic. Blue data points are H12 values estimated for each window. As with SweepFinder2, the red dashed line is the threshold for sweep detection. Inference was performed across 1kb windows for each SNP, with the SNP at the center of each window. c-e) Summary statistics across the simulated region.

**Figure S10:**
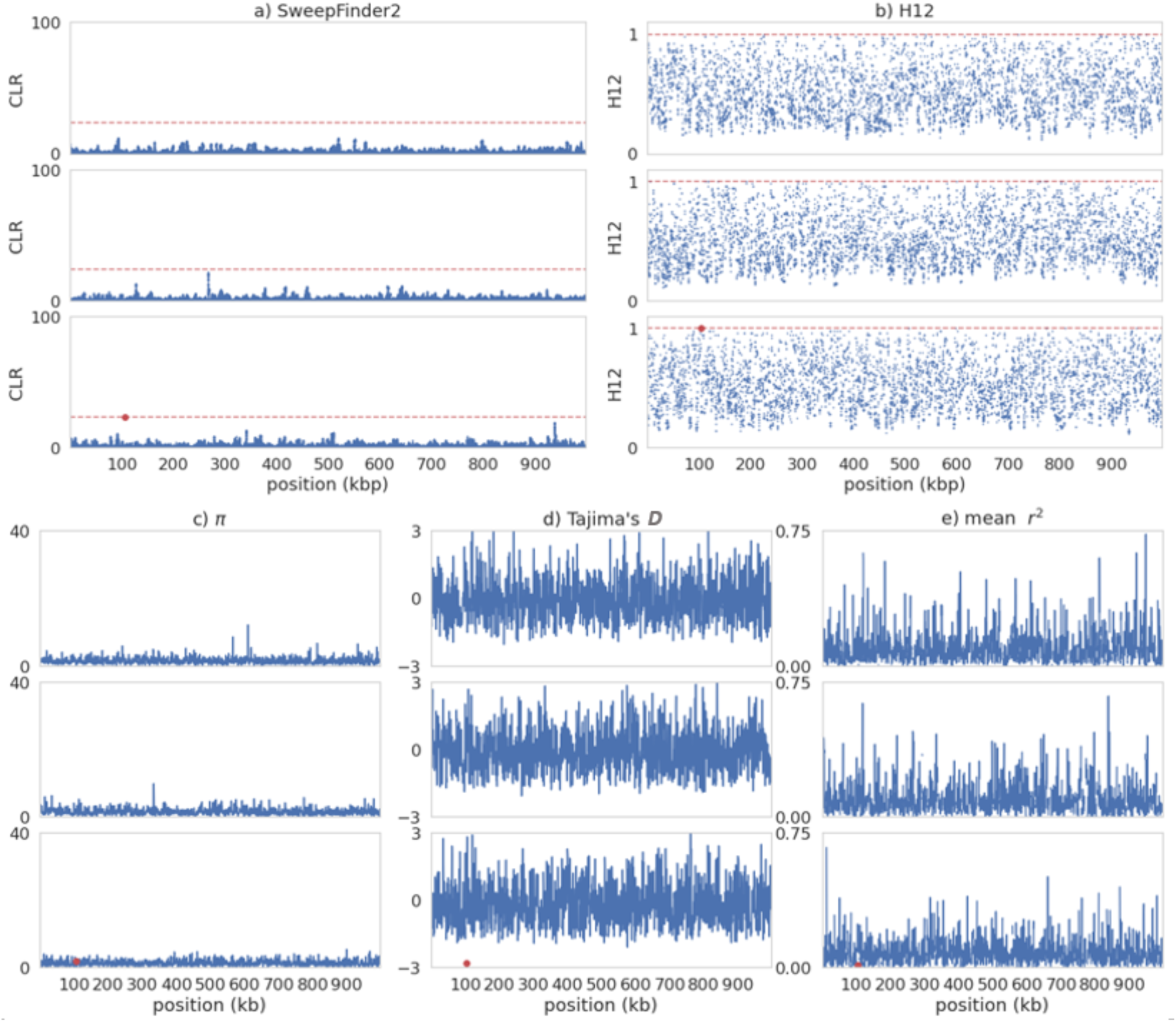
Sweep inference and summary statistics for a single simulation replicate of recurrent selective sweeps for different values of 2*N_e_s* **under instantaneous population contraction (such that *N_current_* = *0.1N_ancestral_*) with fixed mutation and recombination rates**. Population size change occurs 1*N* generations before sampling. In each case, 2*N_e_s* values go from lowest (top panel) to highest: 100; 1000; 10,000. For all panels, red data points are the positions of beneficial fixations with the previous 0.5*N* generations prior to sampling. a) Inference results from SweepFinder2. Blue data points are CLR values inferred for each window. The red dashed line is the threshold for sweep detection, determined by the highest CLR value across 200 simulation replicates in which no beneficial mutations are modelled. Inference was performed at each SNP (see Methods section for further details). b) Sweep inference with the H12 statistic. Blue data points are H12 values estimated for each window. As with SweepFinder2, the red dashed line is the threshold for sweep detection. Inference was performed across 1kb windows for each SNP, with the SNP at the center of each window. c-e) Summary statistics across the simulated region.

**Figure S11:**
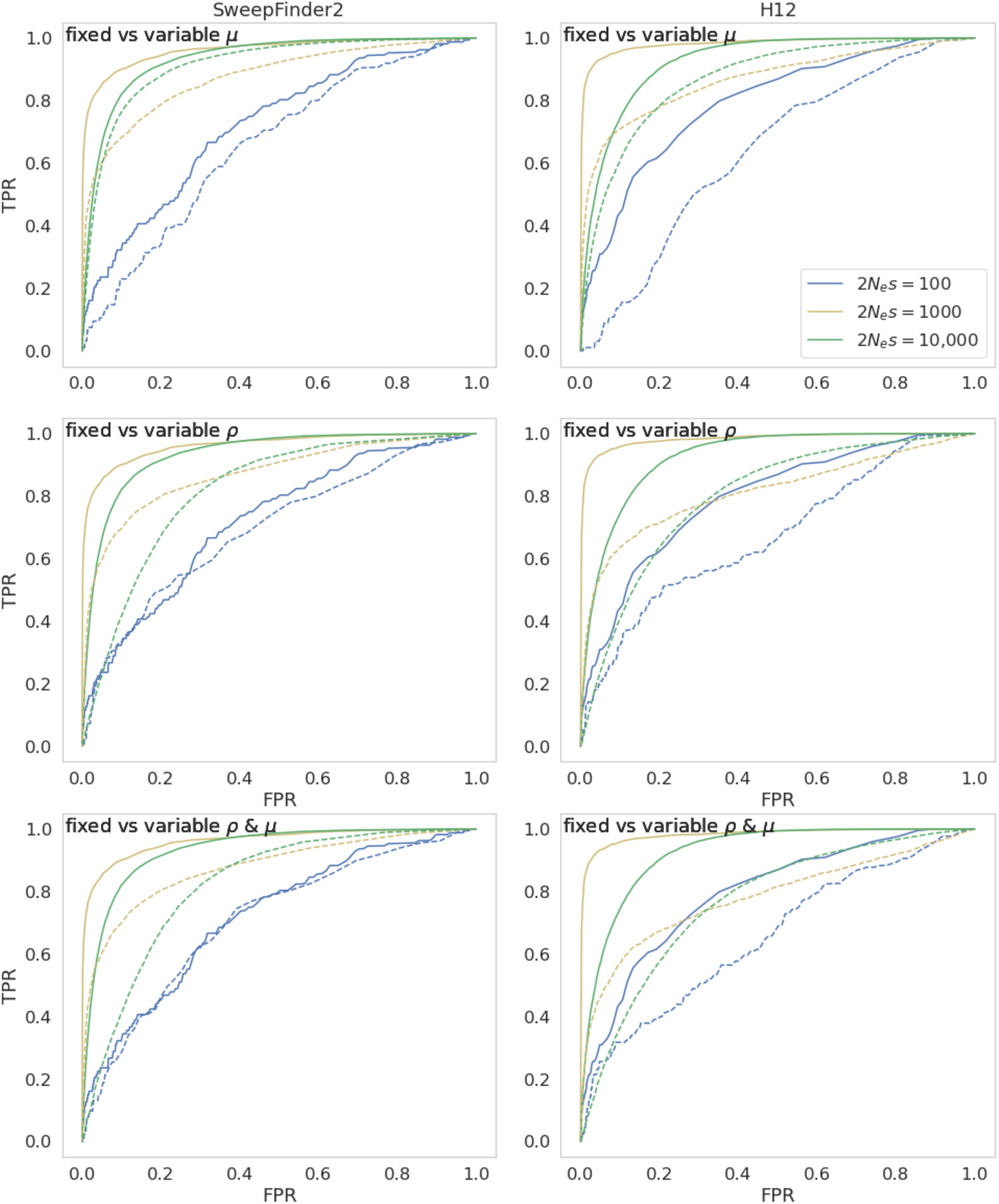
ROC curves comparing sweep inference for fixed and variable recombination and mutation rates under equilibrium demographic conditions, across 200 simulation replicates using SweepFinder2 with the null background SFS (left) and H12 (right), for 1kb windows. Dashed lines indicate variable rates, whilst filled lines indicate fixed rates. For variable rates each 10kb region has a rate drawn from a uniform distribution such that each simulated replicate has the same mean rate as the fixed rate comparison (See Methods for further details).

**Figure S12:**
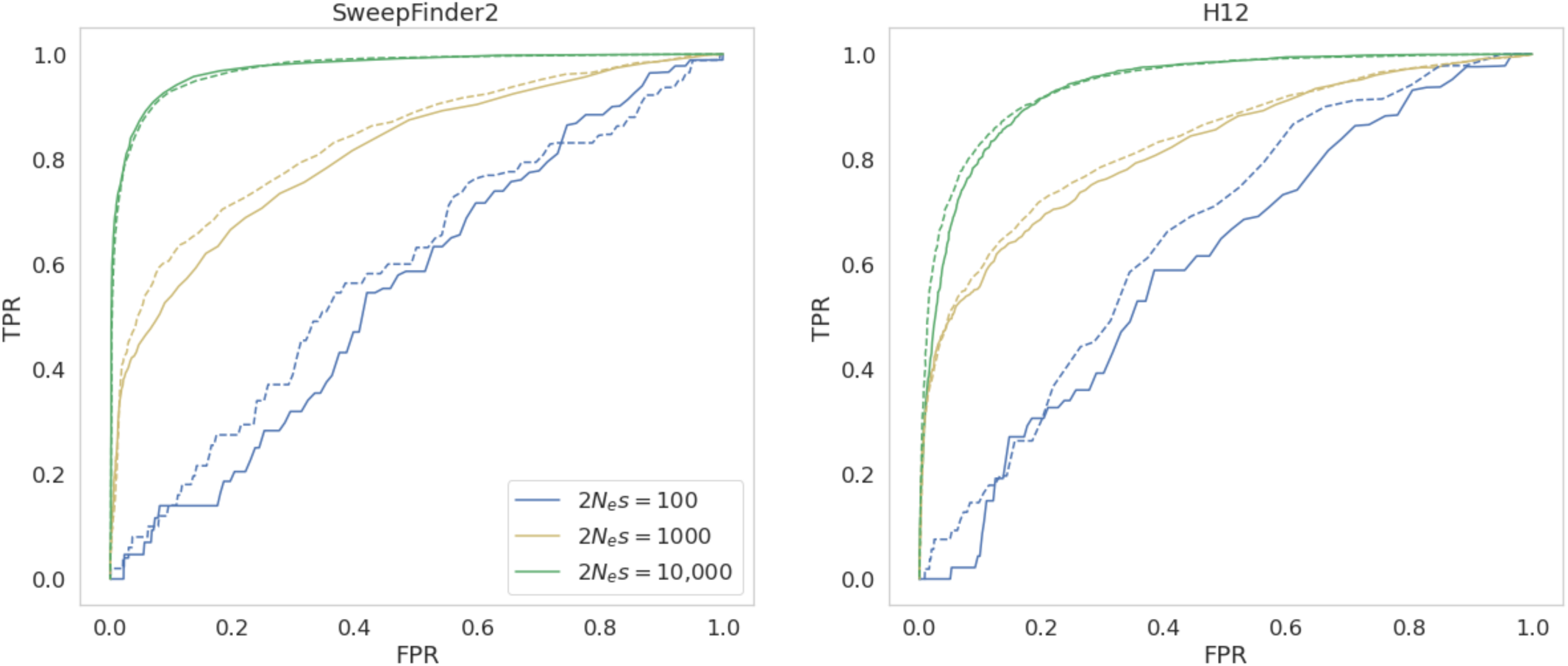
ROC curves for sweep inference for fixed recombination and variable mutation rates under equilibrium population demography with results split into high and low mutation rate bins, for windows of size 10kb. The mean population-scaled mutation rate for the low mutation rate bin is 2.03e-9; for the high mutation rate bin it is 3.65e-9. Dashed lines indicate the high rate bin, and solid lines the low rate bin. For variable rates each 10kb region has a rate drawn from a uniform distribution such that each simulation replicate has the same mean rate as the fixed rate (See Methods for further details).

**Figure S13:**
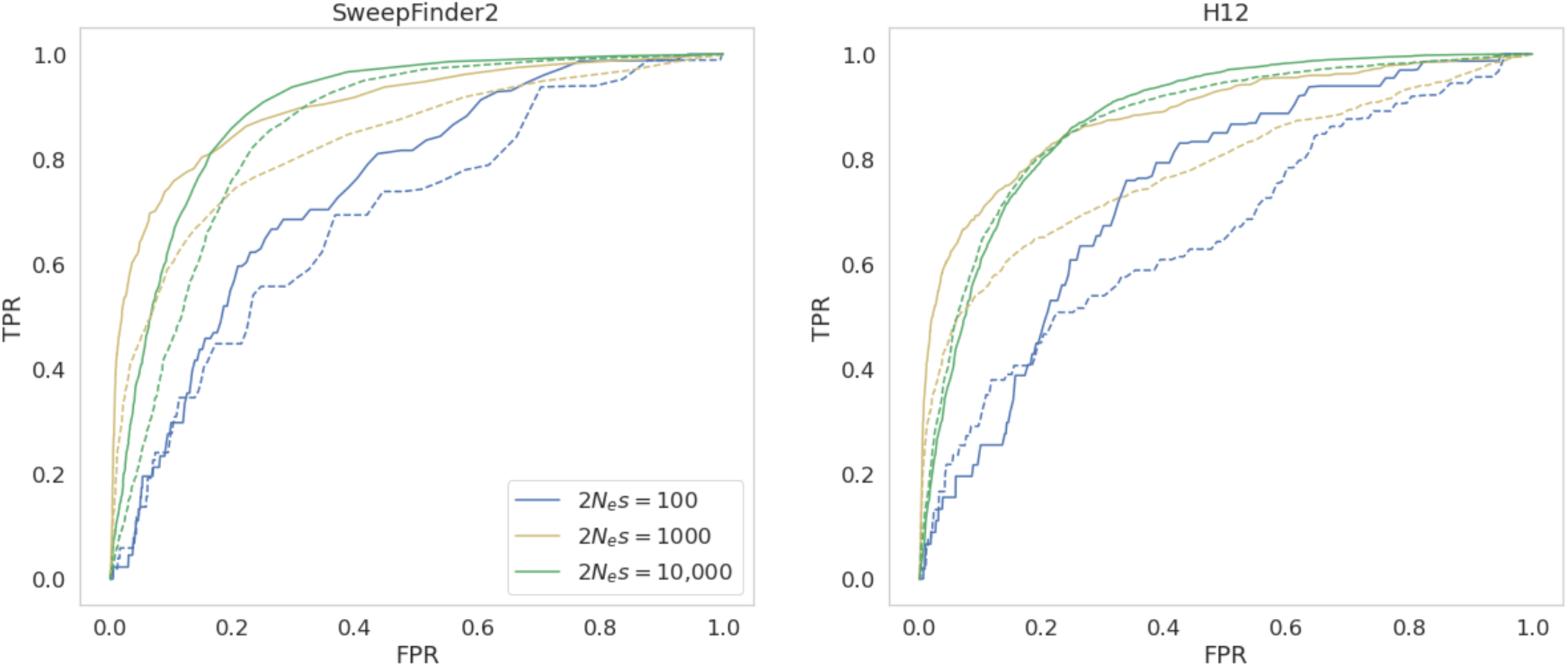
ROC curves for sweep inference for fixed mutation and variable recombination rates under equilibrium population demography with results split into high and low recombination rate bins, for windows of size 10kb. The mean population-scaled mutation rate for the low recombination rate bin is 0.09cM/Mbp; for the high recombination rate bin it is 2.18cM/Mbp. Dashed lines indicate the high rate bin, and solid lines the low rate bin. For variable rates each 10kb region has a rate drawn from a uniform distribution such that each simulation replicate has the same mean rate as the fixed rate (See Methods for further details).

**Figure S14:**
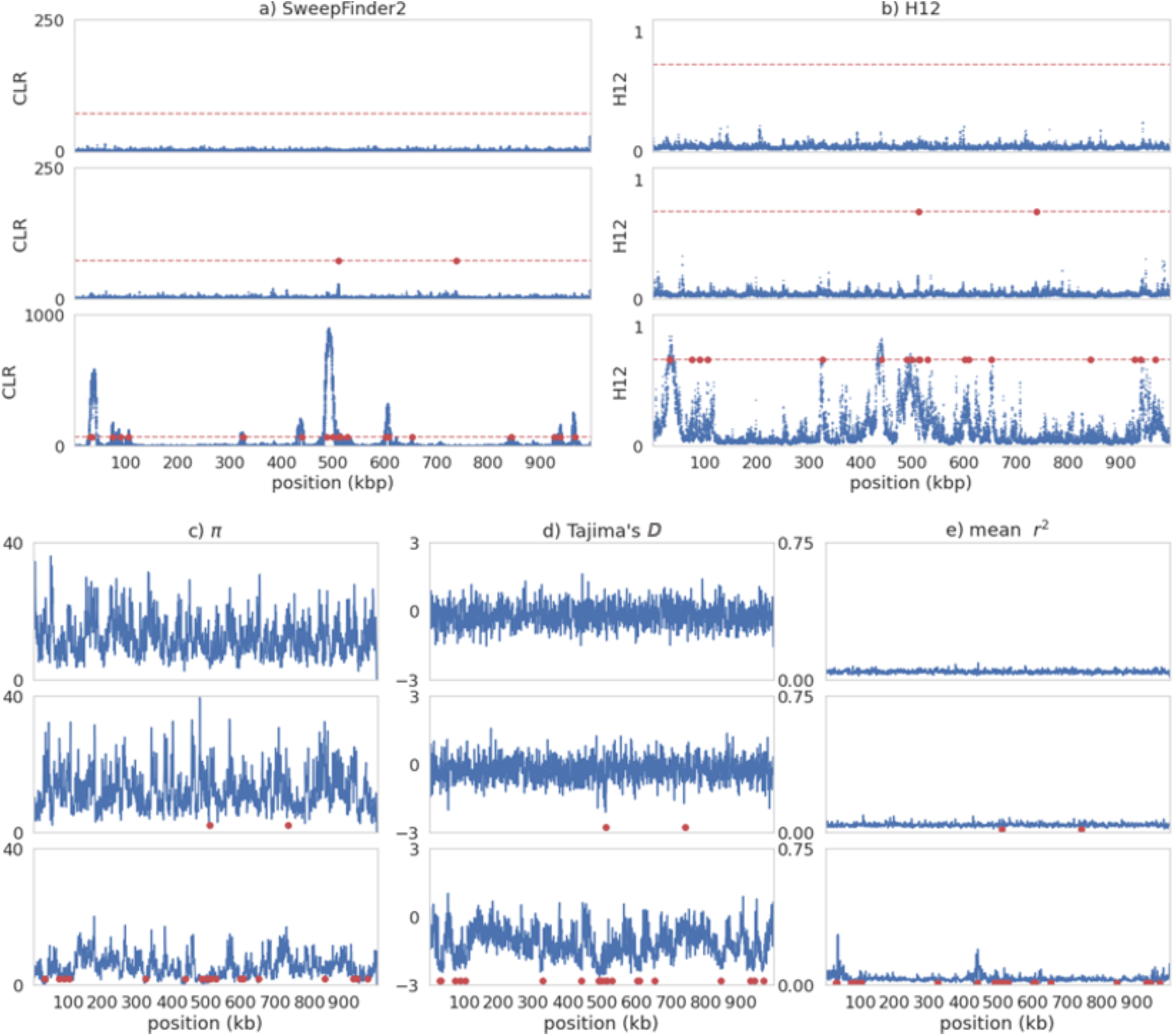
Sweep inference and summary statistics for a single simulation replicate of recurrent selective sweeps for different values of 2*N_e_s* in an **equilibrium population with variable mutation and fixed recombination rates**. In each case, 2*N_e_s* values go from lowest (top panel) to highest: 100; 1000; 10,000. For variable rates each 10kb region has a rate drawn from a uniform distribution such that each simulation replicate has the same mean rate as the fixed rate (See Methods for further details). For all panels, red data points are the positions of beneficial fixations with the previous 0.5*N* generations prior to sampling. a) Inference results from SweepFinder2. Blue data points are CLR values inferred for each window. The red dashed line is the threshold for sweep detection, determined by the highest CLR value across 200 simulation replicates in which no beneficial mutations are modelled. Inference was performed at each SNP (see Methods for further details). b) Sweep inference with the H12 statistic. Blue data points are H12 values estimated for each window. As with SweepFinder2, the red dashed line is the threshold for sweep detection. Inference was performed across 1kb windows for each SNP, with the SNP at the center of each window. c-e) Summary statistics across the simulated region.

**Figure S15:**
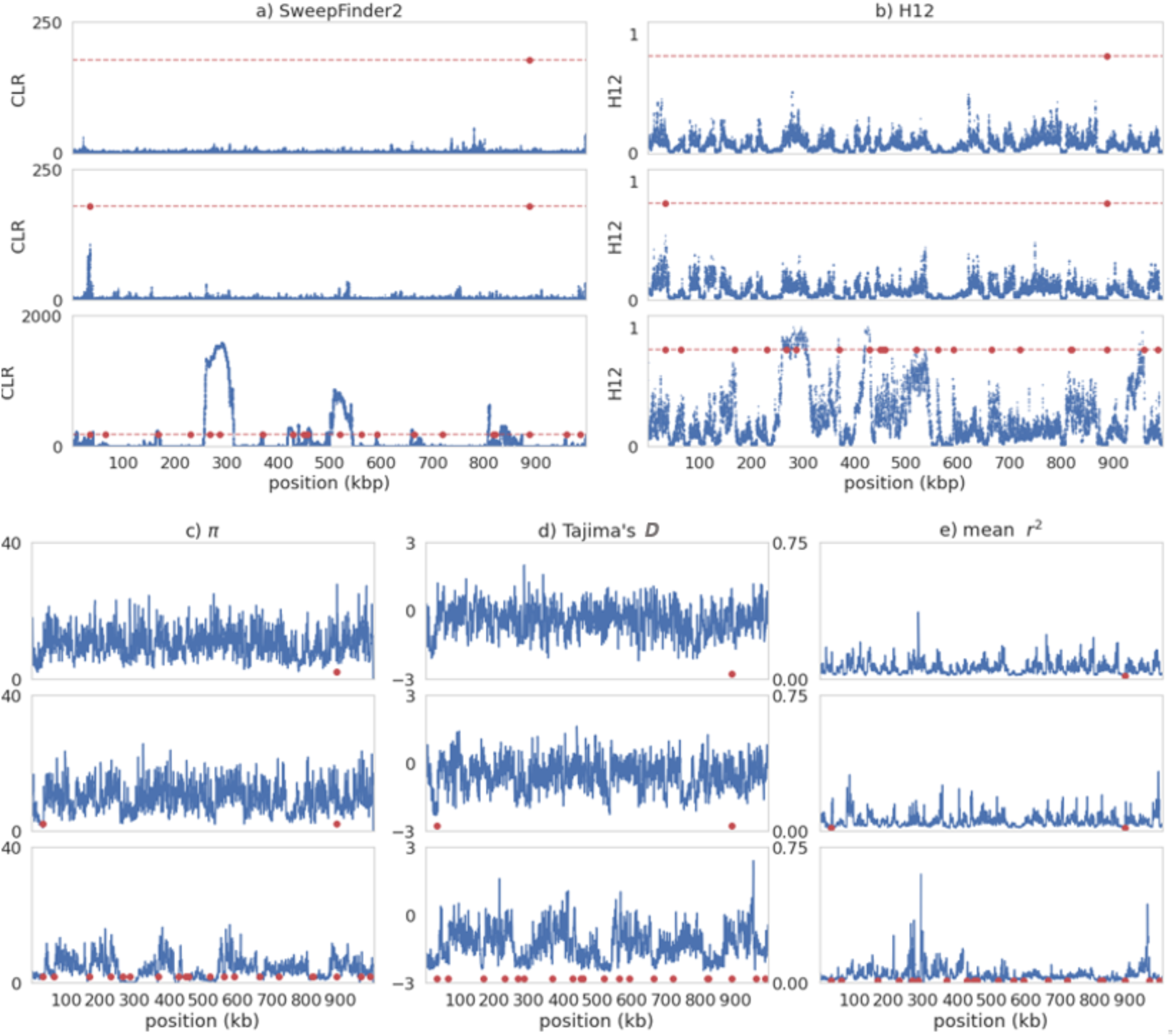
Sweep inference and summary statistics for a single simulation replicate of recurrent selective sweeps for different values of 2*N_e_s* in an **equilibrium population with fixed mutation and variable recombination rates**. In each case, 2*N_e_s* values go from lowest (top panel) to highest: 100; 1000; 10,000. For variable rates each 10kb region has a rate drawn from a uniform distribution such that each simulation replicate has the same mean rate as the fixed rate (See Methods for further details). For all panels, red data points are the positions of beneficial fixations with the previous 0.5*N* generations prior to sampling. a) Inference results from SweepFinder2. Blue data points are CLR values inferred for each window. The red dashed line is the threshold for sweep detection, determined by the highest CLR value across 200 simulation replicates in which no beneficial mutations are modelled. Inference was performed at each SNP (see Methods for further details). b) Sweep inference with the H12 statistic. Blue data points are H12 values estimated for each window. As with SweepFinder2, the red dashed line is the threshold for sweep detection. Inference was performed across 1kb windows for each SNP, with the SNP at the center of each window. c-e) Summary statistics across the simulated region.

**Figure S16:**
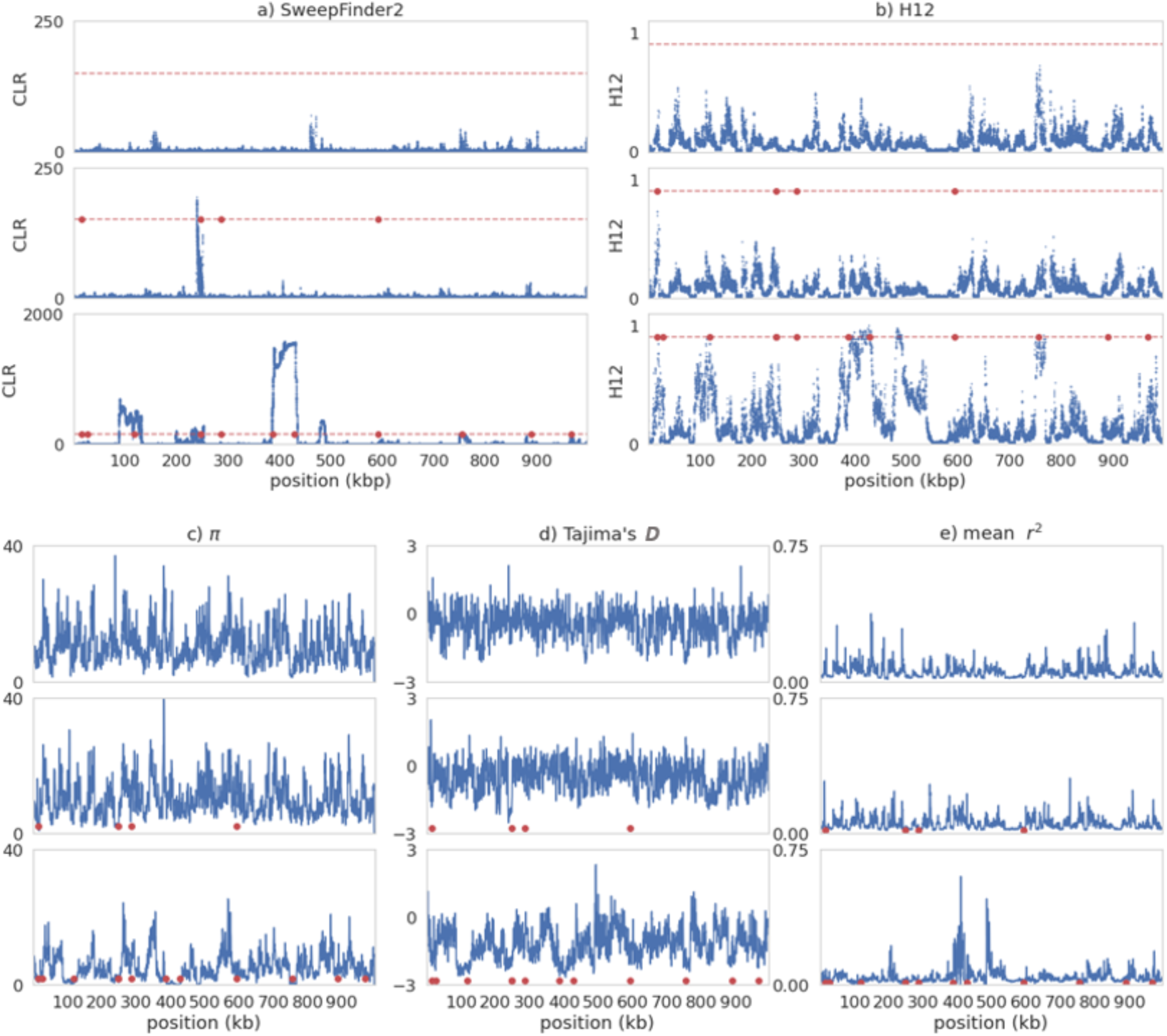
Sweep inference and summary statistics for a single simulation replicate of recurrent selective sweeps for different values of 2*N_e_s* in an **equilibrium population with variable mutation and variable recombination rates**. In each case, 2*N_e_s* values go from lowest (top panel) to highest: 100; 1000; 10,000. For variable rates each 10kb region has a rate drawn from a uniform distribution such that each simulation replicate has the same mean rate as the fixed rate (See Methods for further details). For all panels, red data points are the positions of beneficial fixations with the previous 0.5*N* generations prior to sampling. a) Inference results from SweepFinder2. Blue data points are CLR values inferred for each window. The red dashed line is the threshold for sweep detection, determined by the highest CLR value across 200 simulation replicates in which no beneficial mutations are modelled. Inference was performed at each SNP (see Methods for further details). b) Sweep inference with the H12 statistic. Blue data points are H12 values estimated for each window. As with SweepFinder2, the red dashed line is the threshold for sweep detection. Inference was performed across 1kb windows for each SNP, with the SNP at the center of each window. c-e) Summary statistics across the simulated region.

**Figure S17:**
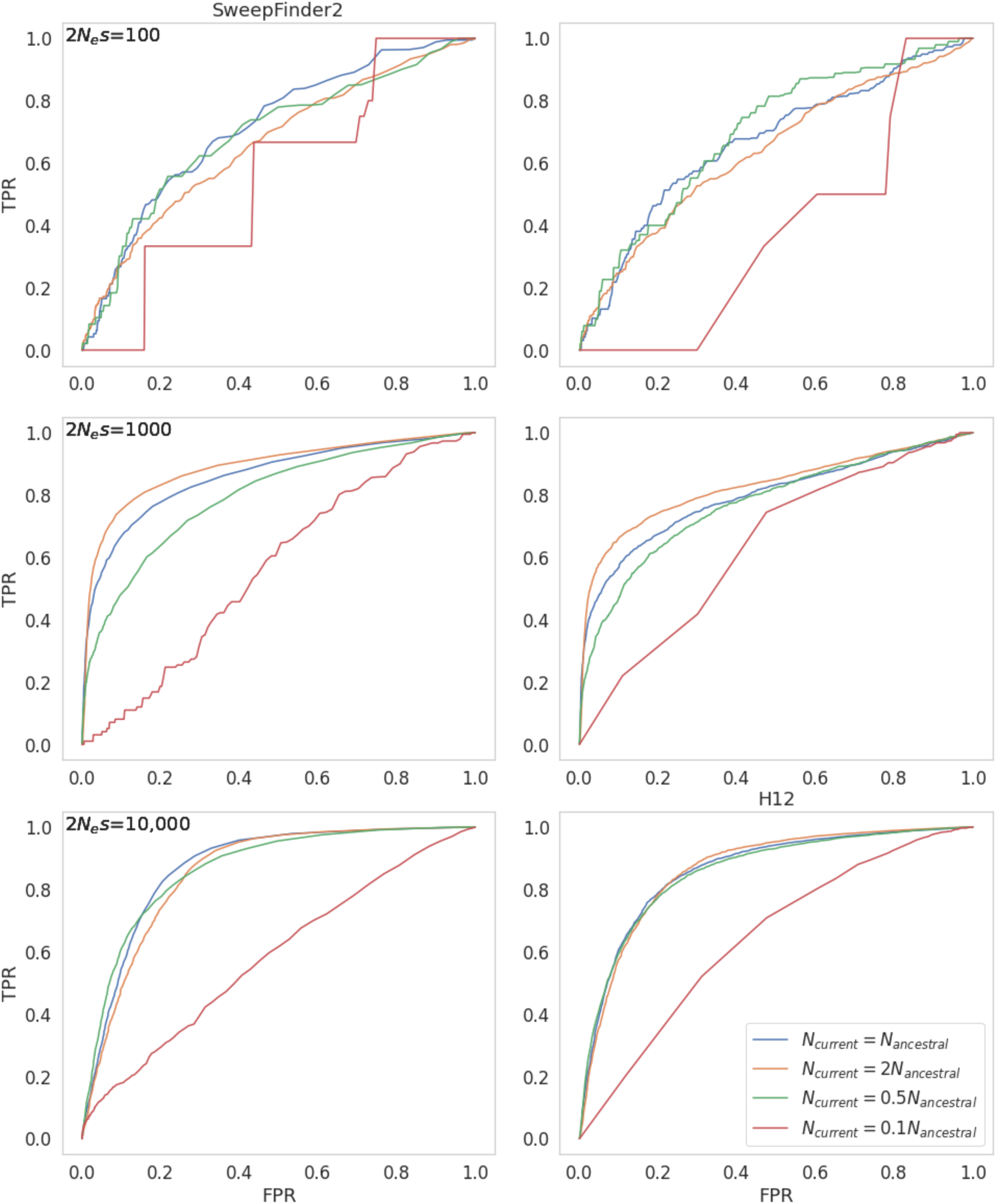
ROC curves for sweep inference for fixed mutation rate and variable recombination rate presented for different demographic histories across 200 simulation replicates using SweepFinder (left) and H12 (right), **for windows of size 10kb**. All population size changes occur instantaneously 1*N* generations before sampling. Each 10kb region has a recombination rate drawn from a uniform distribution such that each simulation replicate has the same mean rate as the fixed rate (See Methods for further details).

**Figure S18:**
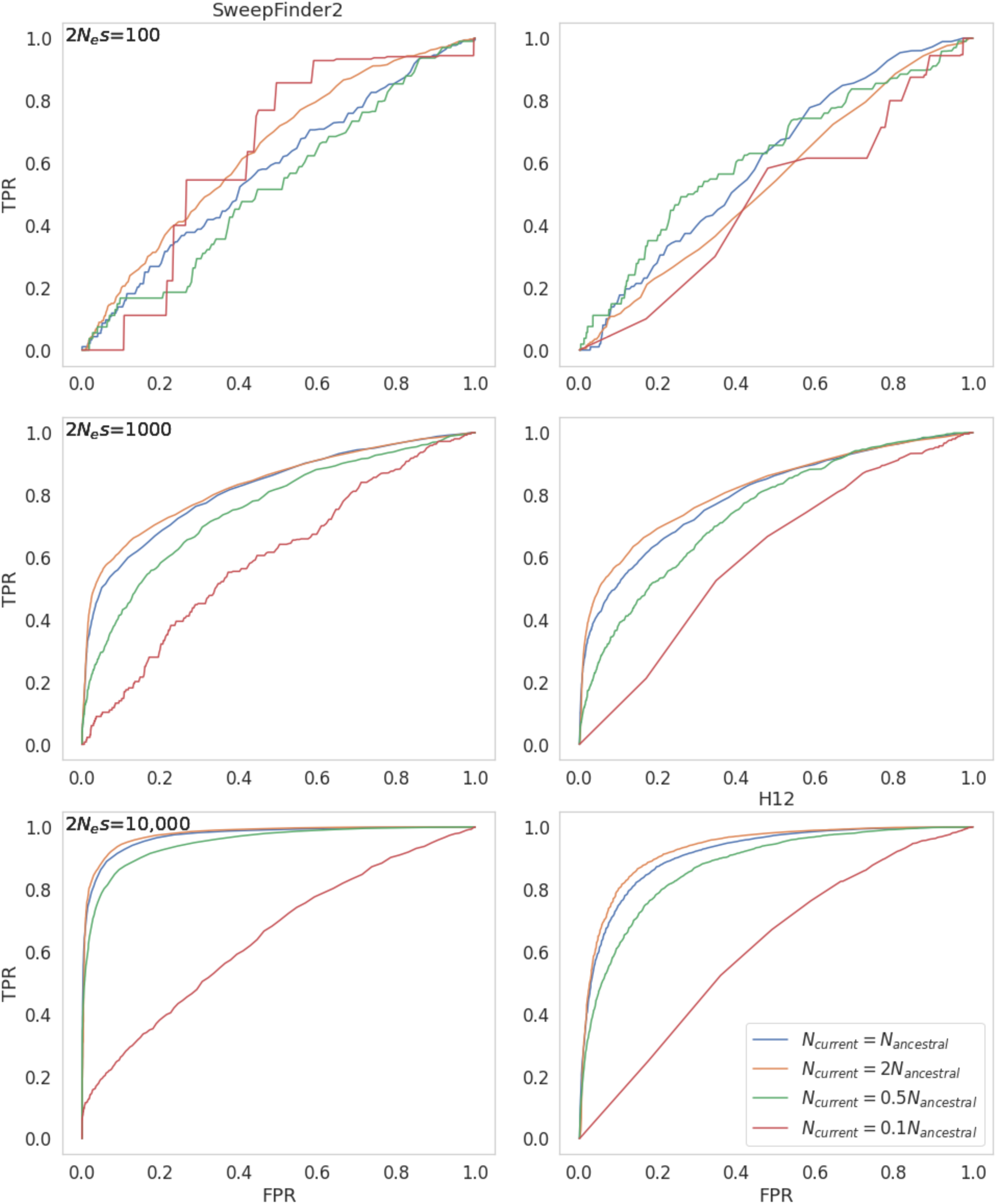
ROC curves for sweep inference for variable mutation rate and fixed recombination rate presented for different demographic histories across 200 simulation replicates using SweepFinder (left) and H12 (right), **for windows of size 10kb**. All population size changes occur instantaneously 1*N* generations before sampling. Each 10kb region has a recombination rate drawn from a uniform distribution such that each simulation replicate has the same mean rate as the fixed rate (See Methods for further details).

**Figure S19:**
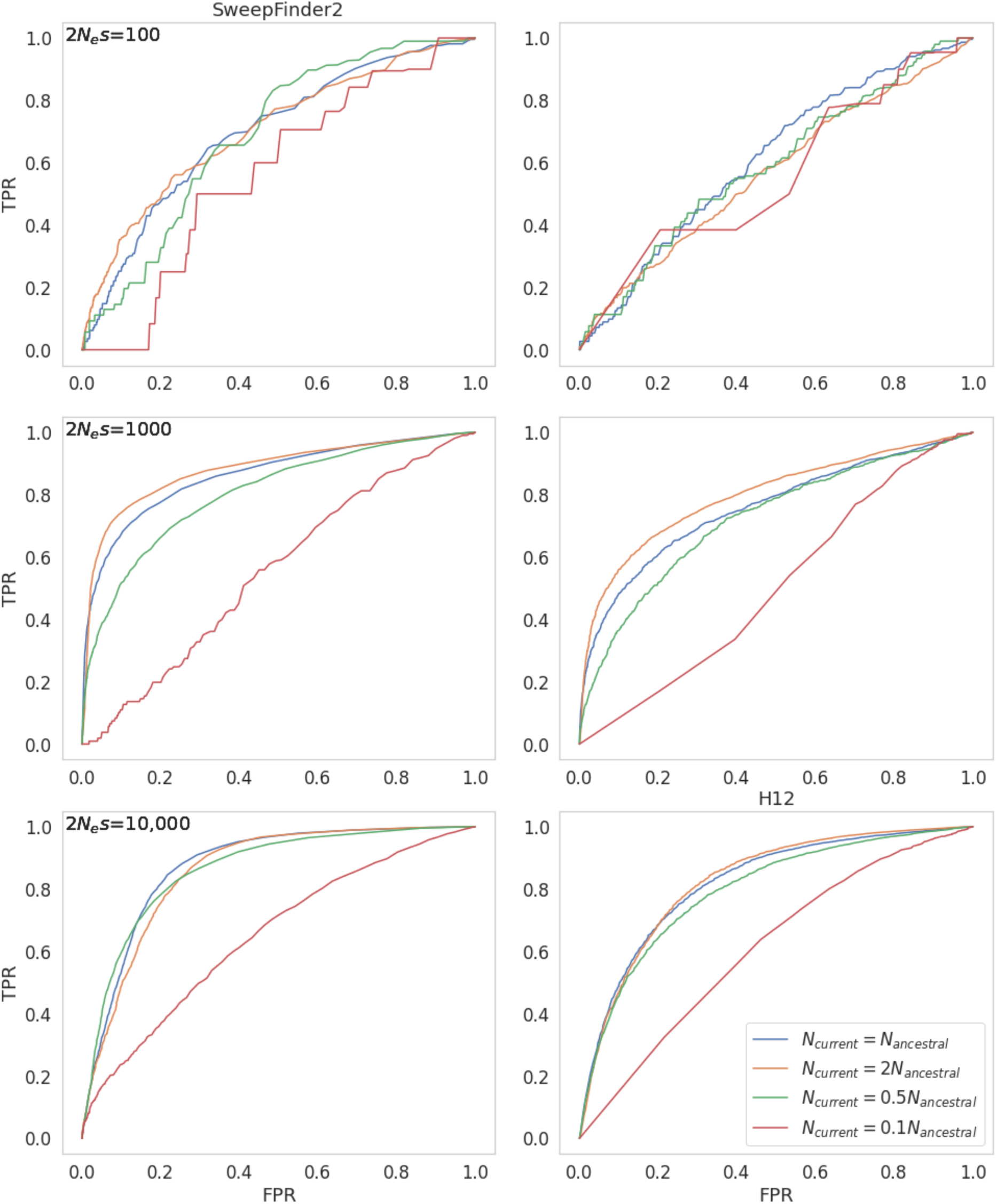
ROC curves for sweep inference for variable mutation rate and variable recombination rate presented for different demographic histories across 200 simulation replicates using SweepFinder (left) and H12 (right), **for windows of size 10kb**. All population size changes occur instantaneously 1*N* generations before sampling. Each 10kb region has a recombination rate drawn from a uniform distribution such that each simulation replicate has the same mean rate as the fixed rate (See Methods for further details).

**Figure S20:**
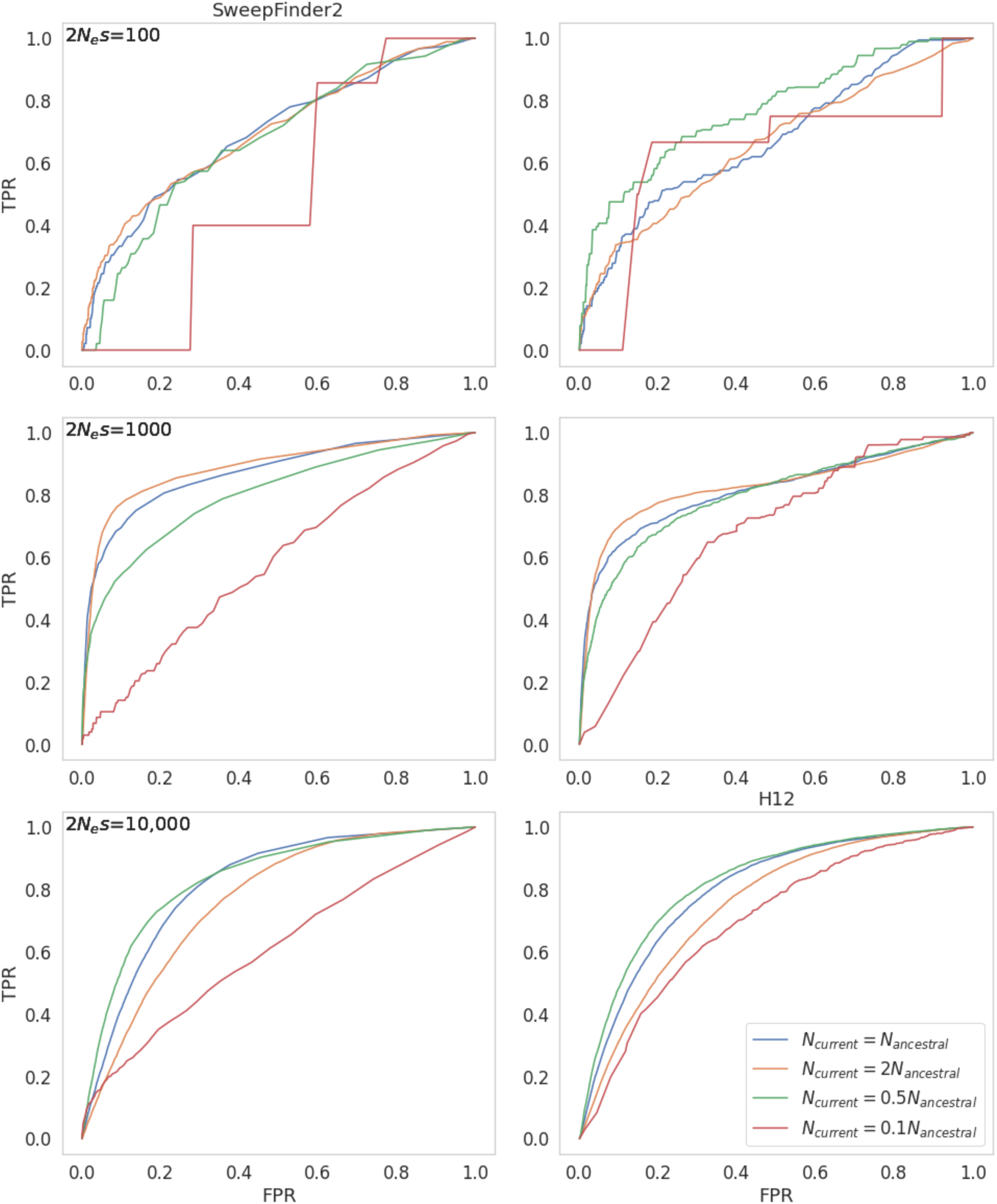
ROC curves for sweep inference for fixed mutation rate and variable recombination rate presented for different demographic histories across 200 simulation replicates using SweepFinder (left) and H12 (right), **for windows of size 1kb**. All population size changes occur instantaneously 1*N* generations before sampling. Each 10kb region has a recombination rate drawn from a uniform distribution such that each simulation replicate has the same mean rate as the fixed rate (See Methods for further details).

**Figure S21:**
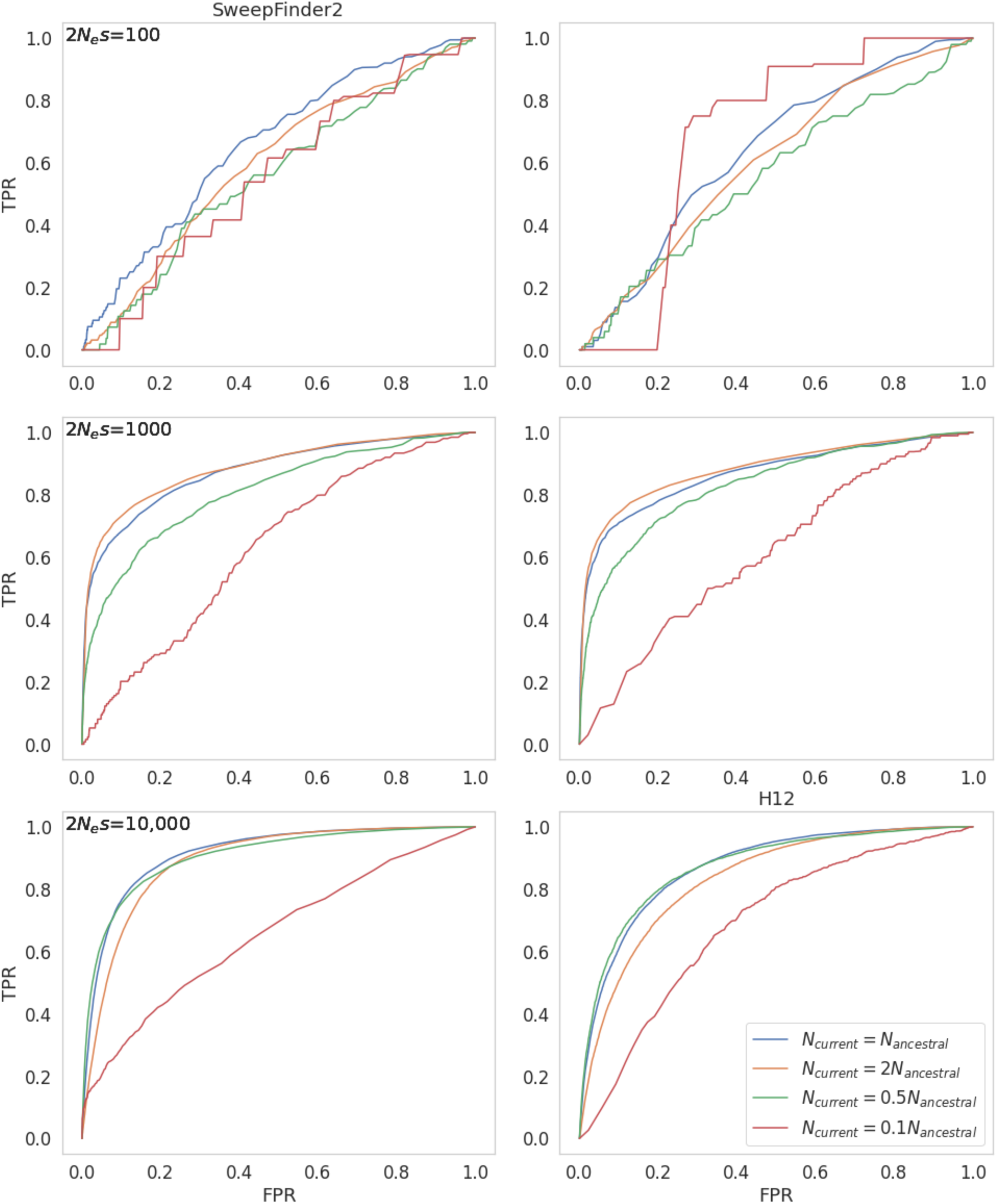
ROC curves for sweep inference for variable mutation rate and fixed recombination rate presented for different demographic histories across 200 simulation replicates using SweepFinder (left) and H12 (right), **for windows of size 1kb**. All population size changes occur instantaneously 1*N* generations before sampling. Each 10kb region has a recombination rate drawn from a uniform distribution such that each simulation replicate has the same mean rate as the fixed rate (See Methods for further details).

**Figure 22:**
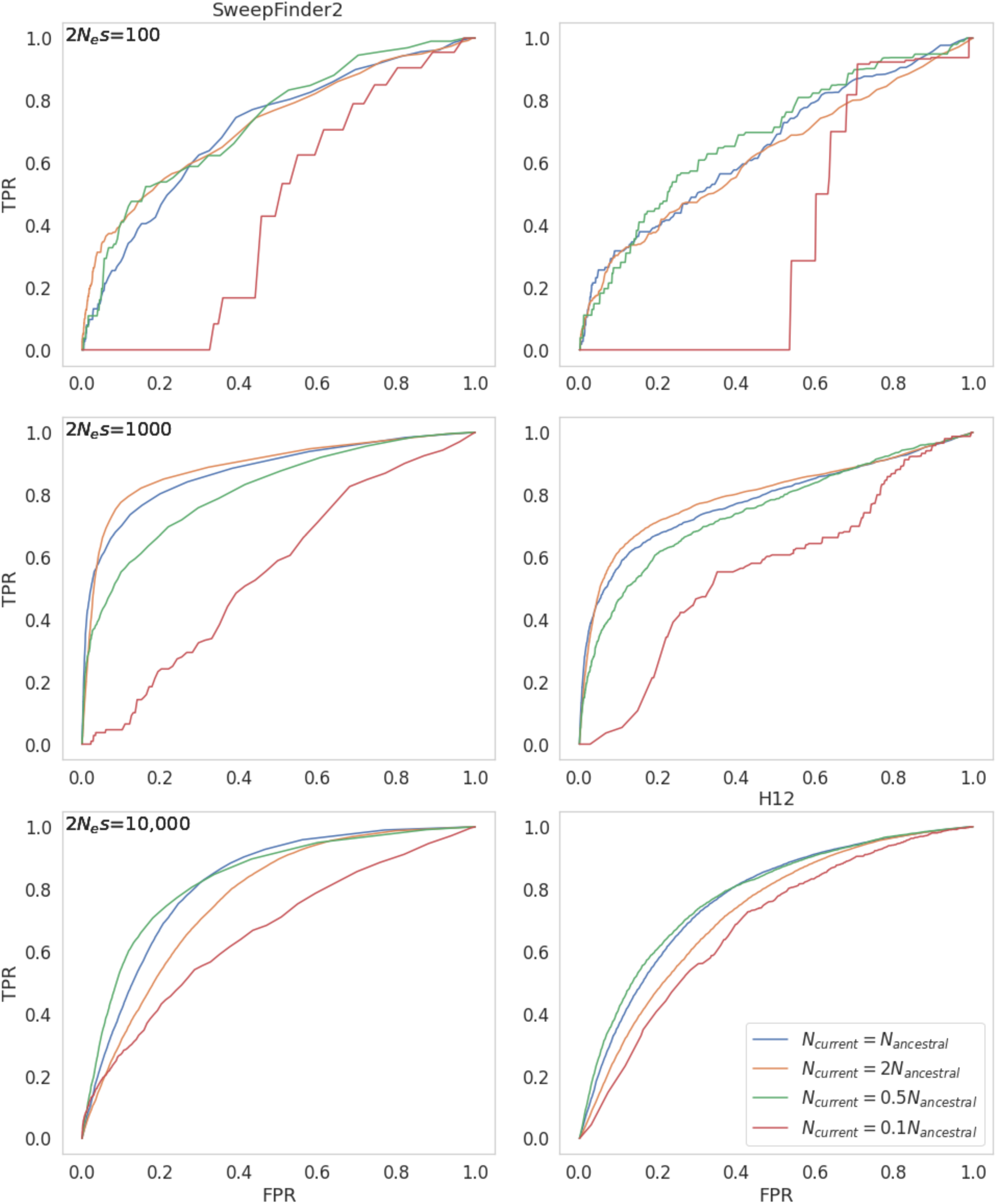
ROC curves for sweep inference for variable mutation rate and variable recombination rate presented for different demographic histories across 200 simulation replicates using SweepFinder (left) and H12 (right), **for windows of size 1kb**. All population size changes occur instantaneously 1*N* generations before sampling. Each 10kb region has a recombination rate drawn from a uniform distribution such that each simulation replicate has the same mean rate as the fixed rate (See Methods for further details).

**Figure S23:**
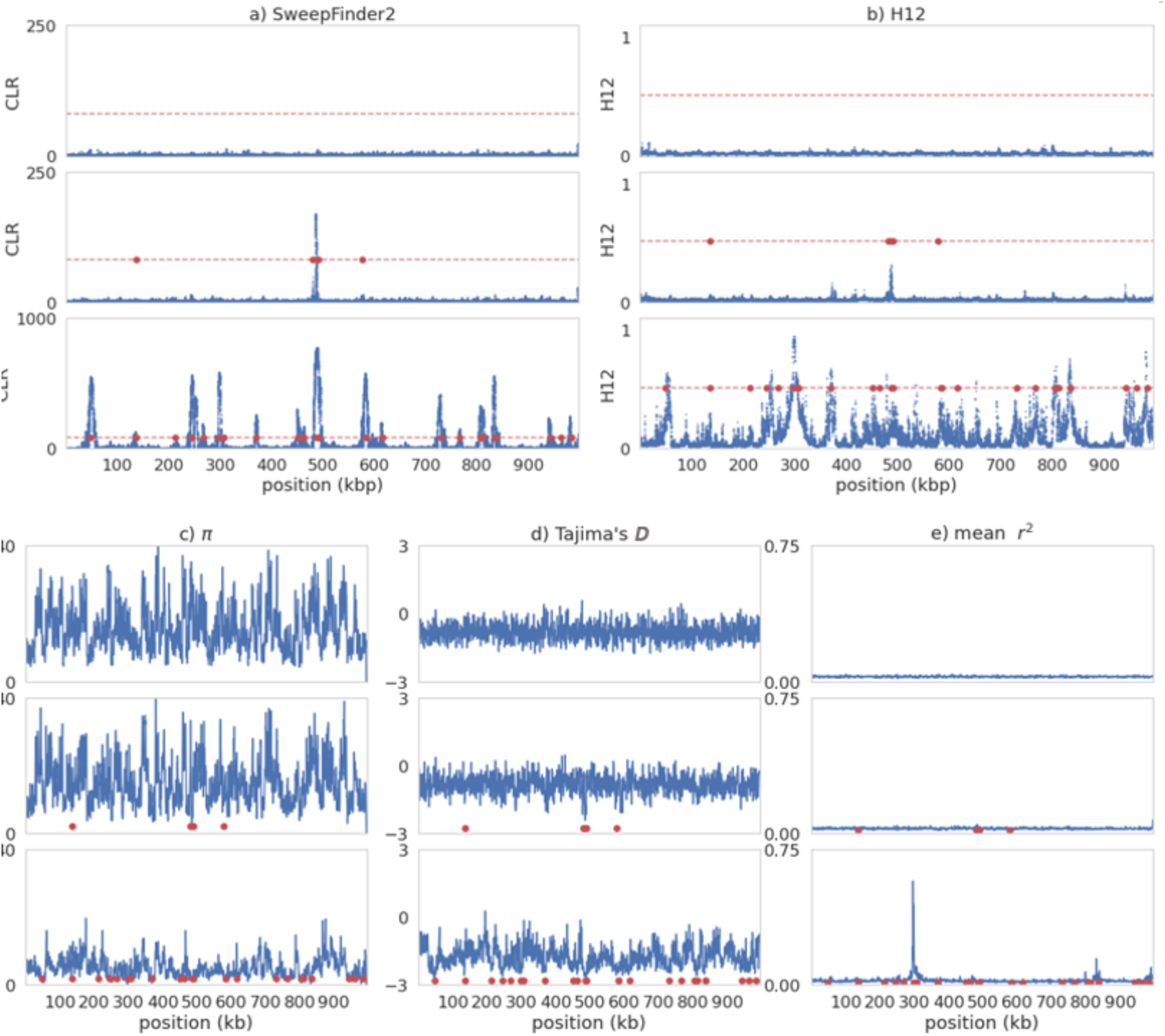
Sweep inference and summary statistics for a single simulation replicate of recurrent selective sweeps for different values of 2*N_e_s* under **instantaneous population expansion (such that *N_current_ = 2N_ancestral_*) with variable mutation and fixed recombination rates**. In each case, 2*N_e_s* values go from lowest (top panel) to highest: 100; 1000; 10,000. For variable rates each 10kb region has a rate drawn from a uniform distribution such that each simulation replicate has the same mean rate as the fixed rate (See Methods for further details). For all panels, red data points are the positions of beneficial fixations with the previous 0.5*N* generations prior to sampling. a) Inference results from SweepFinder2. Blue data points are CLR values inferred for each window. The red dashed line is the threshold for sweep detection, determined by the highest CLR value across 200 simulation replicates in which no beneficial mutations are modelled. Inference was performed at each SNP (see Methods for further details). b) Sweep inference with the H12 statistic. Blue data points are H12 values estimated for each window. As with SweepFinder2, the red dashed line is the threshold for sweep detection. Inference was performed across 1kb windows for each SNP, with the SNP at the center of each window. c-e) Summary statistics across the simulated region.

**Figure S24:**
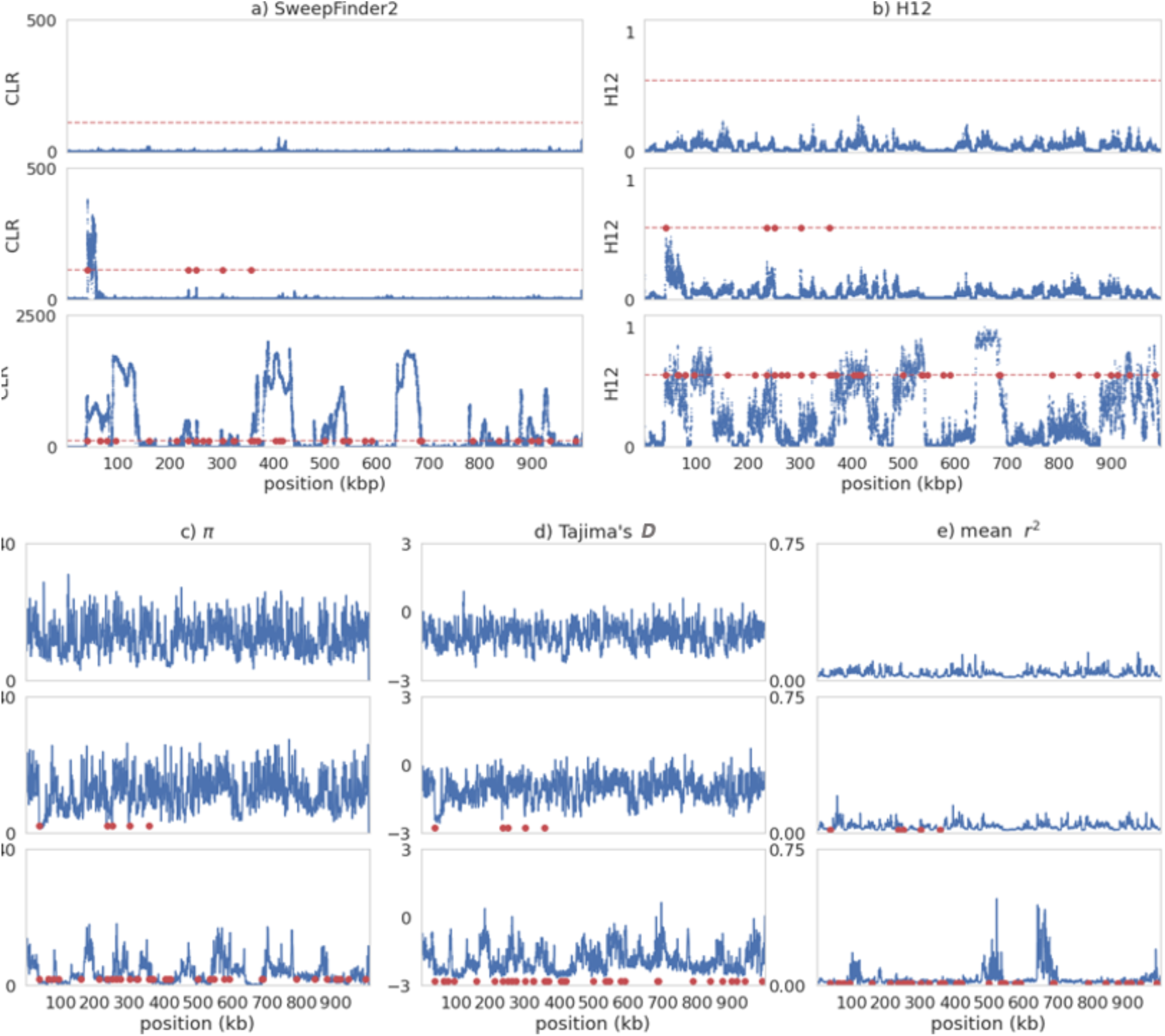
Sweep inference and summary statistics for a single simulation replicate of recurrent selective sweeps for different values of 2*N_e_s* under **instantaneous population expansion (such that *N_current_ = 2N_ancestral_*) with fixed mutation and variable recombination rates**. In each case, 2*N_e_s* values go from lowest (top panel) to highest: 100; 1000; 10,000. For variable rates each 10kb region has a rate drawn from a uniform distribution such that each simulation replicate has the same mean rate as the fixed rate (See Methods for further details). For all panels, red data points are the positions of beneficial fixations with the previous 0.5*N* generations prior to sampling. a) Inference results from SweepFinder2. Blue data points are CLR values inferred for each window. The red dashed line is the threshold for sweep detection, determined by the highest CLR value across 200 simulation replicates in which no beneficial mutations are modelled. Inference was performed at each SNP (see Methods for further details). b) Sweep inference with the H12 statistic. Blue data points are H12 values estimated for each window. As with SweepFinder2, the red dashed line is the threshold for sweep detection. Inference was performed across 1kb windows for each SNP, with the SNP at the center of each window. c-e) Summary statistics across the simulated region.

**Figure S25:**
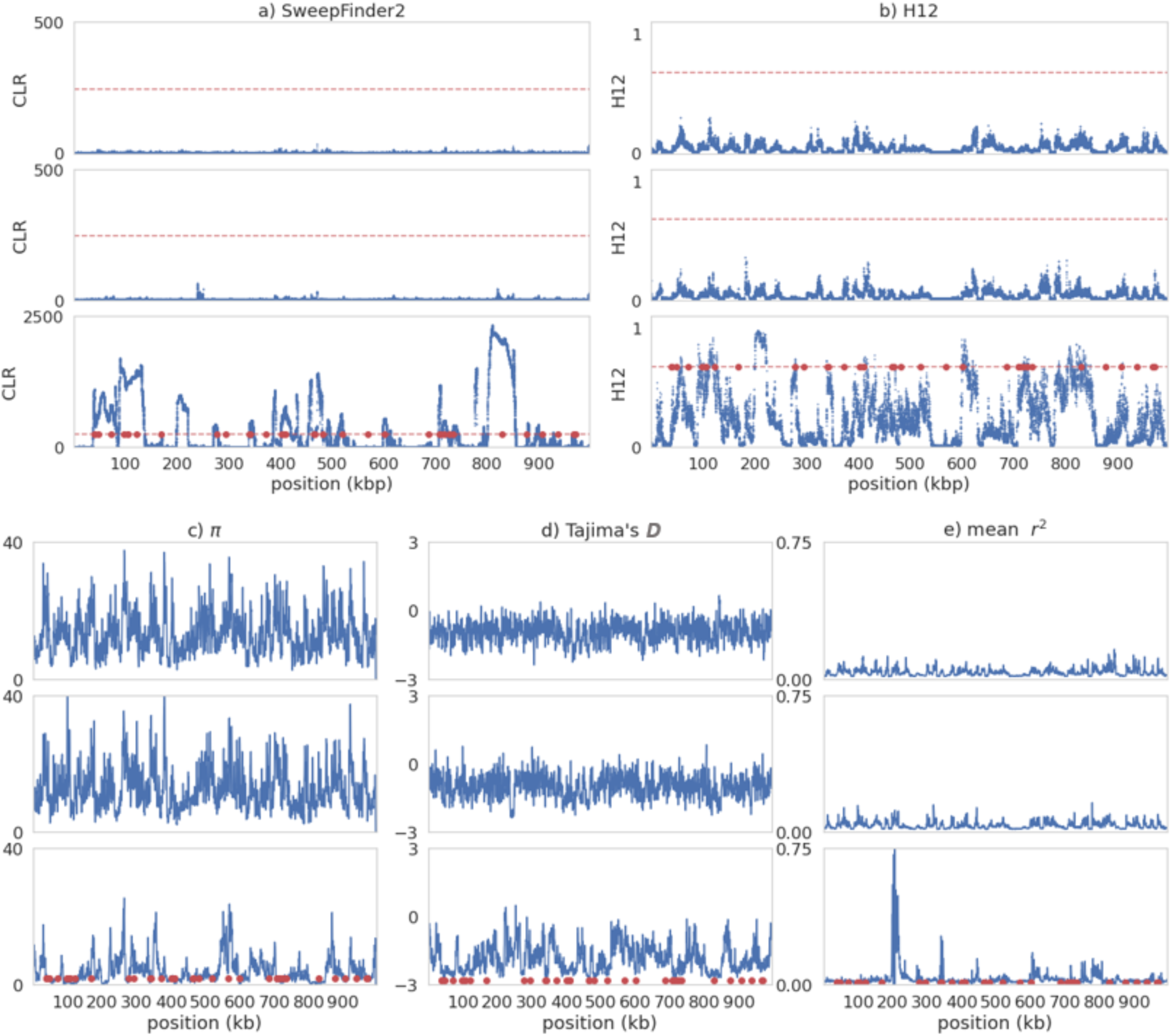
Sweep inference and summary statistics for a single simulation replicate of recurrent selective sweeps for different values of 2*N_e_s* under **instantaneous population expansion (such that *N_current_ = 2N_ancestral_*) with variable mutation and variable recombination rates**. In each case, 2*N_e_s* values go from lowest (top panel) to highest: 100; 1000; 10,000. For variable rates each 10kb region has a rate drawn from a uniform distribution such that each simulation replicate has the same mean rate as the fixed rate (See Methods for further details). For all panels, red data points are the positions of beneficial fixations with the previous 0.5*N* generations prior to sampling. a) Inference results from SweepFinder2. Blue data points are CLR values inferred for each window. The red dashed line is the threshold for sweep detection, determined by the highest CLR value across 200 simulation replicates in which no beneficial mutations are modelled. Inference was performed at each SNP (see Methods for further details). b) Sweep inference with the H12 statistic. Blue data points are H12 values estimated for each window. As with SweepFinder2, the red dashed line is the threshold for sweep detection. Inference was performed across 1kb windows for each SNP, with the SNP at the center of each window. c-e) Summary statistics across the simulated region.

**Figure S26:**
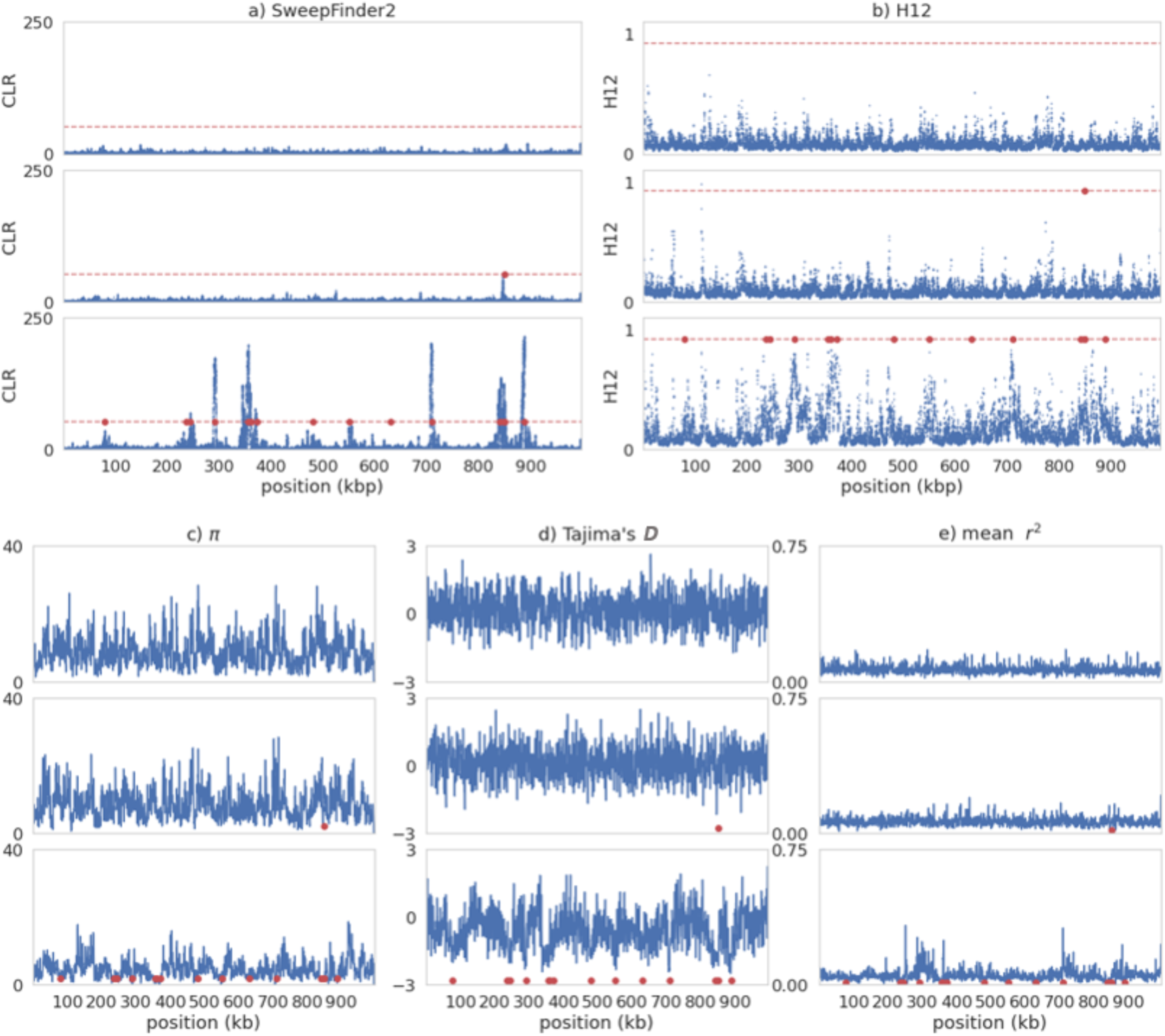
Sweep inference and summary statistics for a single simulation replicate of recurrent selective sweeps for different values of 2*N_e_s* under **instantaneous population contraction (such that *N_current_ = 0.5N_ancestral_*) with variable mutation and fixed recombination rates**. In each case, 2*N_e_s* values go from lowest (top panel) to highest: 100; 1000; 10,000. For variable rates each 10kb region has a rate drawn from a uniform distribution such that each simulation replicate has the same mean rate as the fixed rate (See Methods for further details). For all panels, red data points are the positions of beneficial fixations with the previous 0.5*N* generations prior to sampling. a) Inference results from SweepFinder2. Blue data points are CLR values inferred for each window. The red dashed line is the threshold for sweep detection, determined by the highest CLR value across 200 simulation replicates in which no beneficial mutations are modelled. Inference was performed at each SNP (see Methods for further details). b) Sweep inference with the H12 statistic. Blue data points are H12 values estimated for each window. As with SweepFinder2, the red dashed line is the threshold for sweep detection. Inference was performed across 1kb windows for each SNP, with the SNP at the center of each window. c-e) Summary statistics across the simulated region.

**Figure S27:**
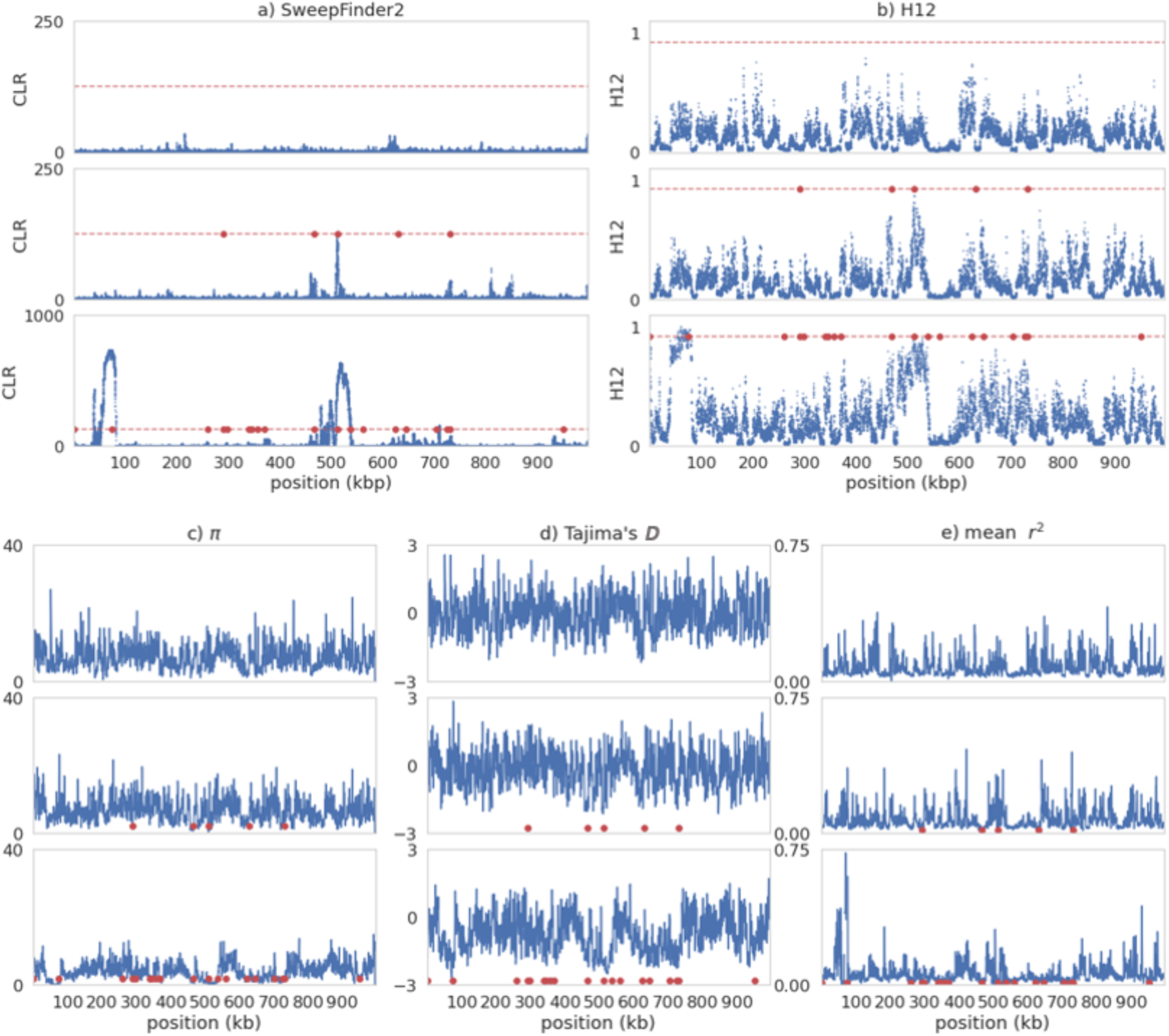
Sweep inference and summary statistics for a single simulation replicate of recurrent selective sweeps for different values of 2*N_e_s* under **instantaneous population contraction (such that *N_current_ = 0.5N_ancestral_*) with fixed mutation and variable recombination rates**. In each case, 2*N_e_s* values go from lowest (top panel) to highest: 100; 1000; 10,000. For variable rates each 10kb region has a rate drawn from a uniform distribution such that each simulation replicate has the same mean rate as the fixed rate (See Methods for further details). For all panels, red data points are the positions of beneficial fixations with the previous 0.5N generations prior to sampling. a) Inference results from SweepFinder2. Blue data points are CLR values inferred for each window. The red dashed line is the threshold for sweep detection, determined by the highest CLR value across 200 simulation replicates in which no beneficial mutations are modelled. Inference was performed at each SNP (see Methods for further details). b) Sweep inference with the H12 statistic. Blue data points are H12 values estimated for each window. As with SweepFinder2, the red dashed line is the threshold for sweep detection. Inference was performed across 1kb windows for each SNP, with the SNP at the center of each window. c-e) Summary statistics across the simulated region.

**Figure S28:**
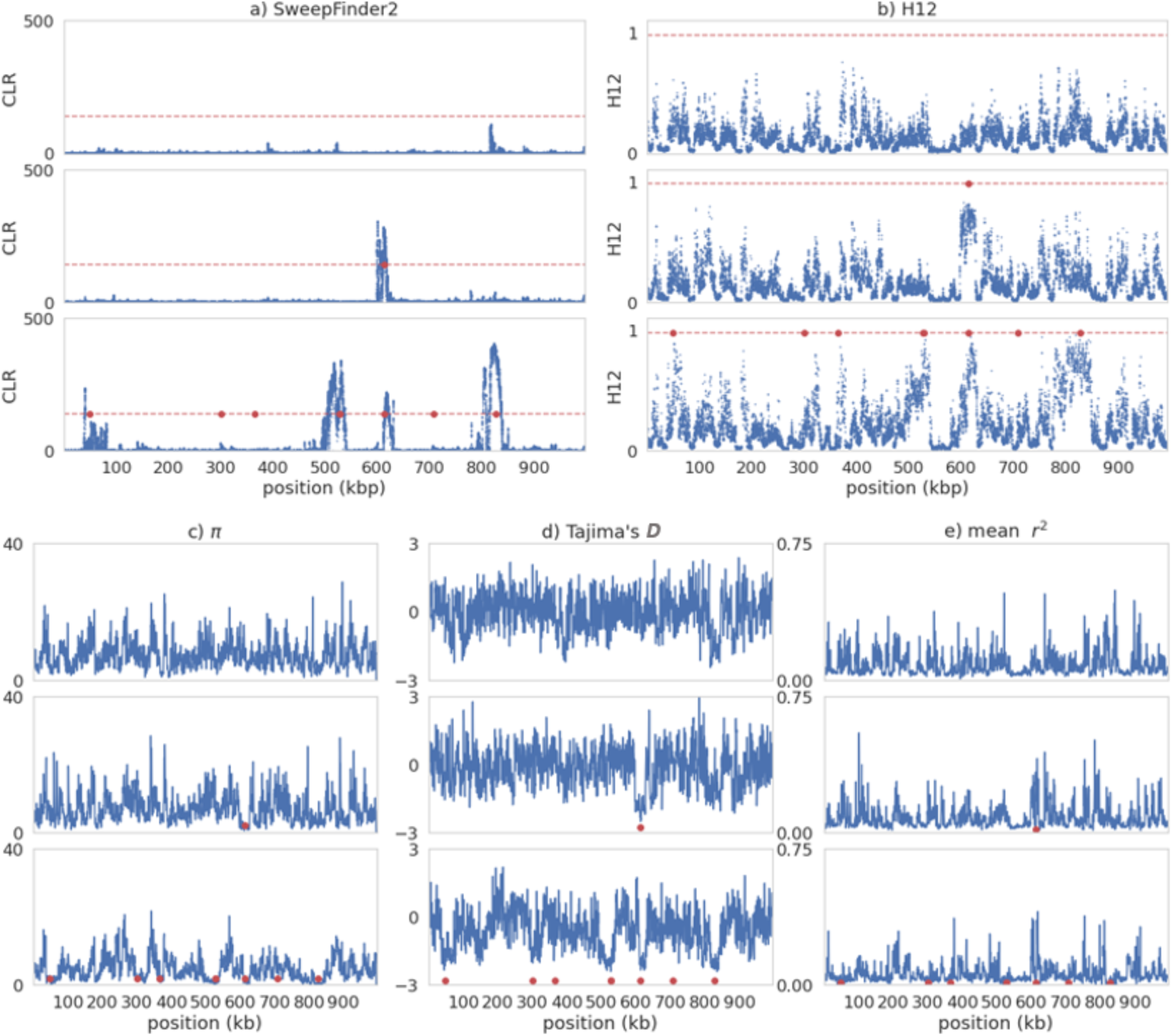
Sweep inference and summary statistics for a single simulation replicate of recurrent selective sweeps for different values of 2*N_e_s* under **instantaneous population contraction (such that *N_current_ = 0.5N_ancestral_*) with variable mutation and variable recombination rates**. In each case, 2*N_e_s* values go from lowest (top panel) to highest: 100; 1000; 10,000. For variable rates each 10kb region has a rate drawn from a uniform distribution such that each simulation replicate has the same mean rate as the fixed rate (See Methods for further details). For all panels, red data points are the positions of beneficial fixations with the previous 0.5N generations prior to sampling. a) Inference results from SweepFinder2. Blue data points are CLR values inferred for each window. The red dashed line is the threshold for sweep detection, determined by the highest CLR value across 200 simulation replicates in which no beneficial mutations are modelled. Inference was performed at each SNP (see Methods for further details). b) Sweep inference with the H12 statistic. Blue data points are H12 values estimated for each window. As with SweepFinder2, the red dashed line is the threshold for sweep detection. Inference was performed across 1kb windows for each SNP, with the SNP at the center of each window. c-e) Summary statistics across the simulated region.

**Figure S29:**
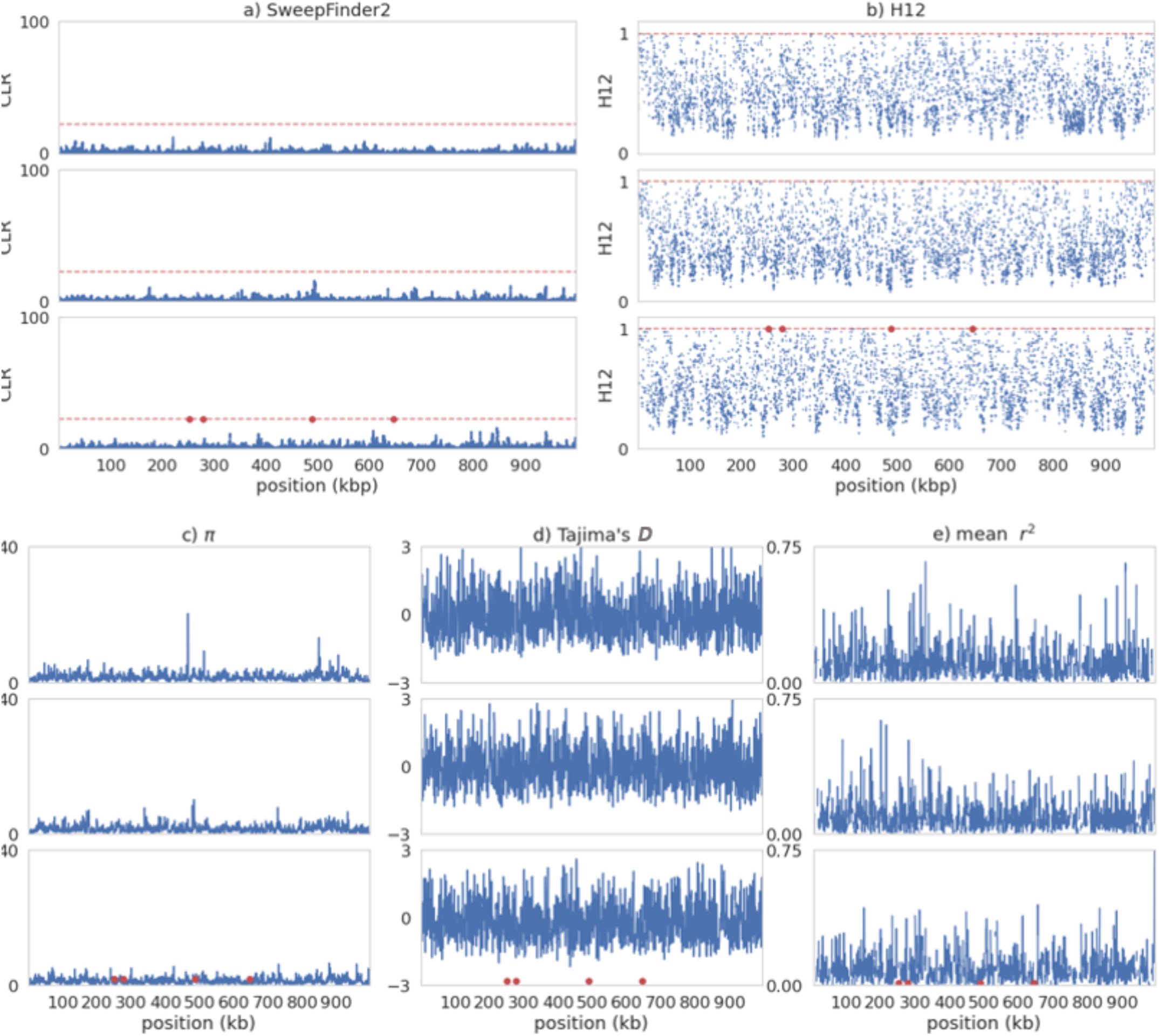
Sweep inference and summary statistics for a single simulation replicate of recurrent selective sweeps for different values of 2*N_e_s* **under instantaneous population contraction (such that *N_current_ = 0.1N_ancestral_*) with variable mutation and fixed recombination rates**. In each case, 2*N_e_s* values go from lowest (top panel) to highest: 100; 1000; 10,000. For variable rates each 10kb region has a rate drawn from a uniform distribution such that each simulation replicate has the same mean rate as the fixed rate (See Methods for further details). For all panels, red data points are the positions of beneficial fixations with the previous 0.5*N* generations prior to sampling. a) Inference results from SweepFinder2. Blue data points are CLR values inferred for each window. The red dashed line is the threshold for sweep detection, determined by the highest CLR value across 200 simulation replicates in which no beneficial mutations are modelled. Inference was performed at each SNP (see Methods for further details). b) Sweep inference with the H12 statistic. Blue data points are H12 values estimated for each window. As with SweepFinder2, the red dashed line is the threshold for sweep detection. Inference was performed across 1kb windows for each SNP, with the SNP at the center of each window. c-e) Summary statistics across the simulated region.

**Figure S30:**
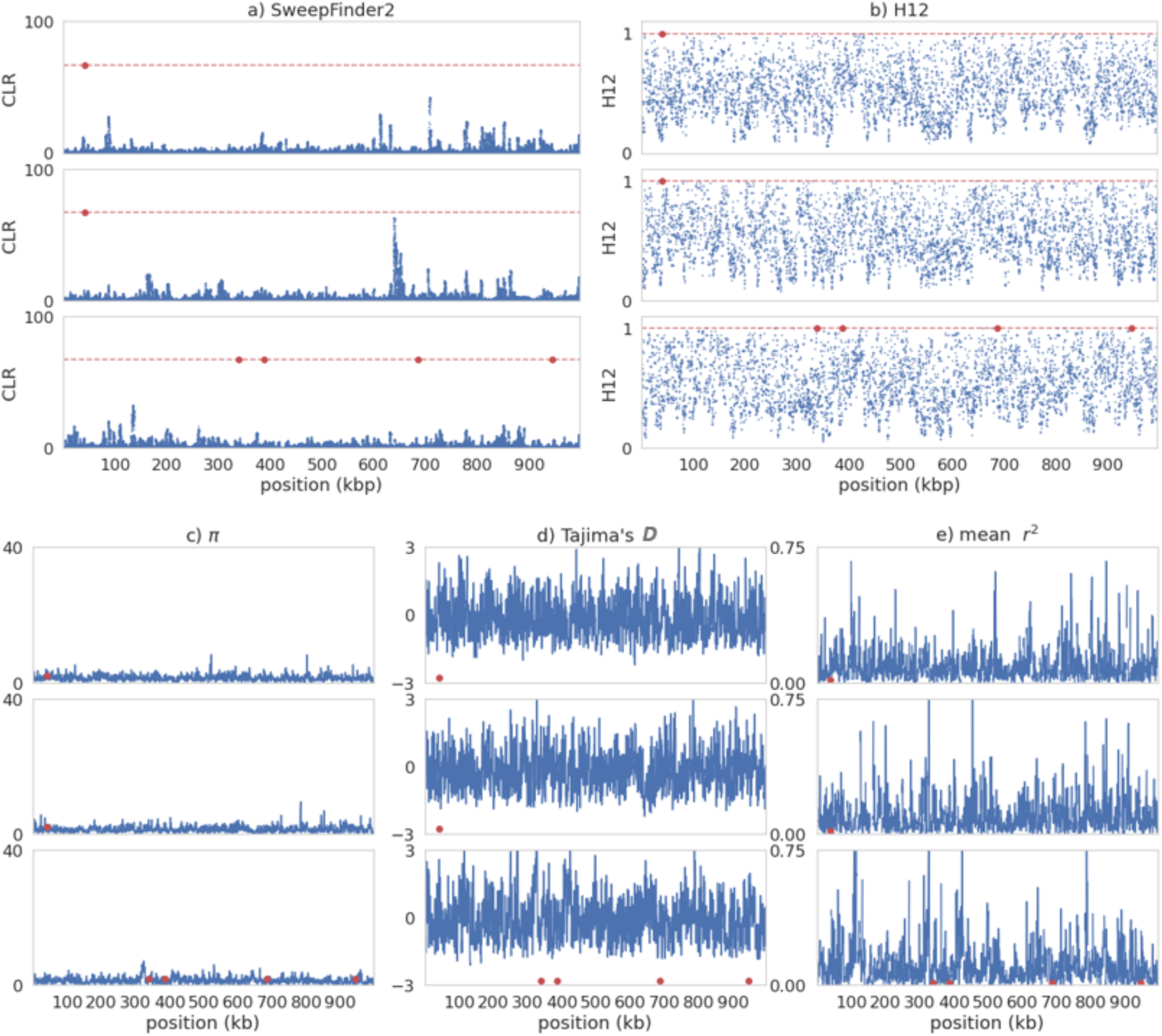
Sweep inference and summary statistics for a single simulation replicate of recurrent selective sweeps for different values of 2*N_e_s* under **instantaneous population contraction (such that *N_current_ = 0.1N_ancestral_*) with fixed mutation and variable recombination rates**. In each case, 2*N_e_s* values go from lowest (top panel) to highest: 100; 1000; 10,000. For variable rates each 10kb region has a rate drawn from a uniform distribution such that each simulation replicate has the same mean rate as the fixed rate (See Methods for further details). For all panels, red data points are the positions of beneficial fixations with the previous 0.5*N* generations prior to sampling. a) Inference results from SweepFinder2. Blue data points are CLR values inferred for each window. The red dashed line is the threshold for sweep detection, determined by the highest CLR value across 200 simulation replicates in which no beneficial mutations are modelled. Inference was performed at each SNP (see Methods for further details). b) Sweep inference with the H12 statistic. Blue data points are H12 values estimated for each window. As with SweepFinder2, the red dashed line is the threshold for sweep detection. Inference was performed across 1kb windows for each SNP, with the SNP at the center of each window. c-e) Summary statistics across the simulated region.

**Figure S31:**
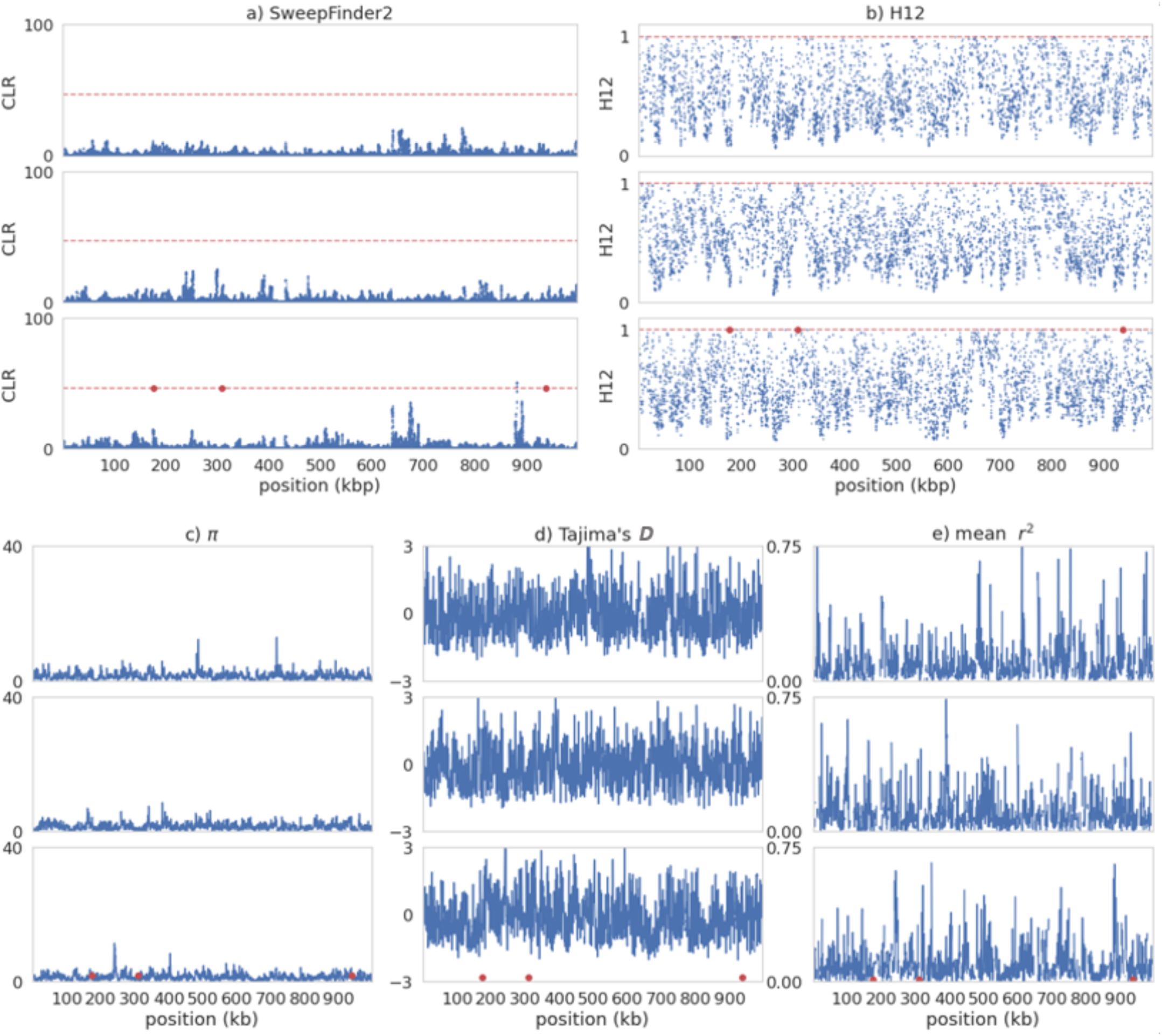
Sweep inference and summary statistics for a single simulation replicate of recurrent selective sweeps for different values of 2*N_e_s* under **instantaneous population contraction (such that *N_current_ = 0.1N_ancestral_*) with variable mutation and variable recombination rates**. In each case, 2*N_e_s* values go from lowest (top panel) to highest: 100; 1000; 10,000. For variable rates each 10kb region has a rate drawn from a uniform distribution such that each simulation replicate has the same mean rate as the fixed rate (See Methods for further details). For all panels, red data points are the positions of beneficial fixations with the previous 0.5*N* generations prior to sampling. a) Inference results from SweepFinder2. Blue data points are CLR values inferred for each window. The red dashed line is the threshold for sweep detection, determined by the highest CLR value across 200 simulation replicates in which no beneficial mutations are modelled. Inference was performed at each SNP (see Methods for further details). b) Sweep inference with the H12 statistic. Blue data points are H12 values estimated for each window. As with SweepFinder2, the red dashed line is the threshold for sweep detection. Inference was performed across 1kb windows for each SNP, with the SNP at the center of each window. c-e) Summary statistics across the simulated region.

**Figure S32:**
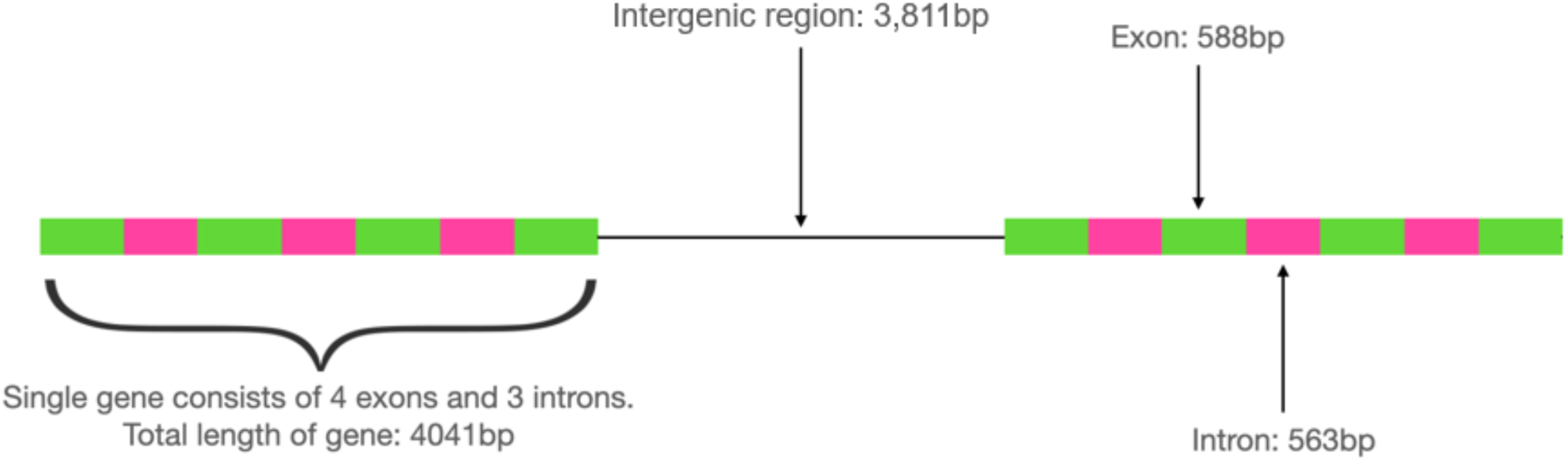
Schematic of genic structure for DFE simulations. Each simulated region is made up of 127 genes, with each pair separated by an intergenic region.

## References

Adams, M. D., Celniker, S. E., Holt, R. A., Evans, C. A., Gocayne, J. D., Amanatides, P. G., Scherer, S. E., Li, P. W., Hoskins, R. A., Galle, R. F., et al. (2000). The genome sequence of *Drosophila melanogaster*. Science, 287(5461), 2185–2195. https://doi.org/10.1126/science.287.5461.2185

Akey, J. M. (2009). Constructing genomic maps of positive selection in humans: Where do we go from here? Genome Research, 19(5), 711–722. https://doi.org/10.1101/gr.086652.108

Akey, J. M., Zhang, G., Zhang, K., Jin, L., & Shriver, M. D. (2002). Interrogating a high-density SNP map for signatures of natural selection. Genome Research, 12(12), 1805–1814. https://doi.org/10.1101/gr.631202

Andolfatto, P. (2005). Adaptive evolution of non-coding DNA in *Drosophila*. Nature, 437(7062), 1149–1152. https://doi.org/10.1038/nature04107

Baer, C. F., Miyamoto, M. M., & Denver, D. R. (2007). Mutation rate variation in multicellular eukaryotes: Causes and consequences. Nature Reviews Genetics, 8(8), 619–631. https://doi.org/10.1038/nrg2158

Bank, C., Ewing, G. B., Ferrer-Admettla, A., Foll, M., & Jensen, J. D. (2014). Thinking too positive? Revisiting current methods of population genetic selection inference. Trends in Genetics, 30(12), 540–546. https://doi.org/10.1016/j.tig.2014.09.010

Barton, N. H. (1998). The effect of hitch-hiking on neutral genealogies. Genetical Research, 72(2), 123–133. https://doi.org/10.1017/S0016672398003462

Barton, N. H. (2000). Genetic hitchhiking. Philosophical Transactions of the Royal Society of London. Series B: Biological Sciences, 355(1403), 1553–1562. https://doi.org/10.1098/rstb.2000.0716

Baudry, E., Viginier, B., & Veuille, M. (2004). Non-African populations of *Drosophila melanogaster* have a unique origin. Molecular Biology and Evolution, 21(8), 1482–1491. https://doi.org/10.1093/molbev/msh089

Bauer DuMont, V., & Aquadro, C. F. (2005). Multiple signatures of positive selection downstream of *Notch* on the X chromosome in *Drosophila melanogaster*. Genetics, 171(2), 639–653. https://doi.org/10.1534/genetics.104.038851

Begun, D. J., & Aquadro, C. F. (1992). Levels of naturally occurring DNA polymorphism correlate with recombination rates in *D. melanogaster*. Nature, 356(6369), 519–520. https://doi.org/10.1038/356519a0

Berry, A. J., Ajioka, J. W., & Kreitman, M. (1991). Lack of polymorphism on the *Drosophila* fourth chromosome resulting from selection. Genetics, 129(4), 1111–1117.

Birky, C. W., & Walsh, J. B. (1988). Effects of linkage on rates of molecular evolution. Proceedings of the National Academy of Sciences, 85(17), 6414–6418. https://doi.org/10.1073/pnas.85.17.6414

Braverman, J. M., Hudson, R. R., Kaplan, N. L., Langley, C. H., & Stephan, W. (1995). The hitchhiking effect on the site frequency spectrum of DNA polymorphisms. Genetics, 140(2), 783–796. https://doi.org/10.1093/genetics/140.2.783

Campos, J. L., & Charlesworth, B. (2019). The effects on neutral variability of recurrent selective sweeps and background selection. Genetics, 212(1), 287–303. https://doi.org/10.1534/genetics.119.301951

Carlson, J., Locke, A. E., Flickinger, M., Zawistowski, M., Levy, S., Myers, R. M., Boehnke, M., Kang, H. M., Scott, L. J., Li, Z.J., et al. (2018). Extremely rare variants reveal patterns of germline mutation rate heterogeneity in humans. Nature Communications, 9(1), 3753. https://doi.org/10.1038/s41467-018-05936-5

Charlesworth, B. (1996). Background selection and patterns of genetic diversity in *Drosophila melanogaster*. Genetical Research, 68(2), 131–149. https://doi.org/10.1017/S0016672300034029

Charlesworth, B., & Jensen, J. D. (2021). Effects of selection at linked sites on patterns of genetic variability. Annual Review of Ecology, Evolution, and Systematics, 52(1), 177–197. https://doi.org/10.1146/annurev-ecolsys-010621-044528

Charlesworth, B., Morgan, M. T., & Charlesworth, D. (1993). The effect of deleterious mutations on neutral molecular variation. Genetics, 134(4), 1289–1303. https://doi.org/10.1093/genetics/134.4.1289

Comeron, J. M., Ratnappan, R., & Bailin, S. (2012). The many landscapes of recombination in *Drosophila melanogaster*. PLOS Genetics, 8(10), e1002905. https://doi.org/10.1371/journal.pgen.1002905

Cox, A., Ackert-Bicknell, C. L., Dumont, B. L., Ding, Y., Bell, J. T., Brockmann, G. A., Wergedal, J. E., Bult, C., Paigen, B., Flint, J., et al. (2009). A new standard genetic map for the laboratory mouse. Genetics, 182(4), 1335–1344. https://doi.org/10.1534/genetics.109.105486

Crisci, J. L., Poh, Y.-P., Mahajan, S., & Jensen, J. D. (2013). The impact of equilibrium assumptions on tests of selection. Frontiers in Genetics, 4. https://doi.org/10.3389/fgene.2013.00235

Cunningham, F., Allen, J. E., Allen, J., Alvarez-Jarreta, J., Amode, M. R., Armean, I. M., Austine-Orimoloye, O., Azov, A. G., Barnes, I., Bennett, R., et al. (2022). Ensembl 2022. Nucleic Acids Research, 50(D1), D988–D995. https://doi.org/10.1093/nar/gkab1049

Cutter, A. D., & Payseur, B. A. (2013). Genomic signatures of selection at linked sites: Unifying the disparity among species. Nature Reviews. Genetics, 14(4), 262–274. https://doi.org/10.1038/nrg3425

David, J., & Capy, P. (1988). Genetic variation of *Drosophila melanogaster* natural populations. Trends in Genetics, 4(4), 106–111. https://doi.org/10.1016/0168-9525(88)90098-4

DeGiorgio, M., Huber, C. D., Hubisz, M. J., Hellmann, I., & Nielsen, R. (2016). SweepFinder2: Increased sensitivity, robustness and flexibility. Bioinformatics, 32(12), 1895–1897. https://doi.org/10.1093/bioinformatics/btw051

Elyashiv, E., Sattath, S., Hu, T. T., Strutsovsky, A., McVicker, G., Andolfatto, P., Coop, G., & Sella, G. (2016). A genomic map of the effects of linked selection in *Drosophila*. PLOS Genetics, 12(8), e1006130. https://doi.org/10.1371/journal.pgen.1006130

Ewing, G. B., & Jensen, J. D. (2016). The consequences of not accounting for background selection in demographic inference. Molecular Ecology, 25(1), 135–141. https://doi.org/10.1111/mec.13390

Excoffier, L., Dupanloup, I., Huerta-Sánchez, E., Sousa, V. C., & Foll, M. (2013). Robust demographic inference from genomic and SNP data. PLOS Genetics, 9(10), e1003905. https://doi.org/10.1371/journal.pgen.1003905

Fay, J. C., & Wu, C.-I. (2000). Hitchhiking under positive Darwinian selection. Genetics, 155(3), 1405–1413. https://doi.org/10.1093/genetics/155.3.1405

Garud, N. R., Messer, P. W., Buzbas, E. O., & Petrov, D. A. (2015). Recent selective sweeps in North American *Drosophila melanogaster* show signatures of soft sweeps. PLOS Genetics, 11(2), e1005004. https://doi.org/10.1371/journal.pgen.1005004

Gillespie, J. H. (2000). Genetic Drift in an infinite population: the pseudohitchhiking model. Genetics, 155(2), 909–919. https://doi.org/10.1093/genetics/155.2.909

Glinka, S., Ometto, L., Mousset, S., Stephan, W., & De Lorenzo, D. (2003). Demography and natural selection have shaped genetic variation in *Drosophila melanogaster*: a multi-locus approach. Genetics, 165(3), 1269–1278. https://doi.org/10.1093/genetics/165.3.1269

Gravel, S., Henn, B. M., Gutenkunst, R. N., Indap, A. R., Marth, G. T., Clark, A. G., Yu, F., Gibbs, R. A., The 1000 Genomes Project, Bustamante, C. D., et al. (2011). Demographic history and rare allele sharing among human populations. Proceedings of the National Academy of Sciences, 108(29), 11983–11988. https://doi.org/10.1073/pnas.1019276108

Gutenkunst, R. N., Hernandez, R. D., Williamson, S. H., & Bustamante, C. D. (2009). Inferring the joint demographic history of multiple populations from multidimensional SNP frequency data. PLOS Genetics, 5(10), e1000695. https://doi.org/10.1371/journal.pgen.1000695

Haller, B. C., & Messer, P. W. (2019). SLiM 3: forward genetic simulations beyond the Wright–Fisher model. Molecular Biology and Evolution, 36(3), 632–637. https://doi.org/10.1093/molbev/msy228

Harr, B., Kauer, M., & Schlötterer, C. (2002). Hitchhiking mapping: A population-based fine-mapping strategy for adaptive mutations in *Drosophila melanogaster*. Proceedings of the National Academy of Sciences, 99(20), 12949–12954. https://doi.org/10.1073/pnas.202336899

Harris, R. B., Sackman, A., & Jensen, J. D. (2018). On the unfounded enthusiasm for soft selective sweeps II: Examining recent evidence from humans, flies, and viruses. PLOS Genetics, 14(12), e1007859. https://doi.org/10.1371/journal.pgen.1007859

Harris, R. B., & Jensen, J. D. (2020). Considering genomic scans for selection as coalescent model choice. Genome Biology and Evolution, 12(6), 871–877. https://doi.org/10.1093/gbe/evaa093

Hermisson, J., & Pennings, P. S. (2005). Soft sweeps. Genetics, 169(4), 2335–2352. https://doi.org/10.1534/genetics.104.036947

Hill, W. G., & Robertson, A. (1966). The effect of linkage on limits to artificial selection. Genetical Research, 8(3), 269–294.

Hodgkinson, A., & Eyre-Walker, A. (2011). Variation in the mutation rate across mammalian genomes. Nature Reviews Genetics, 12(11), 756–766. https://doi.org/10.1038/nrg3098

Howell, A.A., Terbot, J.W., Soni, V., Johri, P., Jensen, J.D., & Pfeifer, S.P. (2023). Developing an appropriate evolutionary baseline model for the study of human cytomegalovirus. Genome Biology and Evolution, 15(4), evad059. https://doi.org/10.1093/gbe/evad059

Huber, C. D., DeGiorgio, M., Hellmann, I., & Nielsen, R. (2016). Detecting recent selective sweeps while controlling for mutation rate and background selection. Molecular Ecology, 25(1), 142–156. https://doi.org/10.1111/mec.13351

Hudson, R. R., & Kaplan, N. L. (1995). Deleterious background selection with recombination. Genetics, 141(4), 1605.

Jensen, J. D. (2009). On reconciling single and recurrent hitchhiking models. Genome Biology and Evolution, 1, 320–324. https://doi.org/10.1093/gbe/evp031

Jensen, J. D. (2014). On the unfounded enthusiasm for soft selective sweeps. Nature Communications, 5(1), 5281. https://doi.org/10.1038/ncomms6281

Jensen, J.D. (2020). Studying population genetic processes in viruses: from drug-resistance evolution to patient infection dynamics. Encyclopedia of Virology, 5, 227–232. https://doi.org/10.1016/B978-0-12-814515-9.00113-2

Jensen, J. D., Kim, Y., DuMont, V. B., Aquadro, C. F., & Bustamante, C. D. (2005). Distinguishing between selective sweeps and demography using DNA polymorphism data. Genetics, 170(3), 1401–1410. https://doi.org/10.1534/genetics.104.038224

Jensen, J.D., & Kowalik, T. F. (2020). A consideration of within-host human cytomegalovirus genetic variation. Proceedings of the National Academy of Sciences, 117(2), 816–817. https://doi.org/10.1073/pnas.191529511

Jensen, J. D., Thornton, K. R., Bustamante, C. D., & Aquadro, C. F. (2007). On the utility of linkage disequilibrium as a statistic for identifying targets of positive selection in nonequilibrium populations. Genetics, 176(4), 2371–2379. https://doi.org/10.1534/genetics.106.069450

Jensen, J. D., Thornton, K. R., & Andolfatto, P. (2008). An approximate Bayesian estimator suggests strong, recurrent selective sweeps in Drosophila. PLOS Genetics, 4(9), e1000198. https://doi.org/10.1371/journal.pgen.1000198

Johri, P., Charlesworth, B., & Jensen, J. D. (2020). Toward an evolutionarily appropriate null model: jointly inferring demography and purifying selection. Genetics, 215(1), 173–192. https://doi.org/10.1534/genetics.119.303002

Johri, P., Charlesworth, B., Howell, E.K., Lynch, M., & Jensen, J.D. (2021a). Revisiting the notion of deleterious sweeps. Genetics, 219(3), iyab094. https://doi.org/10.1093/genetics/iyab094

Johri, P., Riall, K., Becher, H., Excoffier, L., Charlesworth, B., & Jensen, J. D. (2021b). The impact of purifying and background selection on the inference of population history: problems and prospects. Molecular Biology and Evolution, 38(7), 2986–3003. https://doi.org/10.1093/molbev/msab050

Johri, P., Eyre-Walker, A., Gutenkunst, R. N., Lohmueller, K. E., & Jensen, J. D. (2022a). On the prospect of achieving accurate joint estimation of selection with population history. Genome Biology and Evolution, 14(7), evac088. https://doi.org/10.1093/gbe/evac088

Johri, P., Stephan, W., & Jensen, J. D. (2022b). Soft selective sweeps: Addressing new definitions, evaluating competing models, and interpreting empirical outliers. PLOS Genetics, 18(2), e1010022. https://doi.org/10.1371/journal.pgen.1010022

Johri, P., Aquadro, C. F., Beaumont, M., Charlesworth, B., Excoffier, L., Eyre-Walker, A., Keightley, P. D., Lynch, M., McVean, G., Payseur, B. A., Pfeifer, S. P., Stephan, W., & Jensen, J. D. (2022c). Recommendations for improving statistical inference in population genomics. PLOS Biology, 20(5), e3001669. https://doi.org/10.1371/journal.pbio.3001669

Johri, P., Pfeifer, S.P., & Jensen, J.D. (2023). Developing an evolutionary baseline model for humans: jointly inferring purifying selection with population history. Molecular Biology and Evolution, 40(5), msad100. https://doi.org/10/1093/molbev/msad100

Kaplan, N. L., Hudson, R. R., & Langley, C. H. (1989). The ‘hitchhiking effect’ revisited. Genetics, 123(4), 887–899. https://doi.org/10.1093/genetics/123.4.887

Kawakami, T., Smeds, L., Backström, N., Husby, A., Qvarnström, A., Mugal, C. F., Olason, P., & Ellegren, H. (2014). A high-density linkage map enables a second-generation collared flycatcher genome assembly and reveals the patterns of avian recombination rate variation and chromosomal evolution. Molecular Ecology, 23(16), 4035–4058. https://doi.org/10.1111/mec.12810

Keightley, P. D., Ness, R. W., Halligan, D. L., & Haddrill, P. R. (2014). Estimation of the spontaneous mutation rate per nucleotide site in a *Drosophila melanogaster* full-sib family. Genetics, 196(1), 313–320. https://doi.org/10.1534/genetics.113.158758

Kim, Y., & Nielsen, R. (2004). Linkage disequilibrium as a signature of selective sweeps. Genetics, 167(3), 1513–1524. https://doi.org/10.1534/genetics.103.025387

Kim, Y., & Stephan, W. (2000). Joint effects of genetic hitchhiking and background selection on neutral variation. Genetics, 155(3), 1415–1427. https://doi.org/10.1093/genetics/155.3.1415

Kim, Y., & Stephan, W. (2002). Detecting a local signature of genetic hitchhiking along a recombining chromosome. Genetics, 160(2), 765–777. https://doi.org/10.1093/genetics/160.2.765

Kong, A., Gudbjartsson, D. F., Sainz, J., Jonsdottir, G. M., Gudjonsson, S. A., Richardsson, B., Sigurdardottir, S., Barnard, J., Hallbeck, B., Masson, G., et al. (2002). A high-resolution recombination map of the human genome. Nature Genetics, 31(3), 241–247. https://doi.org/10.1038/ng917

Lachaise, D., Cariou, M.L., David, J.R., Lemeunier, F., Tsacas, L., Ashburner, M. (1988). Historical biogeography of the *Drosophila melanogaster* species subgroup. Evol. Biol. 22: 159--225.

Li, H., & Stephan, W. (2006). Inferring the demographic history and rate of adaptive substitution in *Drosophila*. PLOS Genetics, 2(10), e166. https://doi.org/10.1371/journal.pgen.0020166

Lynch, M. (2010). Evolution of the mutation rate. Trends in Genetics, 26(8), 345–352. https://doi.org/10.1016/j.tig.2010.05.003

Lynch, M., Ackerman, M. S., Gout, J.-F., Long, H., Sung, W., Thomas, W. K., & Foster, P. L. (2016). Genetic drift, selection and the evolution of the mutation rate. Nature Reviews Genetics, 17(11), 704–714. https://doi.org/10.1038/nrg.2016.104

Mackay, T. F. C., Richards, S., Stone, E. A., Barbadilla, A., Ayroles, J. F., Zhu, D., Casillas, S., Han, Y., Magwire, M. M., Cridland, J. M., et al. (2012). The *Drosophila melanogaster* genetic reference panel. Nature, 482(7384), 173–178. https://doi.org/10.1038/nature10811

Maruyama, T., & Kimura, M. (1974). A note on the speed of gene frequency changes in reverse directions in a finite population. Evolution, 28, 161–163. https://doi.org/10.1111/j.1558-5646.1974.tb00736.x

Maynard Smith, J., & Haigh, J. (1974). The hitch-hiking effect of a favourable gene. Genetical Research, 23(1), 23–35. https://doi.org/10.1017/S0016672300014634

Morales-Arce, A.Y., Sabin, S.J., Stone, A.C., and Jensen, J.D. (2021). The population genomics of within-host *Mycobacterium tuberculosis*. Heredity, 126(1), 1–9. https://doi.org/10.1038/s41437-020-00377-7

Nielsen, R., Williamson, S., Kim, Y., Hubisz, M. J., Clark, A. G., & Bustamante, C. (2005). Genomic scans for selective sweeps using SNP data. Genome Research, 15(11), 1566–1575. https://doi.org/10.1101/gr.4252305

Orr, H. A., & Betancourt, A. J. (2001). Haldane’s sieve and adaptation from the standing genetic variation. Genetics, 157(2), 875–884. https://doi.org/10.1093/genetics/157.2.875

Pavlidis, P., & Alachiotis, N. (2017). A survey of methods and tools to detect recent and strong positive selection. Journal of Biological Research-Thessaloniki, 24(1), 7. https://doi.org/10.1186/s40709-017-0064-0

Pavlidis, P., Hutter, S., & Stephan, W. (2008). A population genomic approach to map recent positive selection in model species. Molecular Ecology, 185, 907–922. https://doi.org/10.1111/j.1365-294X.2008.03852.x

Pavlidis, P., Jensen, J. D., & Stephan, W. (2010). Searching for footprints of positive selection in whole-genome SNP data from nonequilibrium populations. Genetics, 185(3), 907–922. https://doi.org/10.1534/genetics.110.116459

Payseur, B. A., Cutter, A. D., & Nachman, M. W. (2002). Searching for evidence of positive selection in the human genome using patterns of microsatellite variability. Molecular Biology and Evolution, 19(7), 1143–1153. https://doi.org/10.1093/oxfordjournals.molbev.a004172

Pedregosa, F., Varoquaux, G., Gramfort, A., Michel, V., Thirion, B., Grisel, O., Blondel, M., Müller, A., Nothman, J., Louppe, G., Prettenhofer, P., Weiss, R., Dubourg, V., Vanderplas, J., Passos, A., Cournapeau, D., Brucher, M., Perrot, M., & Duchesnay, É. (2011). Scikit-learn: Machine Learning in Python. Journal of Machine Learning Research, 12, 2825–2830.

Peñalba, J. V., & Wolf, J. B. W. (2020). From molecules to populations: Appreciating and estimating recombination rate variation. Nature Reviews Genetics, 21(8), 476–492. https://doi.org/10.1038/s41576-020-0240-1

Pfeifer, S. P. (2020). Spontaneous mutation rates. In S. Y. W. Ho (Ed.), The Molecular Evolutionary Clock (pp. 35–44). Springer International Publishing. https://doi.org/10.1007/978-3-030-60181-2_3

Poh, Y.-P., Domingues, V. S., Hoekstra, H. E., & Jensen, J. D. (2014). On the prospect of identifying adaptive loci in recently bottlenecked populations. PLOS ONE, 9(11), e110579. https://doi.org/10.1371/journal.pone.0110579

Przeworski, M. (2002). The signature of positive selection at randomly chosen loci. Genetics, 160(3), 1179–1189. https://doi.org/10.1093/genetics/160.3.1179

Przeworski, M. (2003). Estimating the time since the fixation of a beneficial allele. Genetics, 164(4), 1667–1676. https://doi.org/10.1093/genetics/164.4.1667

Rahbari, R., Wuster, A., Lindsay, S. J., Hardwick, R. J., Alexandrov, L. B., Al Turki, S., Dominiczak, A., Morris, A., Porteous, D., Smith, B., et al. (2016). Timing, rates and spectra of human germline mutation. Nature Genetics, 48(2), 126–133. https://doi.org/10.1038/ng.3469

Rockman, M. V., & Kruglyak, L. (2009). Recombinational landscape and population genomics of *Caenorhabditis elegans*. PLOS Genetics, 5(3), e1000419. https://doi.org/10.1371/journal.pgen.1000419

Sabeti, P. C., Schaffner, S. F., Fry, B., Lohmueller, J., Varilly, P., Shamovsky, O., Palma, A., Mikkelsen, T. S., Altshuler, D., & Lander, E. S. (2006). Positive natural selection in the human lineage. Science, 312(5780), 1614–1620. https://doi.org/10.1126/science.1124309

Simonsen, K. L., Churchill, G. A., & Aquadro, C. F. (1995). Properties of statistical tests of neutrality for DNA polymorphism data. Genetics, 141(1), 413–429.

Stapley, J., Feulner, P. G. D., Johnston, S. E., Santure, A. W., & Smadja, C. M. (2017). Variation in recombination frequency and distribution across eukaryotes: Patterns and processes. Philosophical Transactions of the Royal Society B: Biological Sciences, 372(1736), 20160455. https://doi.org/10.1098/rstb.2016.0455

Stephan, W. (1995). Perturbation analysis of a two-locus model with directional selection and recombination. Journal of Mathematical Biology, 34(1), 95–109. https://doi.org/10.1007/BF00180138

Stephan, W. (2019). Selective sweeps. Genetics, 211(1), 5–13. https://doi.org/10.1534/genetics.118.301319

Stephan, W., Wiehe, T. H. E., & Lenz, M. W. (1992). The effect of strongly selected substitutions on neutral polymorphism: Analytical results based on diffusion theory. Theoretical Population Biology, 41(2), 237–254. https://doi.org/10.1016/0040-5809(92)90045-U

Stumpf, M. P. H., & McVean, G. A. T. (2003). Estimating recombination rates from population-genetic data. Nature Reviews Genetics, 4(12), 959–968. https://doi.org/10.1038/nrg1227

Tajima, F. (1989). Statistical method for testing the neutral mutation hypothesis by DNA polymorphism. Genetics, 123(3), 585–595.

Terbot, J.W., Johri, P., Liphardt, S.W., Soni, V., Pfeifer, S.P., Cooper, B.S., Good, J.M., & Jensen, J.D. (2023). Developing an evolutionary baseline model for the study of SARS-CoV-2 patient samples. PLOS Pathogens, 19(4), e1011265. https://doi.org/10.1371/journal.ppat.1011265

Teshima, K. M., Coop, G., & Przeworski, M. (2006). How reliable are empirical genomic scans for selective sweeps? Genome Research, 16(6), 702–712. https://doi.org/10.1101/gr.5105206

Thornton, K. (2003). libsequence: A C++ class library for evolutionary genetic analysis. Bioinformatics, 19(17), 2325–2327. https://doi.org/10.1093/bioinformatics/btg316

Thornton, K., & Andolfatto, P. (2006). Approximate Bayesian inference reveals evidence for a recent, severe bottleneck in a Netherlands population of *Drosophila melanogaster*. Genetics, 172(3), 1607–1619. https://doi.org/10.1534/genetics.105.048223

Thornton, K. R., & Jensen, J. D. (2007). Controlling the false-positive rate in multilocus genome scans for selection. Genetics, 175(2), 737–750. https://doi.org/10.1534/genetics.106.064642

Thornton, K. R., Jensen, J. D., Becquet, C., & Andolfatto, P. (2007). Progress and prospects in mapping recent selection in the genome. Heredity, 98(6), 340–348. https://doi.org/10.1038/sj.hdy.6800967

Wiehe, T. H., & Stephan W. (1993). Analysis of a genetic hitchhiking model, and its application to DNA polymorphism data from Drosophila melanogaster. Molecular Biology and Evolution. https://doi.org/10.1093/oxfordjournals.molbev.a040046

